# Spatiotemporal single-cell profiling of gastrointestinal GVHD reveals invasive and resident memory T cell states

**DOI:** 10.1101/2020.07.20.212399

**Authors:** Victor Tkachev, James Kaminski, E. Lake Potter, Scott N. Furlan, Alison Yu, Daniel J. Hunt, Connor McGuckin, Hengqi Zheng, Lucrezia Colonna, Ulrike Gerdemann, Judith Carlson, Michelle Hoffman, Joe Olvera, Chris English, Audrey Baldessari, Angela Panoskaltsis-Mortari, Benjamin Watkins, Muna Qayed, Yvonne Suessmuth, Kayla Betz, Brandi Bratrude, Amelia Langston, John Horan, Jose Ordovas-Montanes, Alex K. Shalek, Bruce R. Blazar, Mario Roederer, Leslie S. Kean

## Abstract

One of the central challenges in the field of allo-immunity is deciphering the mechanisms driving T cells to infiltrate and subsequently occupy target organs to cause disease. The act of CD8-dominated T cell infiltration is critical to acute graft-versus-host disease (aGVHD), wherein donor T cells become activated, tissue-infiltrating and highly cytotoxic, causing wide-spread tissue damage after allogeneic hematopoietic stem cell transplant (allo-HCT). However, in human and non-human primate studies, deconvolving the transcriptional programs of newly recruited relative to resident memory T cells in the gastrointestinal (GI) tract has remained a challenge. In this study, we combined the novel technique of Serial Intravascular Staining (SIVS) with single-cell RNA-Seq (scRNA-seq) to enable detailed dissection of the tightly connected processes by which T cells first infiltrate tissues and then establish a pathogenic tissue residency program after allo-HCT in non-human primates. Our results have enabled the creation of a spatiotemporal map of the transcriptional drivers of CD8 T cell infiltration into the primary aGVHD target-organ, the GI tract. We identify the large and small intestines as the only two sites demonstrating allo-specific, rather than lymphdepletion-driven T cell infiltration. The donor CD8 T cells that infiltrate the GI tract demonstrate a highly activated, cytotoxic phenotype while simultaneously rapidly developing canonical tissue-resident memory (T_RM_) protein expression and transcriptional signatures, driven by IL-15/IL-21 signaling. Moreover, by combining SIVS and transcriptomic analysis, we have been able to work backwards from this pathogenic T_RM_ programing, and, for the first time, identify a cluster of genes directly associated with tissue invasiveness, prominently including specific chemokines and adhesion molecules and their receptors, as well as a central cytoskeletal transcriptional node. The clinical relevance of this new tissue invasion signature was validated by its ability to discriminate the CD8 T cell transcriptome of patients with GI aGVHD. These results provide new insights into the mechanisms controlling tissue infiltration and pathogenic CD8 T_RM_ transcriptional programing, uncovering critical transitions in allo-immune tissue invasion and destruction.

**One sentence summary:** Flow cytometric and transcriptomic analysis reveals coordinated tissue-infiltration and tissue-residency programs driving gastrointestinal aGVHD.

## INTRODUCTION

T cell infiltration into secondary lymphoid organs (SLOs) and non-lymphoid tissues (NLTs) is central to T cell function, and occurs during homeostatic tissue surveillance (*1, 2*), as well as during T cell-driven immunopathology (including auto- (*3, 4*) and allo-immune disorders (*5, 6*)) and T cell-mediated anti-tumor immune attack (*7, 8*). One of the best-studied clinical instances of T cell infiltration occurs during acute graft-versus-host disease (aGVHD), wherein donor T cells become activated, tissue-infiltrating, and highly cytotoxic after hematopoietic stem cell transplant (HCT) (*9*). During aGVHD, donor T cells first populate SLOs, where they undergo allo-antigen priming, and then home towards and infiltrate non-lymphoid GVHD-target organs (*10–12*). Upon infiltration, these cells induce the immunologic and clinical manifestations of aGVHD, including wide-spread organ damage (*13*). In mouse models, it has been demonstrated that donor T cells acquire tissue resident memory (T_RM_) features (including expression of CD103) during aGVHD (*14–16*). However, central questions remain concerning how these pathogenic T_RM_ are distinguished from protective T_RM_, how donor T_RM_ become activated and cause tissue damage, and perhaps most important, the mechanisms controlling the act of T cell invasion into target organs prior to their evolution into T_RM_. Determining the drivers of this dynamic process represents a critical unmet need, with broad relevance for alloimmunity, as well as for autoimmune diseases and immuno-oncology.

One of the major barriers to understanding the control of T cell infiltration into target tissues has been the inherent difficulty in capturing the key variable of *time* in this process. Time is a critical variable, given that T cells are actively homing towards and infiltrating into target organs, and are concordantly evolving a highly pathogenic transcriptional program. Without understanding the time-dependency of T cell infiltration, we cannot fully comprehend key drivers and their dynamics. Here, we have been able to deconvolute time as a variable during T cell infiltration and tissue destruction during primate aGVHD by applying Serial Intravascular Staining (‘SIVS’) and pairing this with multiparameter flow cytotmetric and single-cell transcriptomic analysis. We have performed these experiments using the non-human primate (NHP) aGVHD model(*17–20*), to best replicate the complex dynamics and molecular drivers of clinical tissue infiltration and disease in an out-bred immunologically experienced transplant recipient. This model takes advantage of the close functional similarity of the immune systems in NHP and humans, such that insights can be most rapidly translated to the clinic. SIVS allowed us to differentially label T cells longitudinally in individual animals during active tissue infiltration, in order to ascertain the time-, tissue- and donor-dependence of T cell-mediated immune pathology during aGVHD. Using SIVS, we demonstrate rapid tissue infiltration followed by differentiation of donor CD8 T cells *in situ* into pathogenic T_RM_, correlating directly with destruction of aGVHD target tissues. These experiments have identified, for the first time, the transcriptional program characterizing actively infiltrating T cells, as well as the transcriptome of pathogenic resident cells that are critical for tissue damage during aGVHD.

## RESULTS

### Applying SIVS to a non-human primate (NHP) aGVHD model

We have built a NHP model of allogeneic HCT (allo-HCT) and aGVHD, in which a donor graft is transplanted into MHC-haploidentical recipients pre-conditioned with myeloablative total body irradiation (*17, 19*). For the experiments described herein, transplant recipients received no post-transplant immunosuppression, which enabled interrogation of the natural history of aGVHD (**Figure 1A**). Two control cohorts, encompassing autologous HCT (auto-HCT) recipients and untransplanted healthy controls (HC), were also analyzed (**Figure 1A**). In accordance with our previous data (*17, 19*), allo-HCT without immunosuppression resulted in early-onset lethal aGVHD, with skin and GI tract-predominant clinical disease (**Figure 1B,C**), and skin, GI-tract and liver histopathology (**Figure 1D**), coinciding with donor T cell engraftment (**Figure 1E**). In contrast, auto-HCT controls survived long-term without aGVHD (**Figure 1C**). aGVHD clinical activity began on ∼Day +5 and peaked on Day +7-8 (**Figure 1B**), similar to the kinetics of mouse alloimmune T cell trafficking to GVHD-target organs (*13, 15*). We performed a series of *in vivo* studies on Day +8 post-transplant, to determine the mechanisms driving tissue infiltration during severe aGVHD.

**Figure 1.**
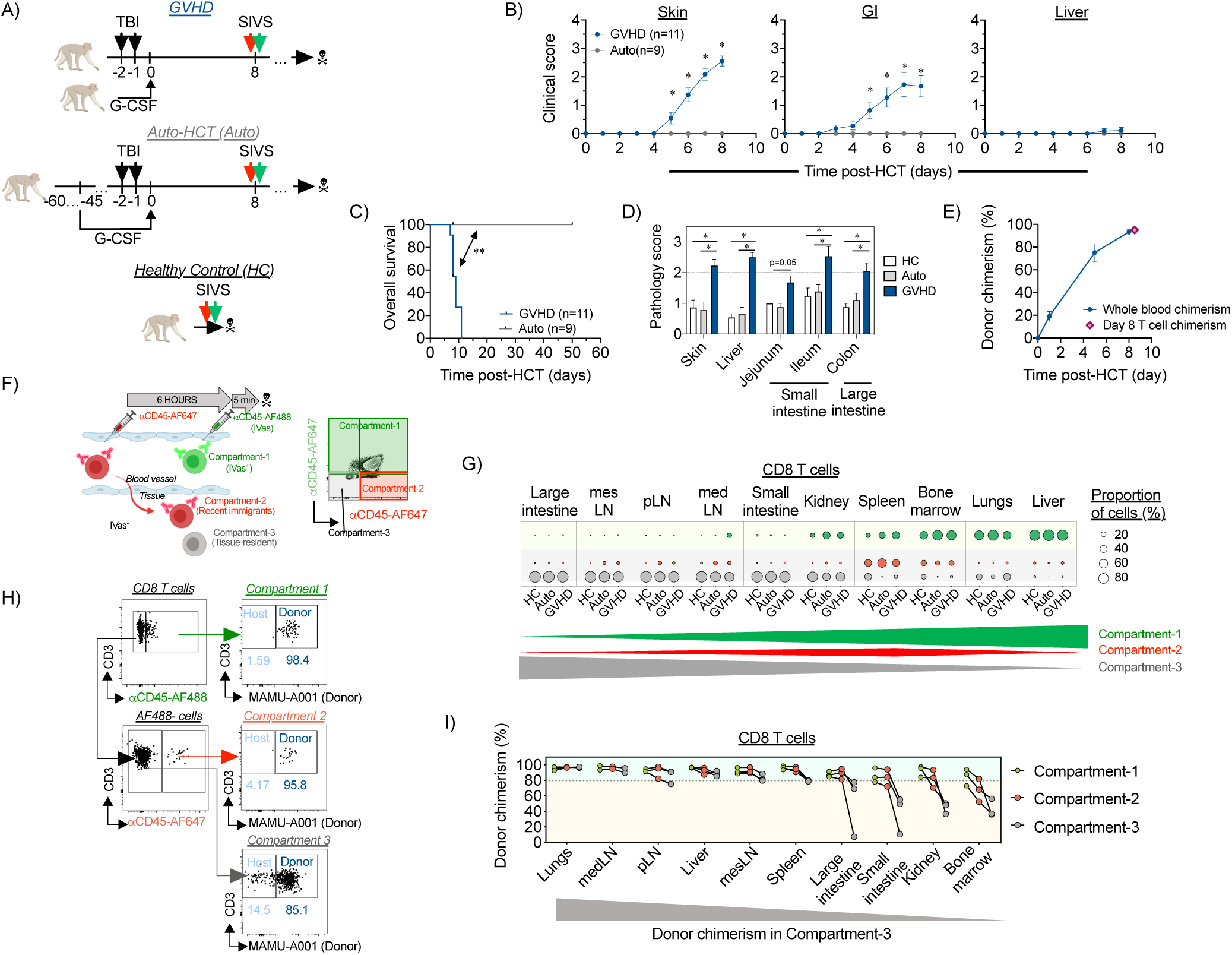
Spatiotemporal compartmentalization of T cells in tissues during aGVHD. A) Experimental schema of Hematopoietic Stem Cell Transplant (HCT) studies in Non-Human Primates (NHP). Abbreviations: GVHD – Graft-versus-Host Disease, auto-HCT – autologous HCT, healthy controls – HC B) Clinical scores for skin (left panel), gastrointestinal tract (middle panel) and liver (right panel) symptoms of acute Graft-versus-Host Disease (aGVHD), determined daily based on published scoring criteria(*19*) in GVHD (n=11) and auto-HCT (n=9) experimental cohorts. *p<0.05, two-way ANOVA with multiple comparison Holm-Sidak post-test. C) Overall Survival of recipients following allo- and auto-HCT without immunosuppression. Studies with pre-set experimental endpoints were censored. **p<0.01 using log-rank (Mantel-Cox) test. D) aGVHD histopathology scores of the indicated organs in healthy control animals or in recipients on Day +8 after allo- or auto-HCT without immunosuppression. *p<0.05, using multiple comparison Holm-Sidak test. E) Donor chimerism in whole blood (*dark blue line*) or FACS-purified peripheral blood T cells (*pink diamond*) of allo-HCT recipients measured by microsatellite analysis. F) Experimental schema and representative flow cytometry plot from healthy control lung, depicting the Serial Intravascular Staining method (SIVS) using fluorescent *α*CD45-antibodies (see **Methods** for details) to discern Intravascular (IVas^+)^ intravascular (Compartment-1 green) and IVas^-^ extravascular compartments: recent infiltrating cells (Compartments-2 red) and resident cells (Compartment-3 ray) and track cellular migration into tissues. G) Relative distribution of CD8 T cells between intravascular (Compartment-1; green dots), the recent infiltrating cell compartment (Compartment-2; red dots) and the resident compartment (Compartment-3; gray dots) in the indicated organs of healthy control animals (HC; n=4), auto-HCT cohort (auto-HCT; n=4) and the allo-HCT cohort (GVHD; n=4) on Day +8 post-transplant. Dot sizes indicate the relative proportion of cells within each compartment, compared to the total number of cells. The green, and gray triangles and the red diamond at the bottom of the figure illustrate the pattern of the three compartments in the tissues examined. pLN – peripheral Lymph Nodes, medLN – mediastinal Lymph Nodes, mesLN – mesenteric Lymph Nodes. H) A representative gating tree for the measurement of donor chimerism in the colon of a recipient from the allo-HCT cohort, based on discordant expression of MHC-I allele MAMU-A001, detected with allomorph-specific antibodies, in the different compartments identified using SIVS. I) Donor CD8 T cell chimerism in Compartments 1-3 (Compartment-1 – green dots, Compartment-2 – red dots, Compartment-3 – gray dots) in different organs on day +8 following MAMU-A001-mismatched MHC-haploidentical allogeneic HCT (n=3). The gray triangle at the bottom of the figure illustrates the pattern of donor chimerism in the tissues examined.

To dissect the spatiotemporal compartmentalization in aGVHD, we used a novel method to study T cell trafficking, known as ‘Serial Intravascular Staining’ (SIVS, described in the companion manuscript (*21*)). SIVS takes advantage of direct intravenous injection of sub-saturating doses of *α*CD45 antibodies, conjugated to different fluorescent tags, which label leukocytes in the systemic circulation at the time of injection, and at different time-points prior to analysis (6 hours and 5 minutes, respectively). This strategy allowed us to differentiate 3 spatiotemporal compartments in multiple tissues (**Figure 1F; Table S1**). The first compartment was identified by injecting a green-fluorescent AlexaFluor488 (AF488)-tagged *α*CD45 antibody 5 minutes prior to euthanasia, which restricted *α*CD45-AF488 labeling to the intravascular compartment (‘IVas^+^’ or ‘Compartment-1’) (**Figure 1F and Figure S1**). Compartments-2 and -3 were identified using a second *α*CD45 antibody, conjugated with the far red-fluorescent AlexaFluor647 (AF647), injected 6 hours prior to *α*CD45-AF488 injection. When far red-fluorescent *α*CD45-AF647 stained cells migrated out of the intravascular space into tissues, they retained their fluorescent tag, and thereby could be distinguished from cells that were in the intravascular space at the time of necropsy. These cells were identified as red (CD45-AF647^+^), but not green (CD45-AF488^-^), and are referred to as ‘Compartment-2’ or ‘recent infiltrating’ cells (**Figure 1F and Figure S1**). These two fluorescently-tagged cell populations could be further distinguished from a third population of cells, which remained negative for both the green and the red CD45 labels. These CD45-AF647^-^CD45-AF488^-^ cells were present in the tissues before the 6-hour CD45-AF647 injection, and were therefore protected from *α*CD45 staining, and are referred to herein as ‘Compartment-3’ or ‘resident’ cells (**Figure 1F and Figure S1**).

Using the SIVS technique, we were able to deconvolute the organ-specific spatiotemporal compartmentalization of both CD4 and CD8 T cells after transplant (gated as shown in **Figure S1**, T cell subset definitions are provided in **Table S2**). Given the central role that CD8 T cells play in aGVHD-mediated organ destruction (*22, 23*), we concentrate our discussion on the trafficking of these cells. Results from CD4 T cells are described in **Supplementary Data**. We identified three different classes of organs/tissues based on their relative balance of Compartment-1, -2 or -3 CD8 T cells (**Figure 1G**). We did not purify T cells from the skin in this study, and so we limit our discussion to visceral organs and tissues. Liver and lungs contained predominantly the intravascular, Compartment-1 T cells. Bone marrow and spleen contained predominantly the recently infiltrating, Compartment-2 cells (as well as a significant proportion of Compartment-1 cells). Lymph nodes, kidneys, small intestine (with T cells isolated from the jejunum) and large intestine contained predominantly resident Compartment-3 CD8 T cells (**Figure 1G**). Similar organ-specific compartmentalization was observed for CD4 T cells (**Figure S2A**). Notably, these distributions did not significantly change between the three experimental conditions (HC, auto-HCT and aGVHD) suggesting that spatiotemporal T cell compartmentalization may be a fundamental characteristic of organ immune structure that is not impacted by either homeostatic or allo-activated T cell trafficking on the time scales studied.

### Combining SIVS and the measurement of Donor Chimerism to Identify T cells In the Act of Infiltrating GVHD Target Organs

While the Compartment-3-predominant organs (lymph nodes, kidney, and GI tract) all appeared similar through SIVS, by specifically tracking donor versus host T cell identity during SIVS, we discovered a critical distinction between these tissues after allo-HCT. This distinction was based on the balance between donor and host T cells in these tissues (**Figure 1H**), reflecting the pace of replacement of recipient T cells by donor cells after allo-HCT. These studies revealed that *donor-origin* Compartment-3 CD8 and CD4 T cells predominated in the lymph nodes on Day +8 (**Figure 1I** and **Figure S2B),** consistent with rapid entry of transplanted T cells and replacement of host cells early after transplant. In contrast, a substantial fraction of *host-origin* T cells remained in the tissue-resident Compartment-3 cells of the large intestine, small intestine, kidney, and bone marrow (**Figure 1I and Figure S2B**) during the period of active infiltration of donor T cells into these tissues. These data suggest a tenacious resident T cell niche in these tissues, in which host T cells persist after transplant, even surviving lethal irradiation (*24*) and further link the timing of donor T cell infiltration into these tissues with the onset of clinical disease. Importantly, while a high proportion of host CD8 T cells remained in Compartment-3 in the intestines, kidney and bone marrow, there were essentially no host T cells in the actively-infiltrating Compartment-2 cells in these same tissues (**Figure 1I** and **Figure S2B**). This dichotomy suggests that our analysis has been able, for the first time, to catch donor T cells *in the act of organ infiltration*. This is particularly relevant for donor T cells infiltrating the intestines, since this infiltration was associated with lethal GI aGVHD (**Figure 1B,C**). This provided a strategy to interrogate the protein and gene expression patterns that mark these cells, and control their transit into aGVHD target tissues.

To rigorously distinguish pathogenic T cell infiltration during aGVHD from the physiologic T cell trafficking that occurs during homeostatic reconstitution, we employed a critical control: applying SIVS after autologous HCT. While our studies in allo-HCT identified multiple organs undergoing T cell infiltration after transplant, homeostatic expansion occurs along side allo-activated T cell expansion during allo-HCT, given the exposure of recipients to pre-transplant lymphodepleting irradiation. Comparing the extent of tissue infiltration by Compartment-2 cells in allo-HCT versus auto-HCT controls (both analyzed on Day +8 post-transplant) allowed us to deconvolute the role that homeostatic reconstitution played in T cell tissue infiltration. To complete this analysis, the extent of tissue migration was determined by calculating the percentage of tissue-infiltrating Compartment-2 T cells compared to total extravascular (IVas^-^, Compartment-2 + Compartment-3) cells (representative flow cytometry from the colon shown in **Figure 2A**). This analysis demonstrated that for many tissues (including peripheral and visceral lymph nodes, spleen, bone marrow, liver, lungs and kidneys), the amount of post-transplant CD8 (**Figure 2B-D**) and CD4 (**Figure 2B-D**, **Figure S3A-E**) T cell migration was not different after allo-HCT compared to post-auto-HCT conditions, suggesting that most of the T cell trafficking was in response to lymphodepletion, which abolished competition for a niche in these organs. Indeed, when corrected using the auto-HCT controls, only two sites exhibited statistically significant differential CD8 T cell infiltration during aGVHD: the large and small intestines (**Figure 2D**), and only the large intestine was identified as a site of GVHD-specific CD4 T cell infiltration (**Figure S3E**). This result is especially notable given that the GI tract has been repeatedly demonstrated in the NHP aGVHD model as key site of clinical and histopathologic disease (*17, 19*).

**Figure 2.**
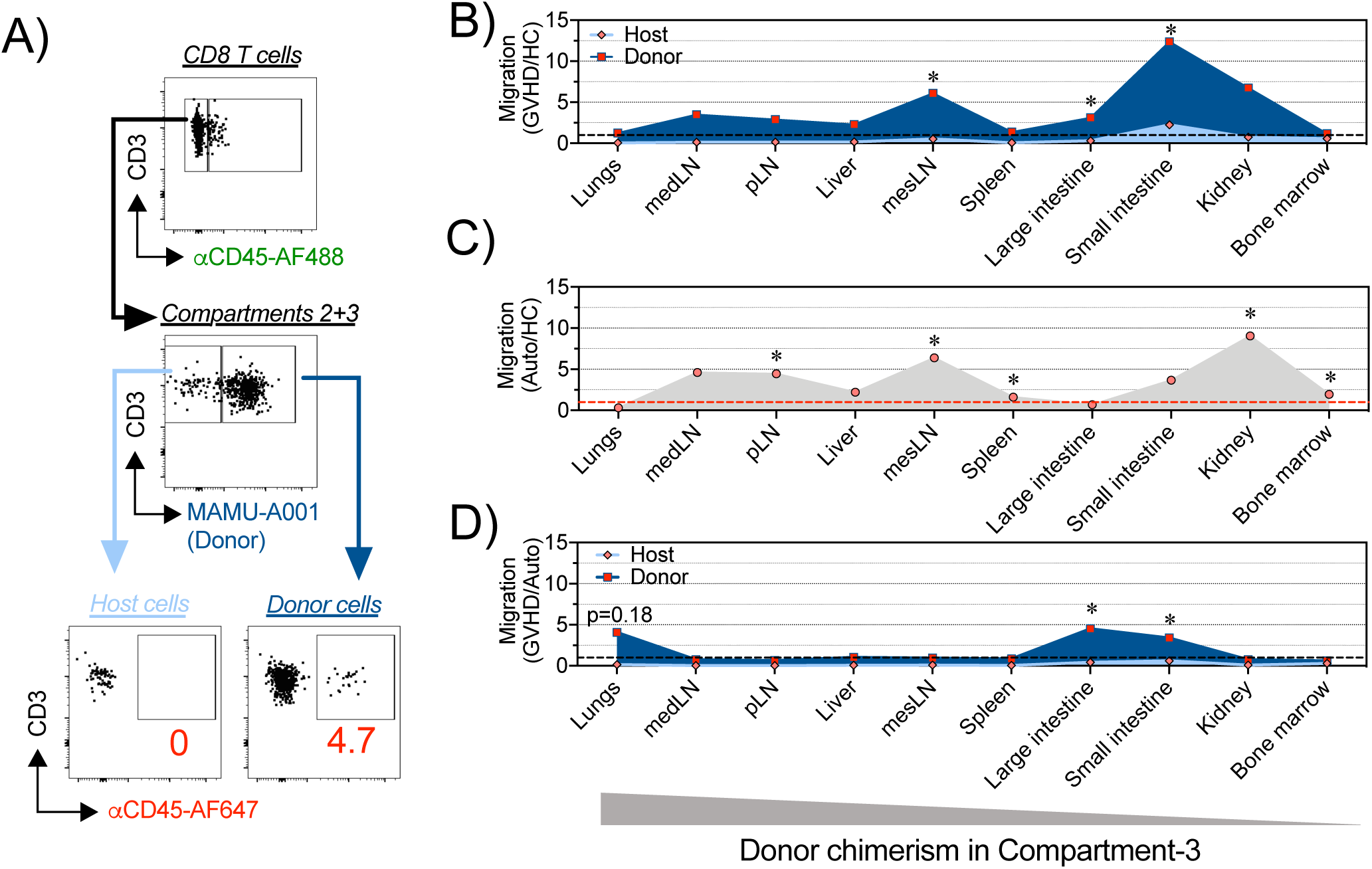
Dynamics of T cell trafficking into GVHD-target organs following HCT in NHP. A) A representative gating tree for flow cytometric SIVS-based measurement of donor and host cell migration in MAMU-A001-mismatched transplantation from the allo-HCT cohort on day +8 after HCT. B) Relative migration of donor and host CD8 T cells in different organs and tissues on day +8 following MAMU-A001-mismatched allo-HCT (n=3) as a percent of recent infiltrating (Compartment-2 cells) out of all extravascular CD8 T cells (sum of Compartments-2 + Compartment-3), normalized to the migration of CD8 T cells in untransplanted healthy controls (n=4). C) Relative migration of CD8 T cells on day +8 from the auto-HCT cohort (n=4) normalized to the migration of CD8 T cells in untransplanted healthy controls (n=4). D) Relative migration of donor and host CD8 T cells in different organs and tissues on day +8 following MAMU-A001-mismatched allo-HCT (n=3), normalized to the migration of CD8 T cells in untransplanted healthy controls (n=4). For B to D: pLN – peripheral Lymph Nodes, medLN – mediastinal Lymph Nodes, mesLN – mesenteric Lymph Nodes. *p<0.05 using t-test with Holm-Sidak multiple comparison correction.

### Defining characteristics of actively–infiltrating Compartment-2 Cells

The major impact that Day +8 GI T cell infiltration had on clinical aGVHD was further underscored by the positive correlation between the extent of CD8 T cell migration into the large and (to a lesser extent) small intestine and the severity of aGVHD pathology in individual transplant recipients – a correlation that was not observed in any other organ (**Figure 3A**). We therefore determined the prominent characteristics of donor cells infiltrating the GI tract, in order to understand the drivers of lethal aGVHD. As shown in **Figures 3B-H**, flow cytometry identified a number of unique attributes of intestinal tissue-infiltrating Compartment-2 CD8 T cells. This included the observation that, while they were rigorously demonstrated to be extra-vascular and tissue-infiltrating by their lack of staining by the green-fluorescent iVas+ CD45 antibody, these intestinal Compartment-2 CD8 T cells did not yet express the canonical T_RM_ CD69^+^CD103^+^ phenotype (**Figure 3B**), consistent with their migratory phenotype. Their rapid evolution towards a T_RM_ phenotype *in situ* was, however, evident, as donor CD8 T cells acquired the canonical T_RM_ CD69^+^CD103^+^ phenotype during their transition from Compartment-2 to Compartment-3, with the highest proportion of CD69^+^CD103^+^ expression in residual host CD8 intestinal T cells (**Figure 3B, Figure S4A,B**). These data suggest that during intestinal aGVHD, donor-derived CD8 T_RM_ cells differentiate *in situ* from migratory precursors, likely in response to microenvironmental instructive signals (as has been demonstrated in other models (*1, 25–27*), including intestinal transplantation (*6, 28*)). Taken together, these results document the rapid acquisition of a T_RM_ phenotype (within 8 days after transplant) as donor CD8 T cells infiltrate and then occupy the GI tract.

**Figure 3.**
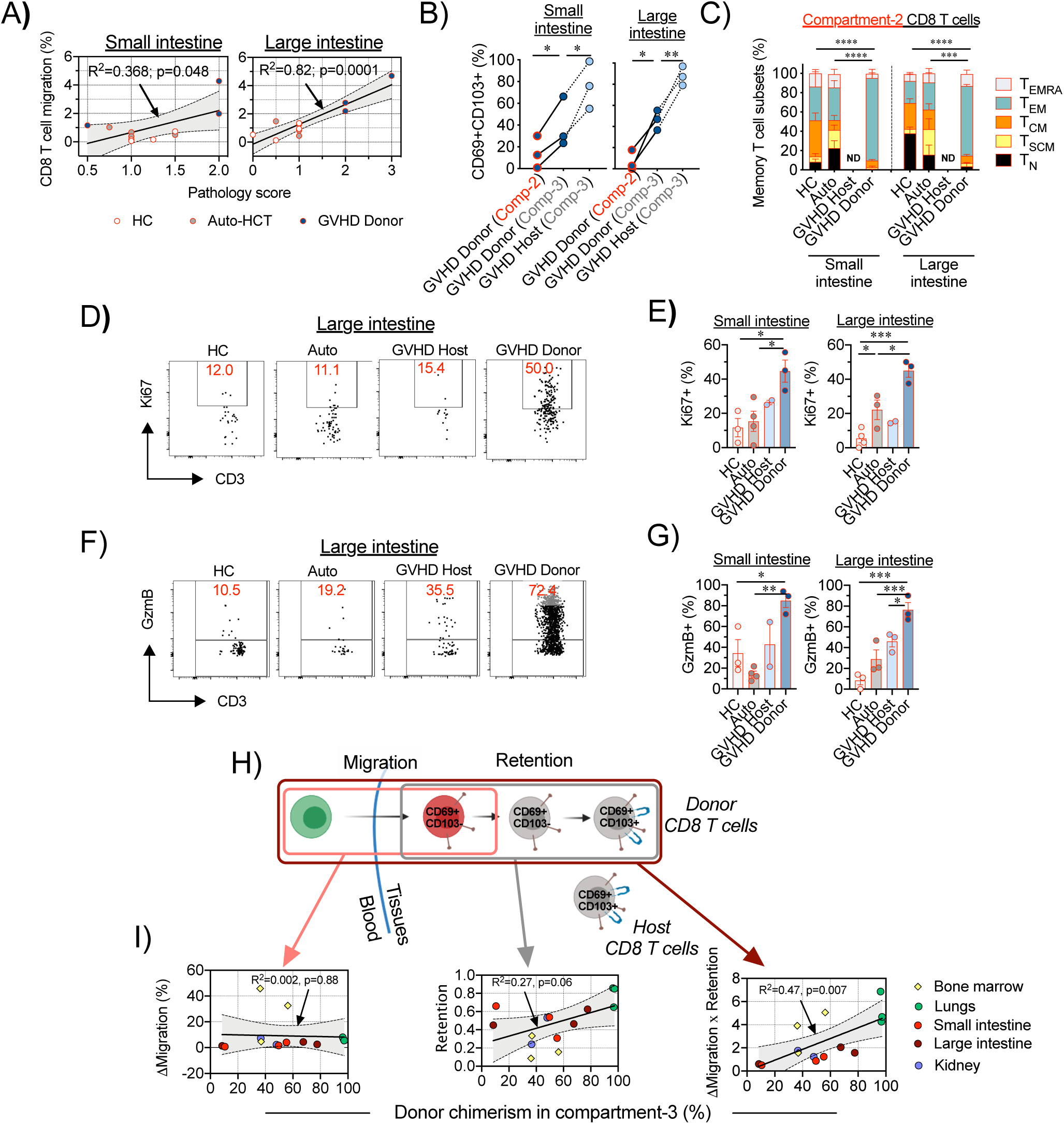
Migration of donor CD8 T cells to the large and small intestine correlates with the severity of tissue pathology during aGVHD. A) Correlation between the migration of CD8 T cells, as a percent of recent infiltrating cells (Compartment-2 cells) out of all extravascular CD8 T cells (sum of Compartments-2 + Compartment-3), and GVHD histopathology scores in the small intestine (jejunum; *left panels*) and large intestine (colon; *right panels*). Lines with tinted areas indicate the 95% CI using linear regression. B) Percentage of donor or host CD8 T cells with the CD69^+^CD103^+^ phenotype within the recent-infiltrating compartment (Compartment-2) and the resident compartment (Comparment-3). Lines connect corresponding values obtained from the same animal. *p<0.05, **p<0.01 using paired t-test. Comp-2 = Compartment-2; Comp-3= Compartment-3 C) Distribution between memory subsets, defined by CCR7, CD45RA and CD95 expression, as detailed in **Methods**, of recent infiltrating CD8 T cells (Compartment-2 CD8 T cells) in the small and large intestines in healthy controls (HC, n=3), auto-HCT cohort on Day +8 (Auto, n=4) and host (GVHD Host) and donor (GVHD Donor) cells following MAMU-A001-mismatched allo-HCT on Day +8 (n=3). Nd – no data. D-E) Representative flow cytometry plots (D) and summary data for Ki67 expression (E) in recent infiltrating (Compartment-2) CD8 T cells in the small and large intestine in healthy controls (HC, n=3), auto-HCT recipients on Day +8 (Auto, n=4) and host and donor cells of MAMU-A001-mismatched allo-HCT on Day +8 (n=3). F-G) Representative flow cytometry plots (F) and the summary data for Granzyme B (GzmB) expression (G) in recent infiltrating (Compartment-2) CD8 T cells in the small and large intestine in healthy controls (HC, n=3), auto-HCT recipients on Day +8 (Auto, n=4) and host and donor cells of MAMU-A001-mismatched allo-HCT on Day +8 (n=3). For E and G: *p<0.05, **p<0.01, ***p<0.001 using one-way ANOVA with Holm-Sidak multiple comparison post-test. H) Schematic representation of donor T cell immigration into GVHD-target NLTs and gradual acquisition of the CD69^+^CD103^+^ T_RM_ phenotype. I) Correlations between donor chimerism in resident CD8 T cells (Compartment-3) on Day +8 following allogeneic MAMU-A001-mismatched HCT (n=3) and the difference between the extent of donor and host CD8 T cell migration (**ΔMigration**, defined as the percentage of recent infiltrating Compartment-2 – donor or host CD8 T cells out of total extravascular – Compartment-2 plus Compartment-3 donor or host CD8 T cells; *left panel*), extent of acquisition of the CD69^+^CD103^+^ T_RM_ phenotype in the immigrated donor CD8 T cells (**Retention**, defined as the ratio of the percentage of donor CD8 T cells expressing the CD69^+^CD103^+^ phenotype to the percentage of host CD8 T cells expressing CD69^+^CD103^+^ phenotype; *middle panel*) and the difference between the extent of donor and host CD8 T cell migration, adjusted to the extent of acquisition of the CD69+CD103+ T_RM_ phenotype (**Migration X Retention**), as detailed in **Methods**, across different non-lymphoid organs and tissues. The line with the tinted area indicates the 95% CI using linear regression.

### Pathogenic hallmarks of actively infiltrating and T_RM_ donor CD8 T cells

Flow cytometric analysis demonstrated that a large proportion of intestinal tissue-infiltrating Compartment-2 CD8 T cells were skewed towards a CCR7^-^CD45RA^-^ effector-memory (T_EM_) phenotype compared to both HC and auto-HCT controls, while other CD8 T cell subpopulations (including naïve, memory stem cells, central memory and effector memory-RA (subpopulation gating in **Table S2** and **Figure S1**) were not overrepresented in Compartment-2 (**Figure 3C**). These intestinal Compartment-2 CD8 T_EM_ cells were highly proliferative and cytotoxic during aGVHD, as measured by their increased expression of Ki67 (**Figures 3D-E**) and Granzyme B (**Figures 3F-G**) compared to untransplanted and auto-HCT controls, or to host Compartment-2 cells, consistent with the observed clinical pathogenicity that accompanied donor T cell tissue infiltration. Notably, in the tissue-resident Compartment-3, CD8 T cells from healthy and auto-HCT controls, and the pathogenic donor T cells during aGVHD, also exhibited a prominent T_EM_ phenotype (**Figure S4C**); however, the Compartment-3 donor cells retained high expression of Ki67 and Granzyme B, documenting their distinction from homeostatic Compompartment-3 cells and consistent with their pathogenicity (**Figure S4D-E**).

### Tissue T cell chimerism is dependent on acquisition of T_RM_ characteristics

It has previously been demonstrated that the expression of CD103 is critical for donor CD8 T cell pathogenicity in the gut, as it enables their retention in proximity to the epithelial layer, where these cells mediate their cytotoxic functions (*14, 16, 29*). However, whether the acquisition of T_RM_ characteristics is also a foundational requirement for donor T cells to anchor in recipient tissues has previously not been a tractable question. By combining SIVS with donor chimerism measurements, we were able to directly address this issue. Our data demonstrate that the acquisition of the CD69^+^CD103^+^ phenotype was critical for establishing long-term residence in non-lymphoid tissues, as both the extent of donor CD8 T cell migration into tissues and *their rate of conversion into T_RM_ cells* (calculations described in detail in **Methods**) together, but not separately, defined tissue donor CD8 T cell chimerism (**Figure 3H-I**).

### Determining transcriptomic features of pathogenic T_RM_ and infiltrating donor CD8 T cells during aGVHD

SIVS allowed us to rigorously identify tissue-associated CD8 T cells in healthy controls and after transplant. We next combined SIVS with single-cell RNA-Sequencing (scRNA-seq) to identify the transcriptional programs associated with the infiltration and pathogenicity of donor-derived tissue-associated CD8 T cells during aGVHD. To do this, we flow cytometrically sorted viable, tissue-associated CD45+ cells from the large intestine of healthy controls (n = 4) and from the aGVHD cohort (n = 4) by gating on non-iVas (non-Compartment-1) cells (**Table S3**). Donor and host cells were flow cytometrically purified separately based on disparate expression of MAMU-A001, where possible (in two of the four aGVHD animals analyzed, **Table S1**). In the two remaining allo-HCT recipients, we used computational methods to discern donor and host cells based on sex-mismatch between the transplant pairs, using Y-chromosome-derived transcripts (see *Methods* for details). Using this methodology, we successfully reconstructed transcriptomes from 21,490 single cells, followed by the identification of 14,185 T cells, based on their expression of canonical T cell genes (**Figure S5**). To identify 4,715 CD8 T cells, we used the annotation tool VISION (*30*) to score each cell using a CD8 vs CD4 signature (*31*), and retained high-scoring clusters.

To systematically determine the features that differentiated pathogenic donor CD8 T_RM_ in the GI tract from physiologic host or HC CD8 T_RM_, we used VISION to score a T_RM_-signature from Milner et al., 2017 (*32*) to all donor, host, and HC CD8 T cells, thereby identifying the continuum of T_RM_ transcriptional programing at a single-cell level in all GI-associated CD8 T cells (**Figure 4A-C).** We chose the Milner T_RM_ signature since it identifies the key genetic regulators of T_RM_ differentiation from multiple tissues regardless of a cell’s activation status, thus providing the most accurate measure of T_RM_ transcriptional features. Notably, the Milner signature was also highly correlated with a tumor-resident T cell signature from patients with colorectal cancer (*33*) (**Figure S6A-B**). As shown in **Figure 4A-C**, and in agreement with flow cytometric analysis (**Figure 3B**), donor-derived GI CD8 T cells had a lower mean T_RM_ score compared to either host- or HC-derived cells (p<0.001 using Wilcoxon test), consistent with there being a higher proportion of actively infiltrating (rather than established T_RM_) donor cells during GI aGVHD. To further determine what differentiated pathogenic donor CD8 T cells from both host and HC CD8 T_RM_, we performed the following analysis: we first defined a T_RM_-high cutoff score by identifying the upper T_RM_ score quartile from HC CD8 T cells (**Figure 4C**) and then applied this score cutoff to both the donor and host CD8 T cells, to identify T_RM_-high subpopulations in each. This allowed us to probe the transcriptional differentiators between pathogenic donor T_RM_-high cells and both host and HC T_RM_ controls through differential expression (DE) calculations, followed by pathway enrichment analysis and classification of up-stream regulators. This pipeline distinguished the following features of pathogenic CD8 T_RM_ cells: they develop rapidly (within 8 days) after allo-HCT, and consist of pathogenic donor cells that simultaneously enact T_RM_ transcriptional programming in addition to upregulation of key T cell activation programs. These include upregulation of specific cellular metabolic pathways (including oxidative phosphorylation and mitochondrial respiration, which have been demonstrated to be central to both T_RM_ development (*34–36*) and aGVHD pathogenesis (*37, 38*)), co-stimulatory signaling pathways (including aGVHD-promoting ICOS- (*39, 40*), IL-2- (*41, 42*) and OX40- (*20, 43*) signaling pathways) and cytotoxicity pathways (*44*)) compared to their physiologic counterparts (**Figure 4D-E, Tables S4-S5**). We also identified a significantly higher enrichment score for the aGVHD pathogenicity signature adopted from *Ichiba et al* (*45*) in donor T_RM_-high cells in comparison with host and HC T_RM_-high cells (**Figure 4F-G**), further substantiating the pathogenicity of the donor T_RM_. Of note, our analysis of the up-stream regulators that control donor T_RM_-high activation pathways revealed involvement of IL-15 and IL-21 cytokine signaling (**Figure 4H, Table S6**), suggestive of a central role for these cytokines in T_RM_ differentiation (particularly during lymphopenic conditions) (*46, 47*), as well as their pathogenic role during aGVHD (*48–52*). We further confirmed that both IL-15 and IL-21 could induce maturation towards a CD69^+^CD103^+^ T_RM_ phenotype in NHP splenocytes *in vitro*, with IL-15 inducing T_RM_ differentiation in splenocytes isolated from either HC or from animals with aGVHD, and IL-21 promoting T_RM_ differentiation only in aGVHD-derived cells (**Figure 4I**). These data thus identify multiple transcriptional pathways and up-stream regulators that are strongly associated with aGVHD-mediated tissue destruction, providing the first direct link between T_RM_ differentiation and donor T cell pathogenesis in primate aGVHD.

**Figure 4.**
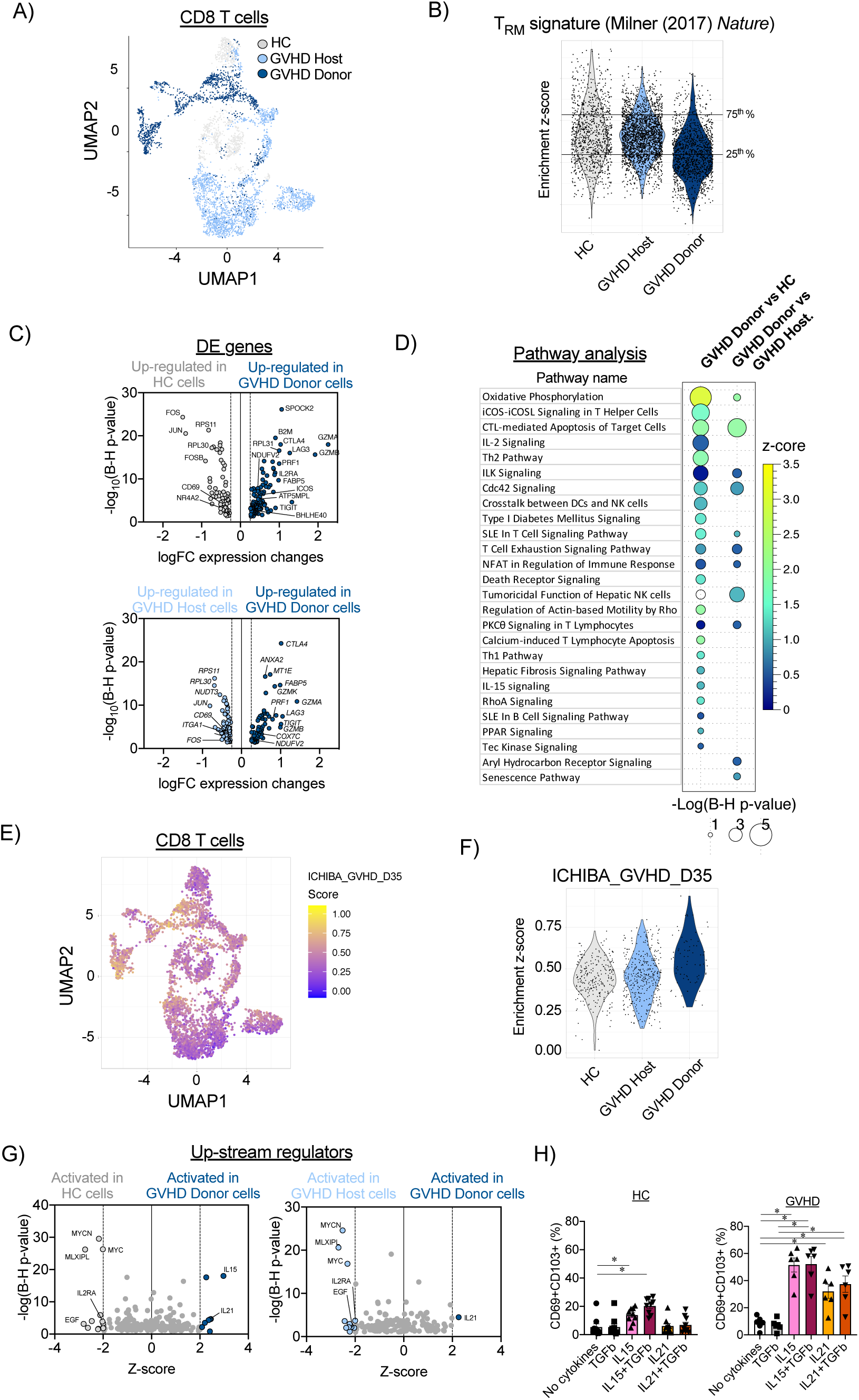
Transcriptomic profile of donor-derived CD8 T_RM_ cells in the large intestine during aGVHD. A) CD8 T cell census (n=4,715) in Uniform Manifold Approximation and Projection (UMAP) space. Cells are colored by experimental cohort. B) Enrichment scores for the T_RM_-signature (adopted from *Milner et al, Nature, (2017)*(*32*)) of healthy controls (HC) host and donor CD8 T cells. Horizontal lines represent the 25^th^ and 75^th^ percentiles for enrichment scores of the healthy control cohort, which were used as cut off values to determine T_RM_-low and T_RM_-high cells in donor and host samples, as defined in **Methods**. C) Differentially expressed genes, identified using the Wilcoxon test, between donor CD8 T cells with high enrichment scores for the T_RM_ signature (>/= 75^th^ percentile as defined for the HC cohort, as detailed in **Methods**), and their counterparts from healthy controls (HC; *top panel*) and host cells (*bottom panel*). D) Pathway analysis performed using Ingenuity Pathway Analysis (IPA, see **Methods**) on differentially expressed genes between donor CD8 T cells with high enrichment scores for the T_RM_ signature (>/= 75^th^ percentile as defined for the HC cohort, as detailed in **Methods**), and their counterparts from healthy controls and host cells, depicting signaling pathways with positive enrichment scores and p<0.05 using a t-test with the Benjamini-Hochberg correction. E) CD8 T cell census (n=4,715), clustered in Uniform Manifold Approximation and Projection (UMAP) space. Cells are colored by the enrichment score for the GVHD signature adopted from *Ichiba et al, Blood, (2003)* (*45*). F) Enrichment scores for the aGVHD signature (adopted from *Ichiba et al* (*45*)) of healthy controls (HC), host, and donor CD8 T_RM_-high cells. G) Predicted up-stream regulators based on Ingenuity Pathway Analysis (IPA, see **Methods**) performed on DE genes between donor CD8 T cells with high enrichment scores for the T_RM_ signature (>/= 75^th^ percentile as defined for the HC cohort, as detailed in **Methods**) and their counterparts from healthy controls (*left panel*) and host CD8 T cells (*right panel*). H) Percentages of spleen CD8 T cells, isolated from healthy control (HC) animals or animals with aGVHD on day +8 after allo-HCT, expressing the CD69^+^CD103^+^ T_RM_ phenotype 48 hours after incubation with the indicated cytokines. *p<0.05, using one-way paired ANOVA with Geisser-Greenhouse correction and Holm-Sidak multiple comparison post-test.

### Identifying the transcriptional control of actively infiltrating donor CD8 T cells

In addition to identifying the drivers of pathogenic CD8 T_RM_, this study also provided an opportunity to interrogate the transcriptional hallmarks of *actively tissue-infiltrating* pathogenic CD8 T cells at an unprecedented level of specificity. To accomplish this, based on the SIVS results (**Figure 3B**) which demonstrated that Compartment-2 actively infiltrating cells display a migratory, rather than T_RM_-like, phenotype, we first identified a T_RM_-low cut-off score (by identifying the lower Milner T_RM_ score quartile from HC CD8 T cells, **Figure 4C**), and then applied this score cut-off to both donor and host CD8 T cells. This allowed us to isolate donor cells that were tissue associated, given that they were first sorted as non-intravascular (Compartment-1-negative) prior to scRNAseq, and simultaneously did not yet display features of T_RM_. We then defined the transcriptional differentiators between pathogenic donor T_RM_-low cells and both host and HC T_RM_-low controls by applying differential expression (DE) calculations (**Figure 5A, Tables S7-S9**). Finally, we focused on the T_RM_-low-specific transcriptome by identifying those DE genes that were unique to the T_RM_-low donor versus control comparisons (and not identified in the T_RM_- high versus control comparisons). This analysis distinguished 53 transcripts (46 up-regulated genes and 7 down-regulated genes) that were specifically dysregulated in donor CD8 T_RM_-low cells, as well as their associated pathways and up-stream regulators (**Figure 5B-D**, **Table S10-S12).** Regulatory network analysis again identified IL-15, as well as type I and type II Interferons (and their corresponding signals) as key up-stream regulators controlling the transcriptional programming of these donor tissue-infiltrating T cells (**Figure 5D**, **Table S12**). Most pertinent to the goal of defining the drivers of T cell infiltration, we identified a subset of genes that have been shown to be critical for cellular adhesion, migration and tissue infiltration in other clinical scenarios. These included chemokines, as well as regulators of their secretion and receptors (*CCL3* (*53*), *CCL4L1* (*54*), *GZMM* (*55*), *CD74* (*56*)), surface adhesion receptors and regulators of their expression (*LTB* (*57*) and *ITGB2* (*58*)); and cytoskeleton components and regulators (*ACTB* (*59*), *ACTG1* (*59*), *CAP1* (*60*), *COTL1* (*61*), *ISG15* (*62*), *LSP1* (*63*), *PFN1* (*64*)) (**Figure 5E).** These data thus provide the first transcriptional map of this unique subpopulation of actively infiltrating donor CD8 T cells, nominating a new class of molecules that enable pathologic CD8 T cell tissue invasion during GVHD.

**Figure 5.**
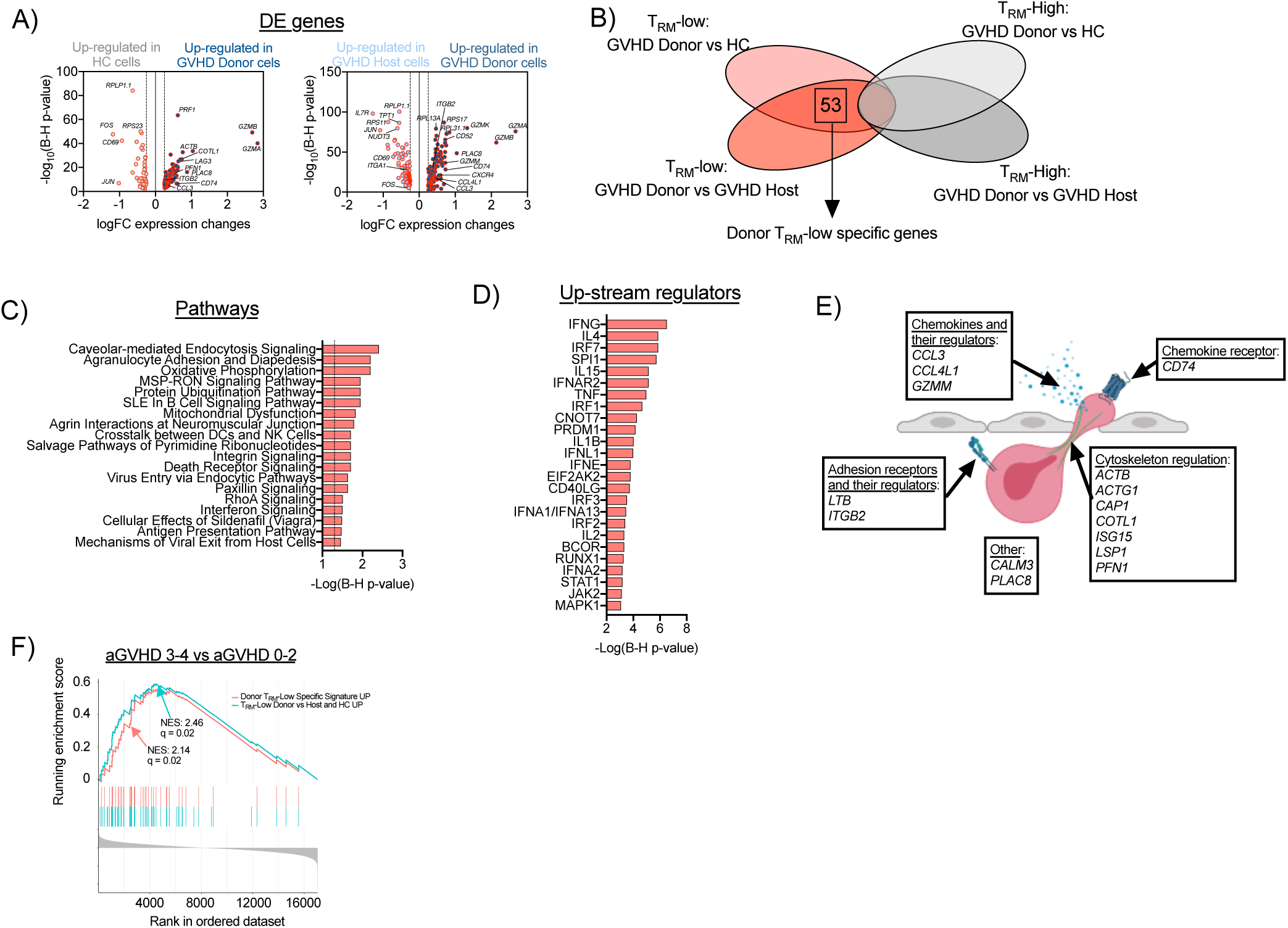
Transcriptomic profile of donor tissue-infiltrating CD8 T cells in the large intestine during aGVHD. A) Differentially expressed genes, identified using the Wilcoxon test (±logFC>0.25; adj. p <0.05), between donor CD8 T cells with low enrichment scores for the T_RM_ signature (</= 25^th^ percentile as defined for the HC cohort, as detailed in **Methods)**, and their counterparts from healthy controls (HC; *left panel*) and host cells (*right panel*). B) Venn diagram illustrating the identification of the donor T_RM_-low-specific gene signature. C-D) Pathway analysis performed using Ingenuity Pathway Analysis (IPA, see **Methods**) on differentially expressed genes between donor CD8 T cells with low enrichment score for the T_RM_ signature and both healthy control and host cells with low enrichment scores for the T_RM_ signature, depicting signaling pathways with p<0.05 using a t-test with the Benjamini-Hochberg correction (C) and the predicted top 25 up-stream regulators (D). E) Functional annotation of migration-related genes differentially expressed between donor CD8 T cells with low enrichment scores for the T_RM_ signature and both healthy control and host cells with low enrichment scores for the T_RM_ signature. F) GSEA plot depicting the enrichment of two T_RM_-low gene signatures compared to a clinical aGVHD CD8 T cell transcriptional signature. The curve labeled “T_RM_-Low Donor vs Host and HC UP” encompasses genes over-represented in donor T_RM_-low CD8 T cells in comparison with both host and HC T_RM_-low counterparts – blue line). The curve labled “Donor T_RM_-Low **specific** signature UP” encompasses genes over-represented **exclusively** in the donor T_RM_-low CD8 T cell comparison (i.e. not also present in the comparison of donor T_RM_-high CD8 T cells versus both host and HC T_RM_-high counterparts) – red line). The gene lists for the “T_RM_-Low Donor vs Host and HC UP” and the ““Donor T_RM_-Low **specific** signature UP” comparisons are found in **Table S15**. Each of these gene lists was used to compare patients with Grade 3-4 aGVHD and patients with Grade 0-2 aGVHD through GSEA. NES – normalized enrichment score.

### Validating the clinical relevance of the NHP tissue-infiltration signature

To validate that the genes and pathways identified through the NHP experiments were relevant to human disease, we compared our NHP transcriptional signatures to a CD8 T cell signature generated from patients with aGVHD. The human aGVHD signature was constructed from peripheral blood CD8 T cells that were sorted on Days +21 and +28 post-transplant from 42 patients receiving an unrelated-donor HCT and standard calcineurin inhibitor/methotrexate aGVHD prophylaxis (**Table S13**). Twenty-eight of these patients developed Grade 2-4 aGVHD (with 27 of 28 having either upper or lower GI disease). All patients with Grade 3-4 aGVHD demonstrated lower GI disease (**Table S13**). The resulting transcriptomes were dichotomized between patients who developed severe Grades 3-4 aGVHD versus those with Grades 0-2 aGVHD (**Figure 5F**) or those with moderate-severe Grades 2-4 aGVHD versus those with Grades 0-1 aGVHD (**Figure S6**). These analyses demonstrated significant overlap between the migratory donor CD8 T_RM_- low signatures and the human CD8 T cell aGVHD signature (**Figure 5F, Figure S6, Tables S14-S15**). These results underscore two important points: first, they provide strong evidence for the relevance of the NHP data to human aGVHD; and second, they demonstrate the ability to probe the peripheral blood for signatures relevant to the migrating pathogenic cells that ultimately infiltrate the GI tract during clinical aGVHD. Thus, while the peripheral blood did not contain the complete pathologic signature derived from GI-tract infiltrating T cells (**Table S14)**, the peripheral blood signature complemented the organ-specific results, which successfully dichotomized patients with significant clinical disease.

## Discussion

One of the central challenges in the field of T cell pathogenesis is the inherent difficulty in directly interrogating the tissue-infiltrating and tissue-resident T cells that are increasingly recognized as essential to immune surveillance and immune-mediated disease. Two issues predominate. First, it is often problematic, especially in large animal models and in patients, to rigorously eliminate intravascular T cells from analysis. This can be a major limitation, as it lowers the specificity of organ-based studies, since these intravascular T cells can contaminate tissue-derived cell preparations. The second issue is the difficulty in independently studying the three distinct cellular components that comprise the intravascular versus tissue-based immune response. These components consist of (1) the intravascular cells that can often contaminate preparations of ostensibly tissue-associated cells; (2) the rare *actively infiltrating cells,* which may be particularly important to the initiation and potentiation of a nascent immune response, and (3) the *established T_RM_ cells* that enact ongoing tissue-based immunity and immunopathology. In this study, we have combined SIVS with scRNA-seq to interrogate both of these critical populations, in order to dissect the mechanisms by which donor CD8 T cells infiltrate and then cause tissue damage during aGVHD.

Here, the development of SIVS and its application to a non-human primate aGVHD model has enabled the interrogation of tissue-based T cell immunity at an unprecedented level of detail in an outbred, pathogen-exposed, immunologically-experienced system. The concept of SIVS is based on the intravascular staining methodology established by the Masopust laboratory (*65*), which enables rigorous exclusion of intravascular cells during preparation and analysis of T_RM_. In this study, by applying *serial* infusions of differentially-labeled anti-CD45 antibodies (‘SIVS’, developed by the Roederer laboratory and described in the companion manuscript (*21*)), we have been able to include time as a variable in this analysis, facilitating the detection of three spatiotemporally-defined T cell compartments: (1) T cells that were intravascular at the time of tissue analysis (‘intravascular’, or ‘Compartment-1’ T cells); (2) T cells that were intravascularly located six hours prior to analysis and had subsequently infiltrated into target tissues (‘recent infiltrating’ or Compartment-2’ T cells); and, (3) T cells that were tissue-associated for at least six hours prior to analysis and did not acquire either of the two CD45 labels (‘resident’ or ‘Compartment-3’ T cells) (**Figure 1**). By overlaying our ability to distinguish cells of donor and host origin after allo-HCT onto these three compartments, we have developed a highly sensitive and specific paradigm for the identification of pathogenic donor CD8 T cells, either in the act of organ infiltration or in the act of tissue destruction during severe GI aGVHD.

Our observations provide strong support for the rapid evolution of a pathogenic T_RM_ transcriptional program in donor CD8 T cells after allo-HCT. This observation represents a striking counterpart to recent observations in solid organ transplant recipients that document the longevity of donor-origin T_RM_ in transplanted lung and intestinal allografts (*5, 6, 28*). Our results suggest that within eight days after allo-HCT, donor T cells infiltrate target organs and rapidly exhibit both protein and transcriptional T_RM_ hallmarks. However, despite clear expression of a T_RM_ transcriptional and protein expression program, when compared to either host or HC GI-tract CD8 T cells, the donor CD8 T_RM_ are clearly not homeostatic. They upregulate multiple transcriptional networks driving T cell activation, including evidence of both metabolic reprograming toward mitochondrial respiration and massively upregulated cytotoxicity pathways. Our results also identify IL-15 and IL-21 as key up-stream regulators, which together coordinate the transcriptional programming responsible for the pathogenic differentiation of donor CD8 T_RM_ cells (**Figure 4H-I**). Notably, these two cytokines have been shown to drive aGVHD in mouse models (*48–51*) and IL-15 signaling is required for the generation of CD8 T_RM_ in multiple barrier tissues (*46, 66*). Compared to IL15, the role of IL-21 in the generation of T_RM_ cells is far less established (*47*), and in this study, we provide new evidence that IL-21 can drive T_RM_ differentiation from circulatory precursors during aGVHD, but not under homeostatic conditions (**Figure 4J**). The coordinate regulation of redundant T cell activation programs that this work has demonstrated may provide an explanation for the well-documented difficulty of reversing clinical aGVHD once it has occurred. The pathogenic GI-tract T_RM_ are not only physically sequestered from therapeutic agents (*67, 68*), they are also potentially protected from monomorphic treatment approaches, given the many non-overlapping pathways that are activated in these pathogenic cells. These non-overlapping pathways include upregulation of oxidative phosphorylation and mitochondrial respiration, activation of multiple co-stimulatory pathways (including ICOS and OX40), in addition to enrichment for several mediators of cytotoxicity, which, in their totality, could be exceedingly difficult to control with singly targeted therapies.

In addition to distinguishing the features associated with pathogenic donor CD8 T_RM_, we have also identified the first transcriptional network controlling donor CD8 T cells ‘caught in the act’ of tissue infiltration (**Figure 5**). These rare cells have previously been intractable to detailed examination, both because of their small numbers and because of their fleeting nature. Through the combination of SIVS and scRNAseq, we were able to identify and deeply interrogate this critical cell population. By enabling flow cytometric sorting of GI-tract T cells that were Compartment-1 (iVas)-negative, SIVS allowed us to rigorously exclude intravascular cells from our analysis. While Compartment-2 cells were too rare in the GI tract to flow cytometrically sort, the fact that these cells were rigorously demonstrated to be non-T_RM_ by flow cytometry (**Figure 3B**) allowed us to identify them by scRNAseq through their T_RM_-low transcriptional signature (**Figure 4C**). Transcriptional analysis subsequently identified a number of central features of these actively infiltrating cells. While these features comprised several ‘usual suspects’ in aGVHD, (including co-stimulatory/co-inhibitory pathways (*69*), the immunoproteosome (*70*) and several targetable metabolic pathways (*37, 38, 71*); **Figure 5A-D**) one of the most important findings was the transcriptional program of adhesion/extravasation/migration that these actively infiltrating cells initiated (**Figure 5E**). Their coordinate upregulation of multiple pathways of cellular transit distinguished these cells both from donor CD8 T cells that had already enacted T_RM_ programming, as well as from both host and HC Compartment-2 cells. Finally, the demonstration of significant overlap between NHP transcriptional signatures and those found in patients with active aGVHD underscores the clinical relevance of these findings. These results thus identify a new class of genes and pathways that characterize actively infiltrating pathogenic donor CD8 T cells, and which could play a major role in driving these cells into target organs prior to tissue destruction. The identification of gene networks associated with active tissue infiltration may have relevance to other immune phenomenon outside of aGVHD. These include solid organ transplant rejection as well as auto-immune diseases, which both require infiltration and long-term persistence of pathogenic T cells in target tissues (*72*). In addition, the discovery of a CD8 T cell infiltration gene network may lead to novel strategies for increasing the ability of anti-tumor T cells to more successfully infiltrate their targets and survive under the otherwise hostile tumor microenvironment, given the mechanistic connections between tumor-infiltrating lymphocytes and T_RM_ biology that have been recently discovered (*7, 8*).

In summary, by combining SIVS with scRNA-seq we have interrogated pathogenic donor CD8 T cells in the act of infiltrating and causing tissue destruction during GI aGVHD. Our results document the first transcriptional network defining T cell infiltration during this disease, and validate the clinical relevance of this network. They nominate a new class of targets by which to control the act of T cell infiltration, which constitutes a necessary prelude to T cell-mediated organ damage.

## Methods

### NHP Ethics Statement

This study was conducted in strict accordance with USDA regulations and the recommendations in the Guide for the Care and Use of Laboratory Animals of the National Institutes of Health. It was approved by the Biomere Inc and the University of Washington Institutional Animal Care and Use Committees.

### Study Design

This was a prospective study in NHP designed to determine the biological role of T cell migration and to identify the molecular signature of pathogenic tissue-infiltrating T cells in aGVHD-induced tissue immunopathology. Several cohorts of transplant recipients were studied: (1) Autologous transplants (abbreviated as ‘Auto-HCT’, n = 9 for clinical analysis and n=4 for immunological analysis); (2) Allogeneic transplants with no GVHD prophylaxis (abbreviated as ‘allo-HCT’ and ‘aGVHD’, n = 11 for clinical analysis and n = 4, including 3 MAMU-A001-mismatched transplantations); (3) A control cohort of healthy, immunologically naïve macaques (abbreviated as ‘HC’, n= 4 for immunological analysis). Animal demographic parameters, transplant characteristics, and doses of *α*CD45 antibodies, administered *in vivo* are shown in **Table S1**.

### Hematopoietic Cell Transplantations (HCT) in NHP

Transplant recipients and donors were chosen from breeding colonies based on their MHC-genotypes. For these studies, we utilized allele-specific MHC typing to choose donor: recipient pairs. All allo-HCT experiments utilized MHC haplo-identical unrelated (n=3) or half-sibling (n=1) donors and recipients (**Table S1**). Apheresis was performed after G-CSF mobilization (Amgen, 50mcg/kg for 5 days), and an unmanipulated G-CSF mobilized apheresis product was transplanted into MHC haplo-identical transplant recipients in the allo-HCT cohort, or was aseptically cryopreserved and administered to auto-HCT recipients 6-8 weeks after the apheresis product collection. The transplanted total nucleated cell dose (TNC) and CD3^+^ cell doses are shown in **Table S1**. The pre-HCT preparative regimen consisted of total body irradiation (TBI) of 10.4 Gy given in two fractions per day for two days (2.6 Gy per fraction). Irradiation was delivered using either a linear accelerator Varian Clinac 23EX with a tissue-adjusted dose rate of 7 cGy/min, (for all animals except R.312) or using gamma-irradiation with a Cobalt-60 isotope (for R.312) at a tissue-adjusted dose rate of 5.5 cGy/min. All transplants were performed with a central venous catheter placed for the length of the experiment. Antibacterial prophylaxis in all transplant recipients included Vancomycin (with a target serum concentration of 5-20 mcg/mL) and Ceftazidime, administered via central catheter, and Enrofloxacin (‘Baytril’), administered IM to all recipients with neutrophil counts less than 500 cells/μL. All transplant recipients received empiric antiviral (acyclovir, 10 mg/kg IV daily; cidofovir, 5 mg/kg IV weekly) and antifungal prophylaxis (fluconazole 5mg/kg oral or IV, given daily). ABO-matched leukoreduced (using an LRF10 leukoreduction filter, Pall Medical) and irradiated (2.2 Gy) platelet-rich plasma or whole blood was given for a peripheral blood platelet counts of ≤ 100 × 10^3^ per μL in the allo-HCT cohort or ≤ 50 × 10^3^ per μL in the auto-HCT cohort, or if clinically significant hemorrhage was noted.

The aGVHD clinical score was assessed daily for allo-HCT recipients as previously described(*19*) based on GI- (presence and severity of diarrhea), liver- (hyperbilirubinemia), and skin-specific (rash) abnormalities. Transplant recipients were not given immunosuppression when aGVHD was diagnosed. Animals were humanely euthanized when they met either the prespecified clinical endpoints, or at a pre-set experimental endpoint on day 8. Histopathologic scoring for aGVHD was performed using a previously validated semi-quantitative scoring system (Grades 0.5-4)(*19*) in the blinded manner.

### αCD45 infusions

Purified αCD45 (clone MB4-6D6) was purchased from Miltenyi Biosciences. Antibodies were conjugated in-house to AlexaFluor dyes. Conjugated antibodies were diluted in sterile Normal Saline Solution, and 5 mLs were slowly infused over 5 minutes to animals via a central catheter placed in the femoral vein (auto-HCT and allo-HCT cohorts) or to sedated animals via a peripheral catheter placed in saphenous vein (HC cohort). To track lymphocyte migration, αCD45 antibodies conjugated with AlexaFluor647 (αCD45-AF647) were administered 6 hours before euthanasia/tissue collection at a dose of 30 mcg/kg. To discriminate tissue-residing cells from cells remaining in the vasculature, αCD45 antibodies, conjugated with AlexaFluor488 (αCD45-AF488), were administered 5 minutes before euthanasia/tissue collection at a dose of 30 or 60 mcg/kg (**Figure 1**, **Figure S1**; **Table S1**). Blood samples were drawn from the catheter before, and 5 minutes after each αCD45 antibody injection to ensure that all blood leukocytes were uniformly labeled with the injected antibodies.

### Necropsy and tissue processing

Animals were euthanized followed by perfusion using sterile PBS (0.5 L/kg body weight), necropsy and tissue harvest. Blood PBMC were isolated using Ficoll-Paque PLUS gradient separation and standard procedures; lymph nodes, spleen and bone marrow (from long bones) were mechanically disrupted by grinding through a metal strainer, followed by filtering through nylon 70 μ and 40 μ cell strainers. Lungs, liver, jejunum colon, and kidney samples were cut into small pieces, then digested using 150 KU DNase (Sigma-Aldrich) with 1 mg/mL Type I collagenase (Invitrogen) (lungs, liver and kidney) or 150 KU DNase (Sigma-Aldrich) with 50 mcg/mL Liberase TL (Roche; colon and jejunum) at 37°C for 1 hour with shaking, followed by passing through a metal strainer, and then 100 μ, 70 μ and 40 μ nylon cell strainers. The resulting cell suspension was cleaned using a double-layer Percoll (GE Healthcare) gradient separation. The leukocyte fraction was collected at the interphase between 70% and 30% Percoll. Cells were either washed and immediately analyzed or were cryopreserved in 10% DMSO/90% FBS and stored in liquid nitrogen.

### Immune Analysis

Blinding was performed on all pathologic analysis. The analysis of flow cytometry data and single-cell RNA-Seq data was performed in an unblinded manner.

### Multiparameter Flow Cytometry

Samples were processed as described and either analyzed fresh or from previously cryopreserved samples. Briefly, cells were washed with PBS and incubated with 200 uL 1:100 LIVE/DEAD AQUA (Invitrogen) for 20 minutes at 37°C using 96- well cell culture plates. After viability staining, cells were washed with staining buffer (PBS supplemented with 2% heat-inactivated FBS) and then stained with extracellular antibody cocktails (**Table S2**) at 4°C for 20 minutes, followed by washing with the staining buffer. Samples, stained with extracellular antibodies only, were fixed with 1X BD Stabilizing Fixative and run within 24 hours after staining. Samples requiring intracellular staining were incubated in 250 μL Cytofix/Cytoperm solution (BD Biosciences) at 22°C for 20 minutes, followed by washing with 1X Perm/Wash buffer (BD Biosciences) and then incubated with intracellular antibody cocktails, mixed in 50 μL of the 1X Perm/Wash buffer, at 4°C for 30 minutes (**Table S 2**). These samples were then washed and analyzed within 24 hours after staining. Samples requiring intranuclear staining were incubated in 250 μL FoxP3 fixation buffer (BioLegend) at 22°C for 20 minutes, followed by a washing step with FoxP3 wash buffer (BioLegend) and then incubated with intranuclear antibody cocktails, mixed in 50 μL of the FoxP3 wash buffer, at 4°C for 30 minutes (**Table S2**). These samples were then washed and analyzed within 24 hours after staining. Flow cytometry was performed on a BD FACS LSRFortessa and analyzed in FlowJo v.10. The gating tree and definitions of the major analyzed T cell populations are shown in **Figure S2** and **Table S2**.

### Measurement of cellular migration using in vivo aCD45 labeling

To estimate the extent of CD4 or CD8 T cell migration, we measured the number of corresponding T cells that had migrated into different tissues within 6 hours prior to necropsy. To accomplish this, we assessed the proportion of Compartment-2 CD4 or CD8 T cells, positive for aCD45-AF647, compared to all extravascular/tissue-residing Compartment-2 + Compartment-3, IVas^-^ CD4 or CD8 T cells (negative for *α*CD45-AF488, administered 5 minutes before euthanasia/tissue collection). To estimate donor- and host-specific CD4 or CD8 T cell migration in the allo-HCT cohort, we assessed donor and host origin flow cytometrically using a fluorescent-MAMU-A001 antibody, where possible (**Figure S2**).

We also calculated the difference between donor and host migration (*Δ*Migration) by subtracting the amount of host CD8 T cell migration from the amount of donor CD8 T cell migration (both measured as the percent of recently infiltrated ‘Compartment-2’ host or donor CD8 T cells, respectively, out of total extravascular IVas^-^ CD8 T cells, using the following formula:

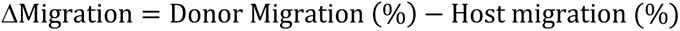

Next, we calculated the rate of donor CD8 T cell conversion into T_RM_ cells (referred as ‘Retention’) by calculating the ratio between the percentage of donor CD8 T cells within the resident Compartment-3 (expressing the CD69^+^CD103^+^ T_RM_ phenotype) and the percentage of host CD8 T cells within the resident Compartment-3 (expressing the CD69^+^CD103^+^ T_RM_ phenotype) as follows: To calculate migration that was adjusted for the rate of T_RM_ differentiation from infiltrating precursors, (referred to as ‘(*Δ*Migration x Retention’), we multiplied the *Δ*Migration by the rate of donor CD8 T cell conversion into T_RM_ cells, as described above. These calculations were performed using the following formula:

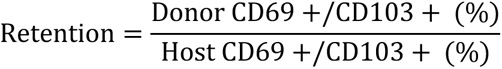

### In vitro stimulation with cytokines

Viably frozen spleen cells from HC animals or from aGVHD recipients on day +8 post-HCT were thawed and cultured in 96-well flat-bottom plates at a density of 2×10^5^/well for 48 hours in the presence or absence of the following cytokines (all purchased from RnD systems, Minneapolis, MN): IL-15 (50 ng/mL), IL-21 (50 ng/mL), TGF*β*1 (20 ng/mL). At the end of culture, cells were harvested and stained for viability using LIVE/DEAD Aqua stain (Invitrogen), and cell surface markers. Expression of CD69 and CD103 was measured in LIVE/DEAD aqua^-^/CD3^+^/CD14^-^/CD20^-^/CD8a^+^/CD4^-^ viable CD8 T cells.

#### Single-cell RNA-sequencing (scRNA-seq) sample preparation, library construction and sequencing

Four colon cellular preparations from the allo-HCT cohort and four colon cellular preparations from the untransplanted HC cohort were aseptically thawed, labeled with the viability dye LIVE/DEAD Violet (Invitrogen), then stained with *α*CD45 antibodies, conjugated with PerCP-Cy5.5 (BD Biosciences). For all samples, tissue-associated T cells were rigorously identified by sorting only CD45-AF488 (IVas)-negative cells. Viable LIVE/DEAD Violet-negative CD45-positive cells were sorted from two out of four GVHD-derived samples into host and donor fractions, based on MAMU-A001 expression using PE-conjugated *α*MAMU-A001 antibodies (NHP Reagent Resource). In the two GVHD samples for which antibodies were not available to distinguish disparate donor- and host-MHC alleles, donor- and host-derived cells were defined computationally based on male/female disparity (see the corresponding **Methods** section, below). Single cell libraries from sorted samples were generated using a UMI-based droplet-partitioning platform with the Chromium single cell 3’ library and gel bead kit v2 and v3 (10X Genomics) and then sequenced using a NovaSeq S2 (Illumina). Technical information relevant to the scRNA-Seq samples is found in **Table S3**.

#### Alignment and QC of scRNAseq Data

We used genome assembly MMul_10 as our reference genome for macaca mulatta and used Ensembl’s (v98) (*73*) annotations for this genome to identify genes. We used cellranger v3.1.0 (https://support.10xgenomics.com/single-cell-gene-expression/software/pipelines/latest/what-is-cell-ranger) “mkgtf” and “mkref” commands to construct a reference transcriptome. We used FASTQC (https://www.bioinformatics.babraham.ac.uk/projects/fastqc/) to assess the quality of our fastq files. We aligned the data using cellranger’s “count” command, and then combined the samples using the aggregate command. During the aggregation step, we set “normalization=none” to retain all data for filtering and normalization in the next step.

Using Seurat v3.1, we filtered out genes that met any one of the following conditions: present in less than 200 cells (comprising ∼1% of the final cell count after filtering), lacked both a gene symbol or description in ensembl, were not protein-coding, or were lncRNA’s (with the exception of two genes we subsequently used to identify males and females). The stringent threshold of removing genes present in <200 cells was stable compared to less stringent thresholds, with similar results obtained when we removed genes present in <5 cells. Following the recommendations in Lucken and Theis (*74*), we analyzed the total transcripts detected and total genes detected for each sample to determine appropriate thresholds, and retained cells with 300 to 3,000 genes detected and 1,200 to 20,000 transcripts detected. The version of Ensembl used for this analysis did not contain genes mapping to mtDNA, but the genes ENSMMUG00000028699 and ENSMMUG00000028695 appeared to be mitochondrial, based on their high abundance in low-quality cells. Furthermore, Ensembl orthologs for ENSMMUG00000028699 map to ND1 on mtDNA for humans and other primates, and while no orthologs are listed for ENSMMUG00000028695, the associated UniProt entry for ENSMMUG00000028695 is ND2 (Q6IYH7), a mitochondrial gene. Therefore, we used these genes as proxies for mtDNA, and removed any cells where > 0.5% of the expression mapped to these genes. This filtering resulted in 21,490 cells for downstream analysis. We then normalized the data using SCTransform in Seurat, and included “batch” as the batch variable to correct for the two batches of sequencing.

#### Identification of Donor and Host Samples within Sex Mismatched Transplants

A subset of samples contained host and donor cells obtained from sex-mismatched transplants. To find candidate genes to disaggregate these by gender, we performed a Wilcoxon test on the other samples, using Seurat’s FindMarkers command to identify genes that were highly differential between male and female cells. We found that 97.7% percent of our known male cells expressed RPS4Y1, and only 0.6% of our known female cells had detectable expression for this gene, and that 58.4% of our male cells expressed RPS4Y2 and 0.3% of our female cells expressed this gene. Using expression of either of these genes to mark samples as male, 98.2% of our known male cells were marked as male, and 99.3% of our known female cells were marked as female. Satisfied with this performance, we used expression of either gene to mark cells as male, and assigned them to the corresponding donor or host status for the sample.

#### Clustering of Cells and Identification of T Cells

We next ran principal component analysis (PCA) to reduce the dimensionality of the dataset, and ran Seurat’s FindNeighbors, FindClusters (resolution = 0.2), and RunUMAP methods on the first 22 principal components to identify clusters in the data. We retained clusters enriched for TRAC, CD3E, IL2RA, CD4, CD8A, or IL7R as our T cell clusters. Our T cell dataset contained 14,185 cells. We then filtered out any genes that were detected in less than 100 cells, or only detected in one batch and not the other.

#### Clustering of T Cell Dataset and Identification of CD8 T Cells

As above, we ran SCTransform and PCA on this dataset, and then FindNeighbors, FindClusters (resolution = 0.8), and RunUMAP using the first 29 principal components for the T cell dataset. Since *CD8* expression is not always detectable in CD8 T cells, we identified CD8 T cells in the T cell dataset by assessing their scores on the “GSE7460_CD8_TCELL_VS_CD4_TCELL_ACT” signature from Hill JA et al, *Immunity* (*2007*) (*31*) in MSigDB’s C7 signatures, using the program VISION (*30*). VISION calculates a score for each cell in the dataset, and then identifies clusters positively or negatively enriched for that cluster by applying a Wilcoxon test between a given cluster and the rest of the dataset (VISION converts the Wilcoxon statistic to an Area Under The Curve (AUC) value). We retained clusters that were positively enriched for the signature, by starting with the most enriched cluster, and adding on subsequent clusters, and conducting a Wilcoxon Test of the candidate CD8 T cells versus all remaining cells and converting this test statistic to an AUC. Our CD8 T Cell dataset contained 4,715 cells and had an AUC of 0.847.

#### Differential Expression Analysis of Donor, Host, and Healthy Control CD8 T Cells

We used Wilcoxon Rank-Sum tests in Seurat to conduct differential expression analysis on pairwise comparisons of our three conditions. We only tested genes where at least 25% of the cells in one of the conditions had detectable expression for the gene, and where there was a log fold change of at least 0.25.

#### Identification of T_RM_ High and Low Cells

We scored the 4,715 CD8 T cells by using VISION to apply a T_RM_-signature adopted from *Milner et al, Nature, (2017)* (*32*). We obtained the 25^th^ and 75^th^ percentiles for this score on the HC CD8 T cells, and marked all CD8 T cells below the 25^th^ HC percentile as T_RM_ Low and above the 75^th^ HC percentile as T_RM_ High. We used the FindMarkers command in Seurat to identify DE results in pairwise comparisons, again only testing genes that were expressed in 25% of cells in HC, with a log fold change threshold of 0.25. To ensure that our results were not sensitive to our choice of 25% cutoff values for classifying T_RM_-High and T_RM_-Low groups, we tested several choices of bounds in increments of 5%, (between 15% and 35%) and investigated the DE gene lists. Results for the bounds 20%, 25% (used for analysis) and 30% were similar (**Figure S7**).

#### Functional annotation of genes, pathway analysis and analysis of up-stream regulators

Pathway analysis on the cluster-specific genes or differentially expressed genes was performed using Ingenuity Pathway Analysis (IPA; https://www.qiagenbioinformatics.com/products/ingenuity-pathway-analysis/). Pathways with p values less than 0.01 for Benjamini-Hochberg-corrected t-test were considered statistically enriched. Up-stream regulators (limited to the following terms: growth factors, cytokines, G-coupled receptors, transmembrane receptors, transcriptional regulators, ligand-dependent nuclear receptor) were considered activated or inhibited if their enrichment z-scores were greater than 1.25 or less than −1.25, respectively, and p values were less than 0.05 using the Benjamini-Hochberg-corrected t-test.

#### Alignment of Human CD8 T Cell RNA-Seq

We obtained single-end bulk RNA-Seq from 69 human CD8 T cell samples from patients in the placebo-control arm of the ABA2 clinical trial (NCT# NCT01743131). These patients were transplanted using standard calcineurin inhibitor/methotrexate aGVHD prophylaxis (**Table S.13**) We assessed the quality of the RNA-Seq using FastQC, and trimmed low quality bases and adapters using Trimmomatic(*75*) v.39 with the settings “-phred33 ILLUMINACLIP:Nextera-SE.fa:2:30:10 LEADING:15 TRAILING:15 SLIDINGWINDOW:4:15 MINLEN:80” We built a reference transcriptome with kallisto v0.46.1(*76*), following the instructions at https://github.com/pachterlab/kallisto-transcriptome-indices/blob/master/README.md and using the cDNA file for Ensembl99. We aligned the RNA-Seq data to this transcriptome with kallisto using the settings “--threads 4 --single --fragment-length 357.8 --sd 29.7-b 100”.

#### Differential Expression Tests

We used DESeq2 (*77*) to conduct differential expression analysis between samples from patients with Grades 3-4 aGVHD versus Grades 0-2 aGVHD and Grades 2-4 aGVHD versus Grades 0-1 aGvHD. We filtered the gene set to protein-coding genes, and limited the genes to those with at measured abundance of at least 10 total across all samples. The models we fit were “∼ Batch + Grade_2to4_GvHD”, ““∼ Batch + Grade_3to4_GvHD”, and “∼ Batch + GvHD_Grade”, to run each of our tests. For the last model, our test was a contrast between GvHD grades 3-4 and 0-1.

GSEA: We used clusterProfiler(*78*) to conduct gene set enrichment analysis, which implements fGSEA(*79*). Rhesus macaque genes were mapped to human orthologs using the “hsapiens_homolog_associated_gene_name” attribute from ensembl.

#### Statistical analysis for flow cytometry experiments

Both paired and unpaired Student’s t-test were used for comparing two groups, where appropriate. One-way ANOVA with Holm-Sidak multiple comparison post-test or two-way ANOVA analysis with Holm-Sidak multiple comparison post-test were used for comparing multiple groups, where appropriate. Groups were considered as significantly different when p<0.05.

## ACKNOWLEDGEMENTS

**Funding:** This project was funded by grants to the following authors: LSK (NIH 2U19 AI051731, NIH 2R01 HL095791, NIH/FDA, R01 FD004099, CURE Childhood (https://curechildhoodcancer.org), BW (NIH K23HL136900), BRB (NIH R01 HL56067; NIH R37 AI34495), AKS (Sloan Fellowship in Chemistry, NIH 2R01 HL095791) and JOM (Richard and Susan Smith Family Foundation). The authors gratefully acknowledge the veterinary and animal husbandry staff at the Washington National Primate Research Center.

## Author Contributions

Conceptualization: VT, MR, LSK; Data Curation: VT, SNF, ELP, JK, DJH, CMG, BRB, JOM, AKS, MR, LSK; Formal analysis: VT, SNF, ELP, JK, APM; Funding

acquisition: LSK, AKS, BRB, JOM; Investigation: VT, SNF, ELP, DJH, CMG, HZ, LC, JC, MH, JO, AB, APM, BW, MQ, AL, JH, BB, KB, YS; Methodology: VT, ELP, MR, LSK; Project

administration: VT, AY, MH, LSK; Resources: MR, LSK; Supervision: MR, LSK; Visualization: VT, JK, LSK; Writing – original draft: VT, LSK; Writing – Review & editing: all authors.

## Data availability

The NHP scRNAseq data discussed in this publication were deposited in the NCBI’s Gene Expression Omnibus database (GEO …). The RNA-Seq data from ABA2 clinical trial discussed in this publication were deposited in the NCBI’s Gene Expression Omnibus database (GEO …).

## Supplemental data

**Figure S1.**
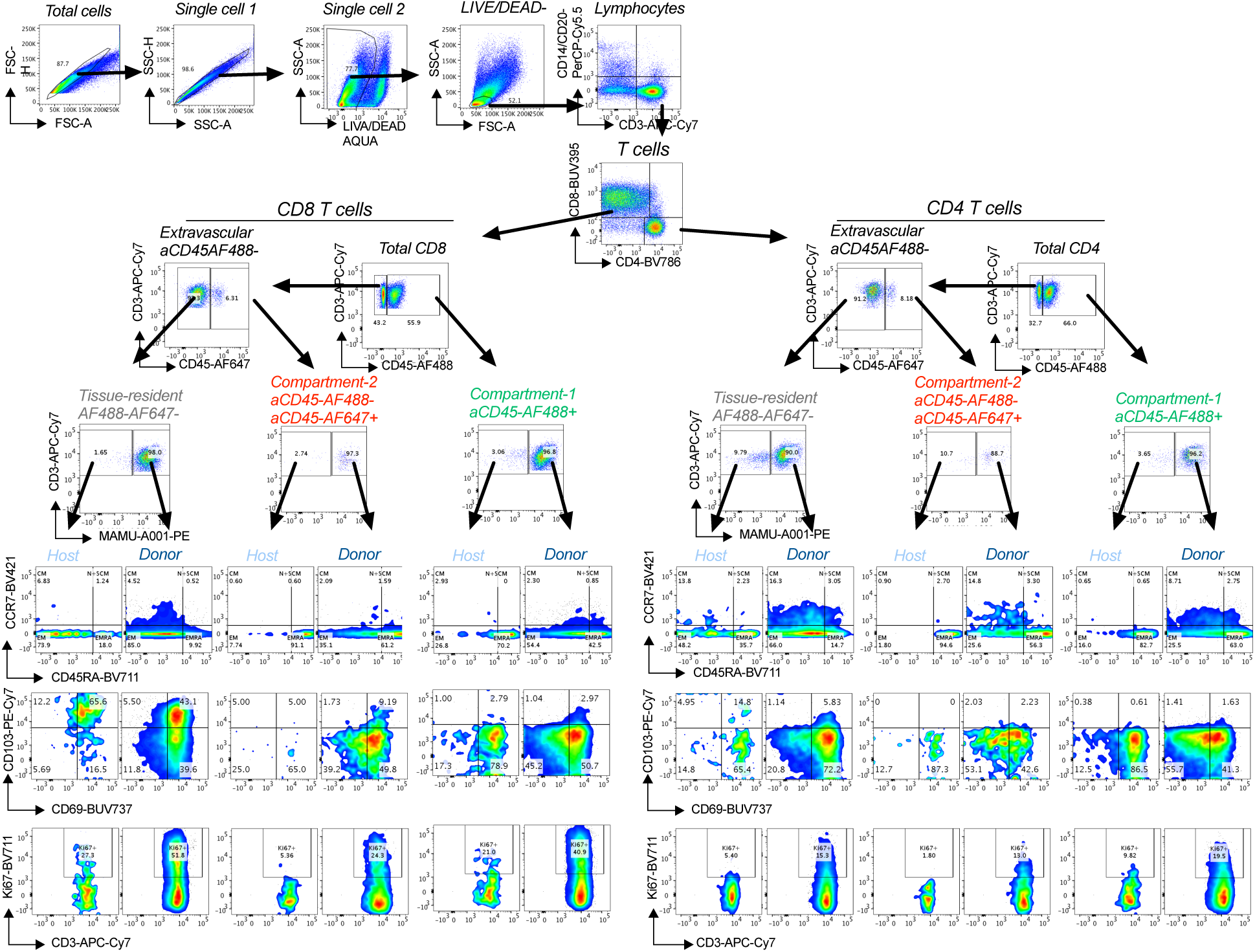
Gating tree for flow cytometric identification and characterization of T cells within different spatiotemporal compartments using Serial Intravascular Staining (SIVS). Flow cytometry plots show representative data from a healthy control lung sample. Populations gated on the plots are shown on top of each panel in *Italics*.

**Figure S2.**
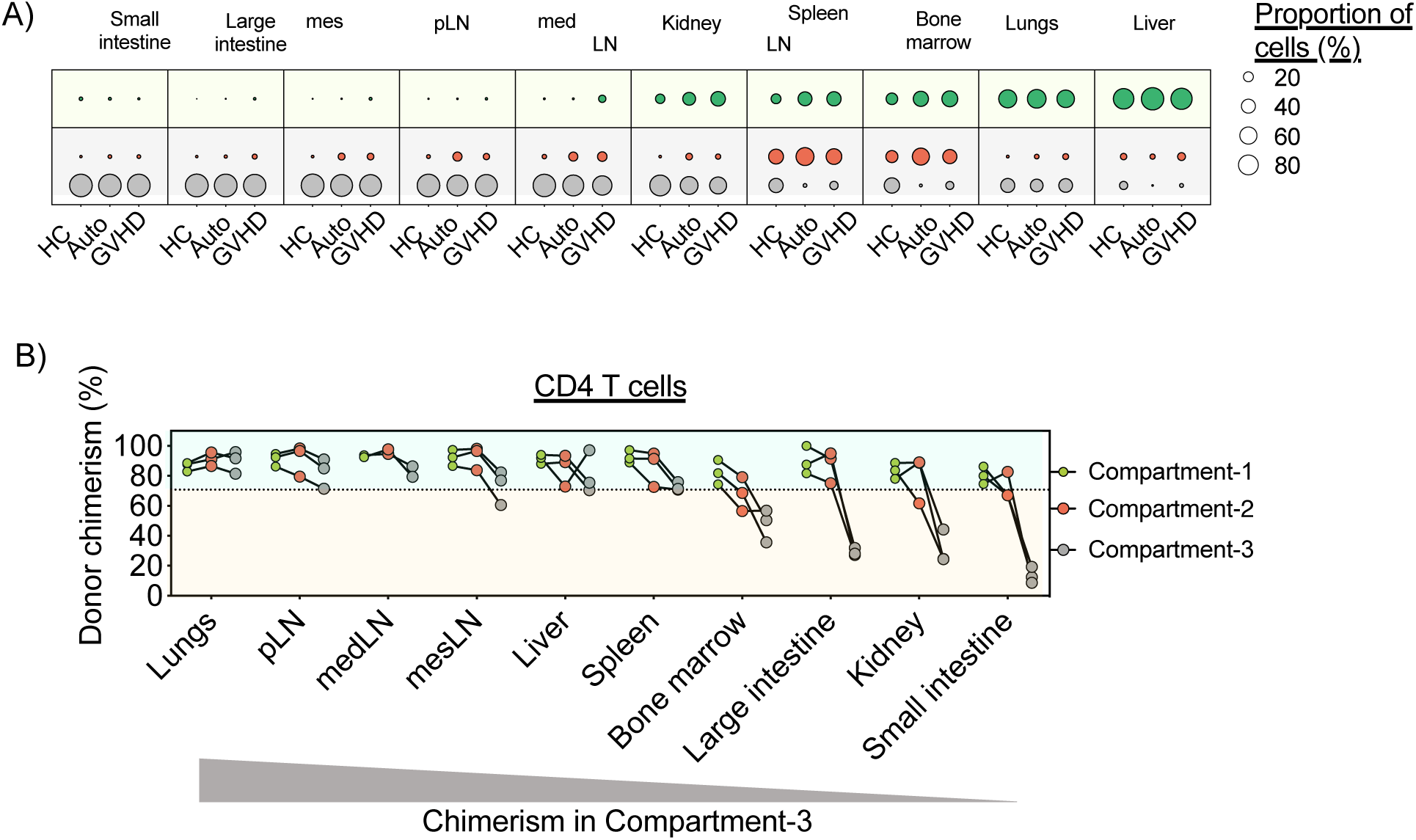
Compartmentalization of CD4 T cells at steady state and after HCT. A) Relative distribution of CD4 T cells between intravascular (Compartment-1; green dots), the recent infiltrating cell compartment (Compartment-2; red dots) and the resident compartment (Compartment-3; gray dots) in the indicated organs of healthy control animals (HC; n=4), auto-HCT cohort (auto-HCT; n=4) and the allo-HCT cohort (GVHD; n=4) on Day +8 post-transplant. Dot sizes indicate the proportion of cells within each compartment, compared to the total number of cells. pLN – peripheral Lymph Nodes, medLN – mediastinal Lymph Nodes, mesLN – mesenteric Lymph Nodes, SPL – spleen, BM – bone marrow, SI – small intestine (jejunum), LI – large intestine (colon). B) Percentage donor chimerism on day +8 following MAMU-A001-mismatched MHC-haploidentical allo-HCT (n=3) of CD4 T cells in different compartments: Compartment-1 (Green dots), Compartment-2 (Red dots) and Compartment-3 (Gray dots). The gray triangle at the bottom of the figure illustrates the pattern of donor chimerism in the tissues examined.

**Figure S3.**
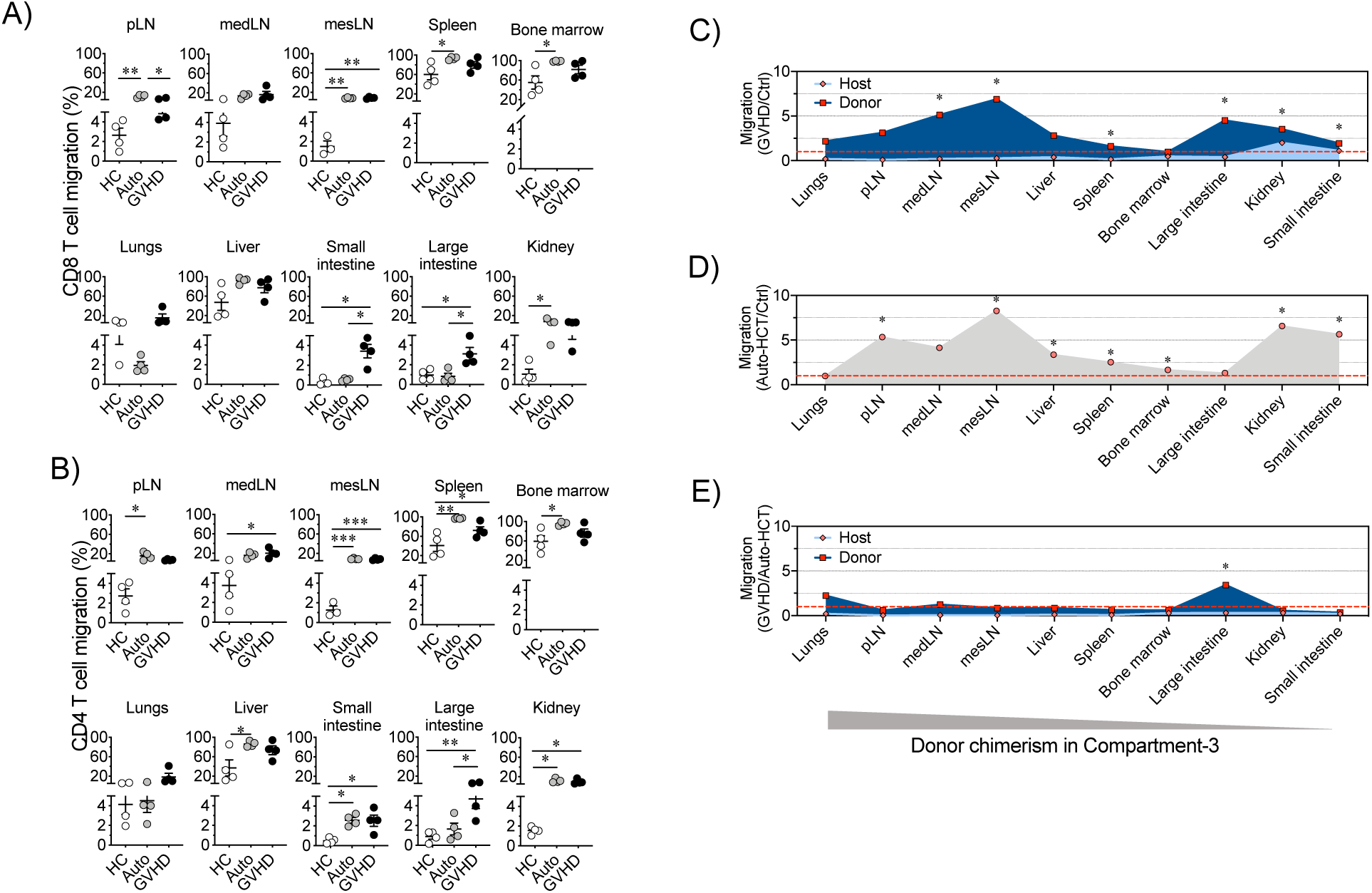
Migration of CD4 and CD8 T cells at steady state and after HCT. A-B) CD8 (A) and CD4 (B) T cell migration, measured as the percent of recently migrated (Compartment-2) cells out of all extravascular IVas^-^ cell (Compartment-2 plus Compartment-3 cells) at steady state in the healthy control cohort (n=4), on Day +8 in the auto-HCT cohort (n=4) and on Day +8 in the allo-HCT cohort (n=4) in different organs and tissues. *p<0.05, **p<0.01 and ***p<0.001 using a one-way ANOVA with the Tukey multiple comparison post-test. C) Relative migration of donor and host CD4 T cells in different organs and tissues on Day +8 following MAMU-A001-mismatched transplantation from the allo-HCT cohort (n=3) normalized to migration of CD4 T cells in healthy control animals (n=4). D) Relative migration of CD4 T cells on Day +8 from the auto-HCT cohort (n=4) normalized to the migration of CD4 T cells in healthy control animals (n=4). E) Relative migration of donor and host CD4 T cells in different organs and tissues on Day +8 following MAMU-A001-mismatched allo-HCT (n=3), normalized to the migration of CD4 T cells from the auto-HCT cohort on Day +8 (n=4). The gray triangle at the bottom of the figure illustrates the pattern of donor chimerism in the tissues examined. pLN – peripheral Lymph Nodes, medLN – mediastinal Lymph Nodes, mesLN – mesenteric Lymph Nodes.

**Figure S4.**
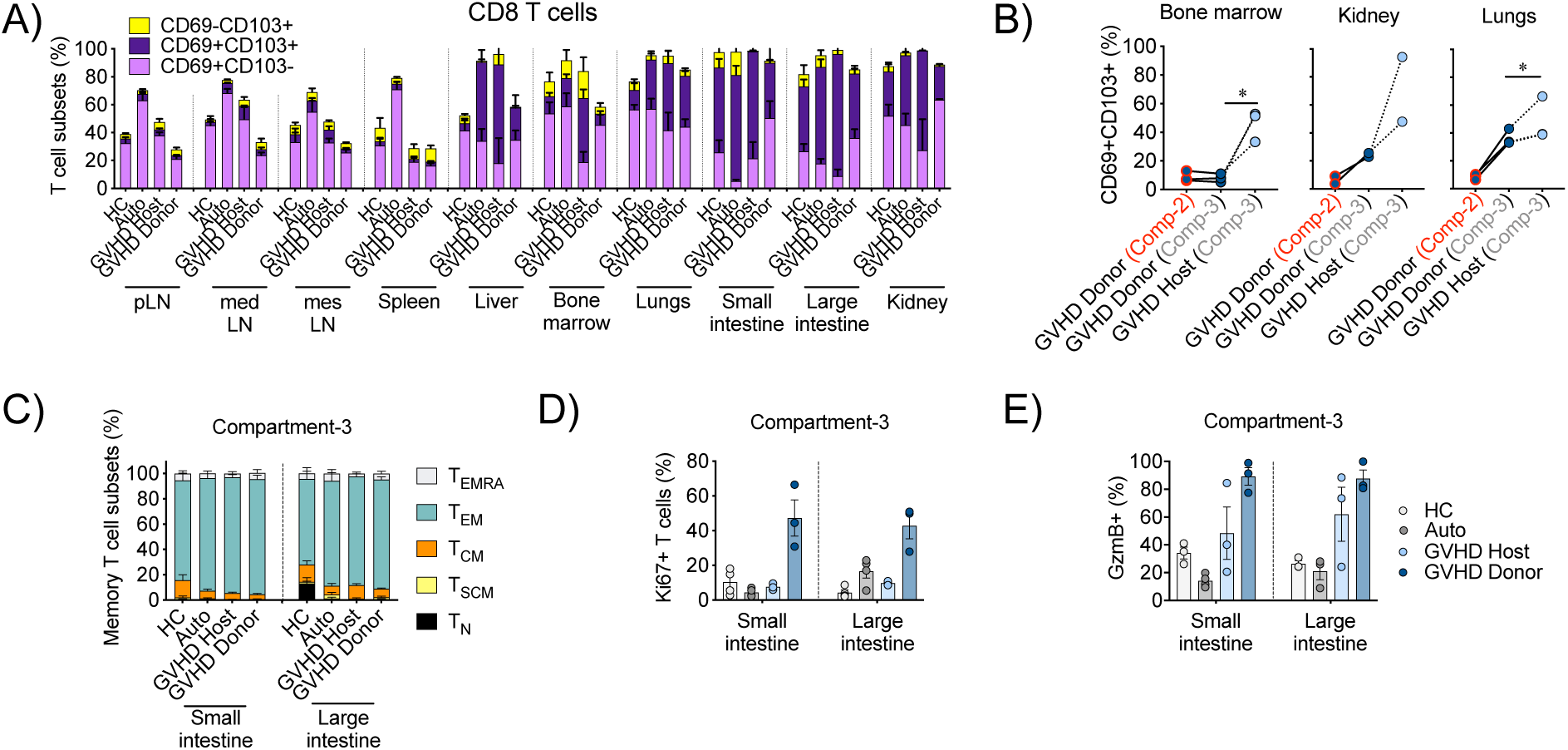
Expression of the T_RM_ markers CD69 and CD103 by CD4 and CD8 T cells in different organs and tissues at steady state and following HCT. A) Proportions of CD8 T cells with CD69^+^CD103^-^, CD69^+^CD103^+^ and CD69^-^CD103^+^ phenotypes within the resident compartment (Compartment-3) in different organs and tissues in healthy control animals (HC, n=4); in the auto-HCT cohort on Day +8 (Auto, n=4); host and donor T cells in the allo-HCT cohort on Day +8 following MAMU-A001-mismatched allo-HCT (n=3). B) Percentage of donor or host CD8 T cells with the CD69^+^CD103^+^ phenotype within recent infiltrating cells (Compartment-2) and resident cells (Compartment-3) in the indicated organs, plotted in a pair-wise fashion. Lines connect corresponding values were obtained from the same animal. *p<0.05 using paired t-test. C) Distribution between the memory subsets of CD8 T cells within the tissue-resident compartment (Compartment-3) in the small and large intestines in healthy controls (HC, n=3), the auto-HCT cohort on Day +8 (Auto, n=4) and host and donor cells of MAMU-A001-mismatched HCT from the allo-HCT cohort on Day +8 (n=3). D-E) Expression of Ki67 (D) and Granzyme B (E) in CD8 T cells in the resident compartment (Compartment-3) in the small and large intestines in healthy controls (HC, n=3), the auto-HCT cohort on day +8 (Auto, n=4) and host and donor cells of MAMU-A001-mismatched HCT from the allo-HCT cohort on day +8 (n=3).

**Figure S5.**
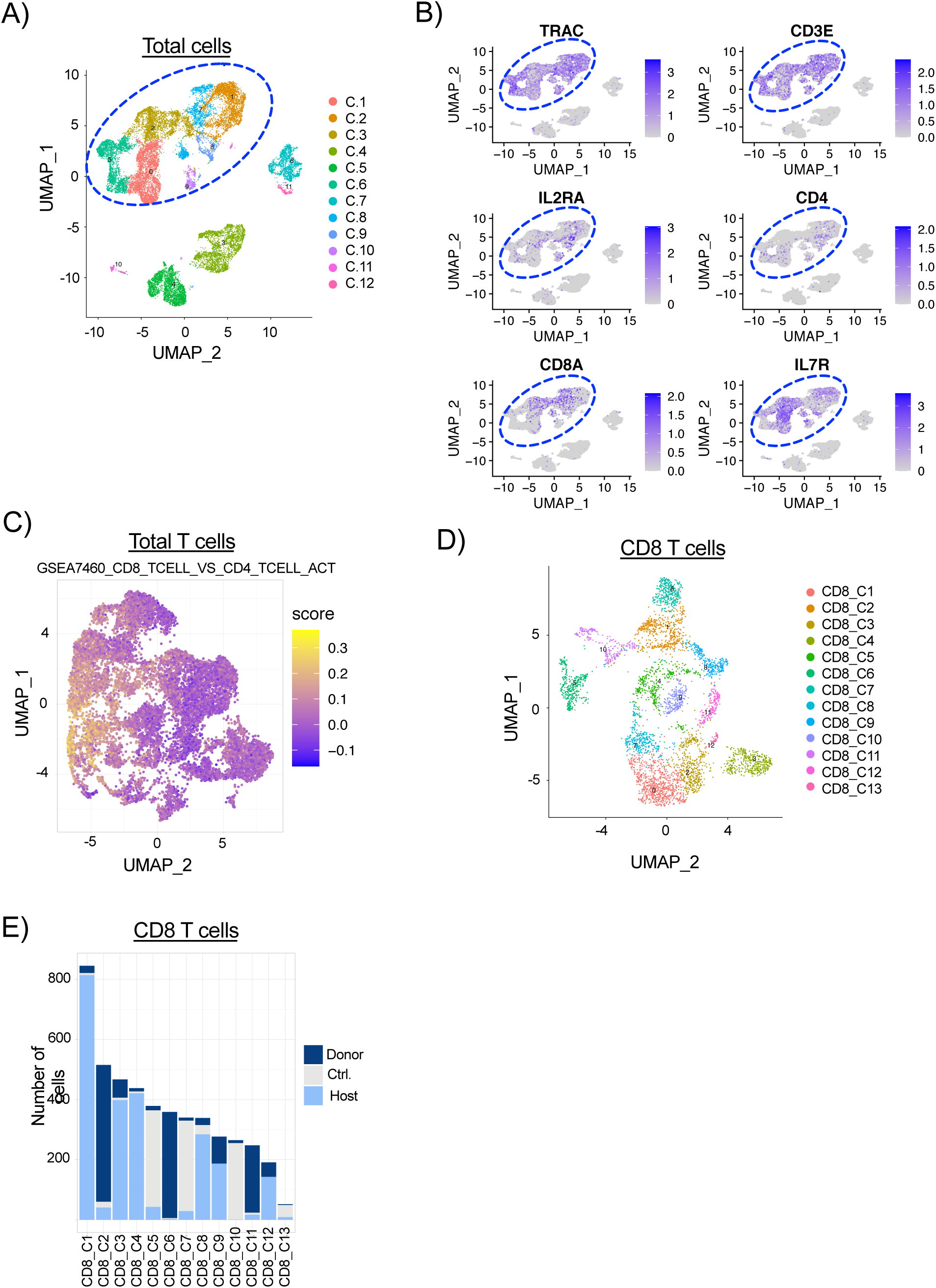
Transcriptomic identification of intestinal CD8 T cells. A-B) UMAP plots of the total single cell census (n= 14,185) after QC and filtering. Samples were sorted from four healthy control samples and two GVHD samples as LIVE/DEAD AQUA^-^ CD45AF488(IVas)^-^ CD45^+^ cells and as LIVE/DEAD AQUA^-^ CD45AF488(IVas)^-^ CD45^+^ MAMU-A001^-^ or MAMU-A001^+^ cells from two GVHD samples; see **Methods** for details). Cells are colored by clusters (A), or by expression of T cell-specific transcripts (B). Blue circle indicates T-cell clusters, which were used for further analysis. C) UMAP plot of T cell census, colored by CD8-signature enrichment scores. Cells are colored by CD8-signature enrichment score, adopted from Hill J.A. et al, (2007) Immunity(*31*) and applied with VISION(*30*). D) UMAP plot of CD8 T cell census, colored by clusters. E) Number of cells of donor, host or healthy control origin in each CD8 T cell cluster.

**Figure S6.**
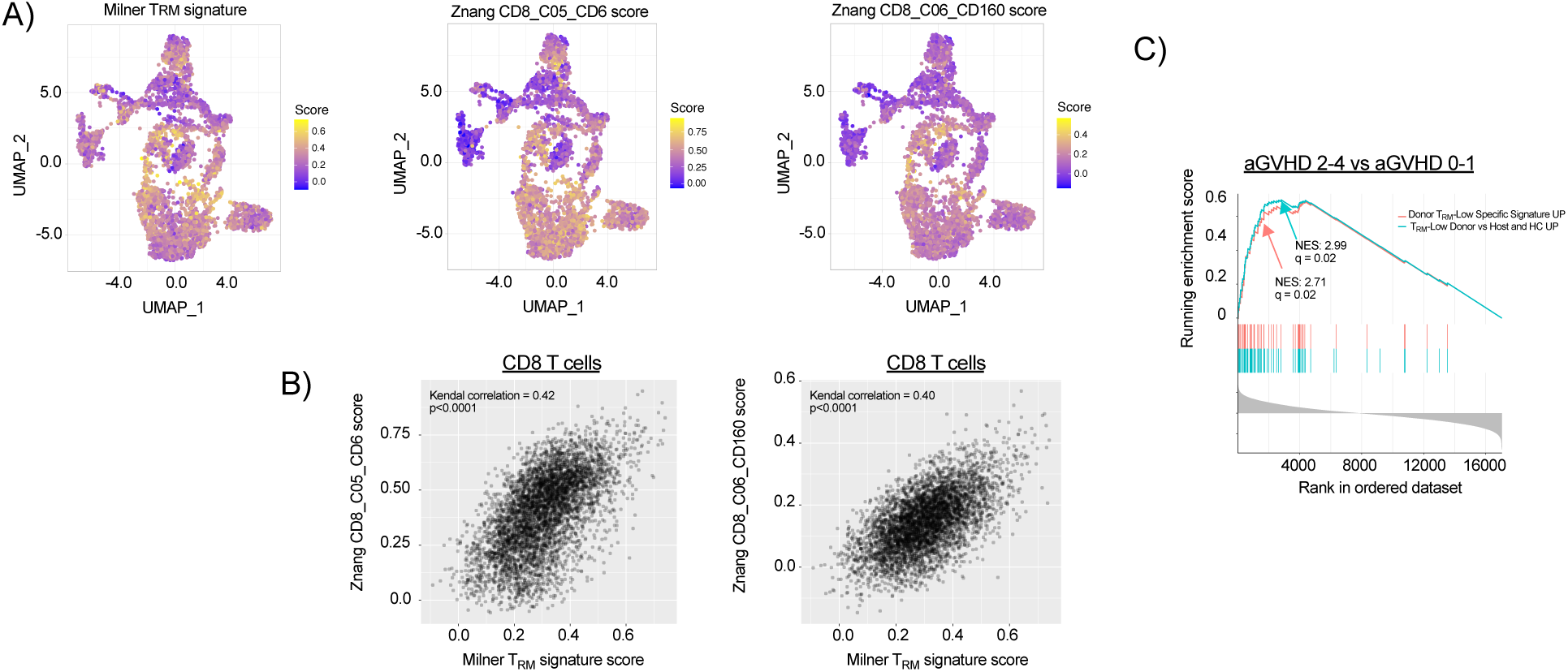
Transcriptomic characteristics of CD8 T cells from the large intestine. A) CD8 T cell census (n=4,715), clustered in Uniform Manifold Approximation and Projection (UMAP) space. Cells are colored by the enrichment score for the T_RM_-signature, adopted from *Milner et al, Nature,* (*2017*)(*30*) (*left panel*), and by the enrichment scores for the indicated T_RM_- related signatures adopted from *Zhang et al, Nature (2018)* (*31*) *(central and right panels)*. B) Correlation between the T_RM_-signature adopted from *Milner et al, Nature, (2017)*(*30*) and the indicated T_RM_-related signatures adopted from *Zhang et al, Nature (2018)* (*31*). C) GSEA plot depicting the enrichment of two T_RM_-low gene signatures compared to a clinical aGVHD CD8 T cell transcriptional signature. The curve labeled “T_RM_-Low Donor vs Host and HC UP” encompasses genes over-represented in donor T_RM_-low CD8 T cells in comparison with both host and HC T_RM_-low counterparts – blue line). The curve labled “Donor T_RM_-Low **specific** signature UP” encompasses genes over-represented **exclusively** in the donor T_RM_-low CD8 T cell comparison (i.e. not also present in the comparison of donor T_RM_-high CD8 T cells versus both host and HC T_RM_-high counterparts) – red line). The gene lists for the “T_RM_-Low Donor vs Host and HC UP” and the ““Donor T_RM_-Low **specific** signature UP” comparisons are found in **Table 15**. Each of these gene lists was used to compare patients with Grade 2-4 aGVHD and patients with Grade 0-1 aGVHD through GSEA. NES – normalized enrichment score.

**Figure S7.**
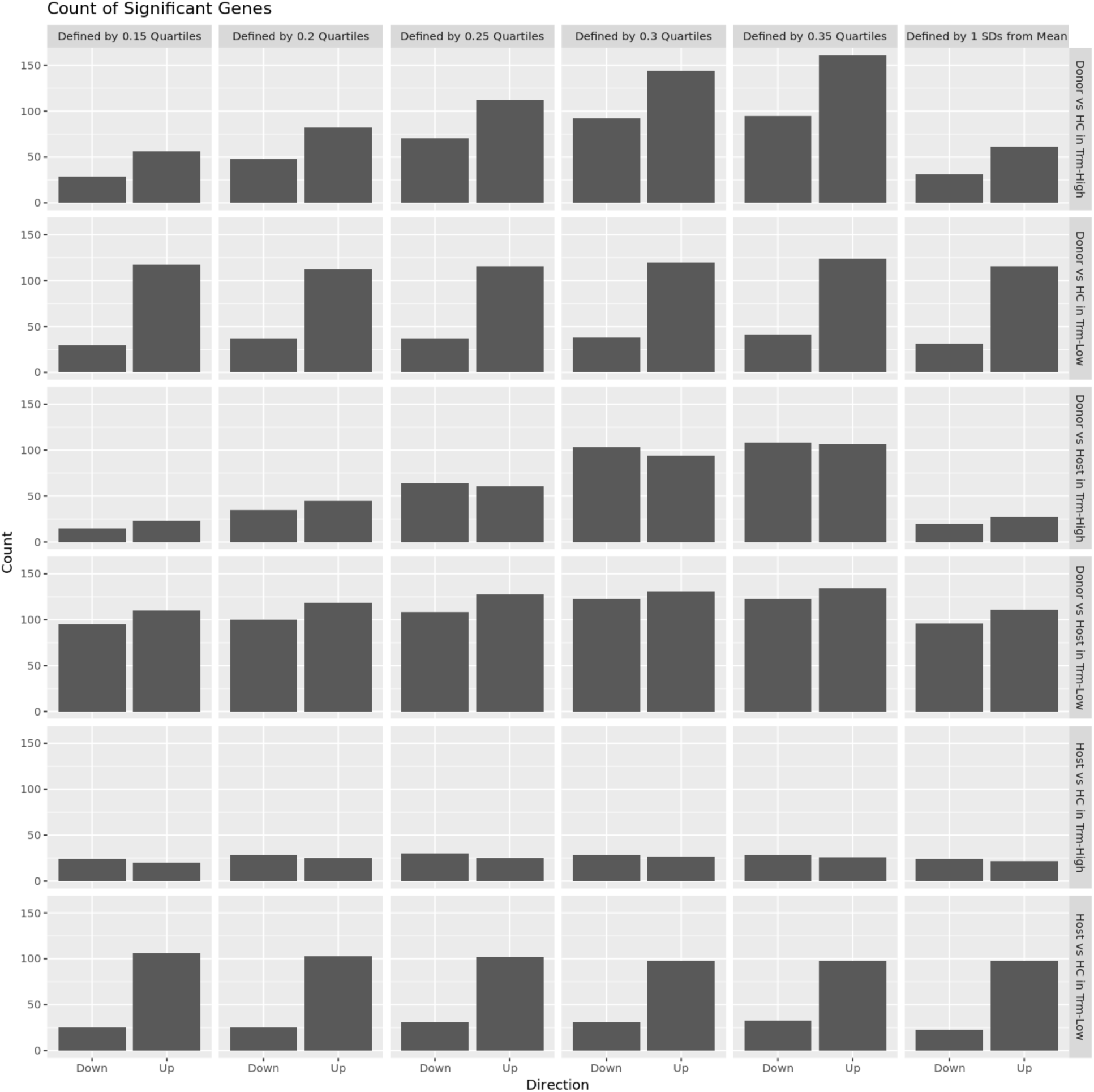
Threshold testing of T_RM_-High and T_RM_-Low CD8 T cells. Number of differentially expressed (DE) genes between donor, host and HC CD8 T cells with high and low enrichment scores for the T_RM_ signatures, adopted from *Milner et al, Nature, (2017)*(*30*) using different cutoff values for defining T_RM_ high and T_RM_ low.

**Table S1.**
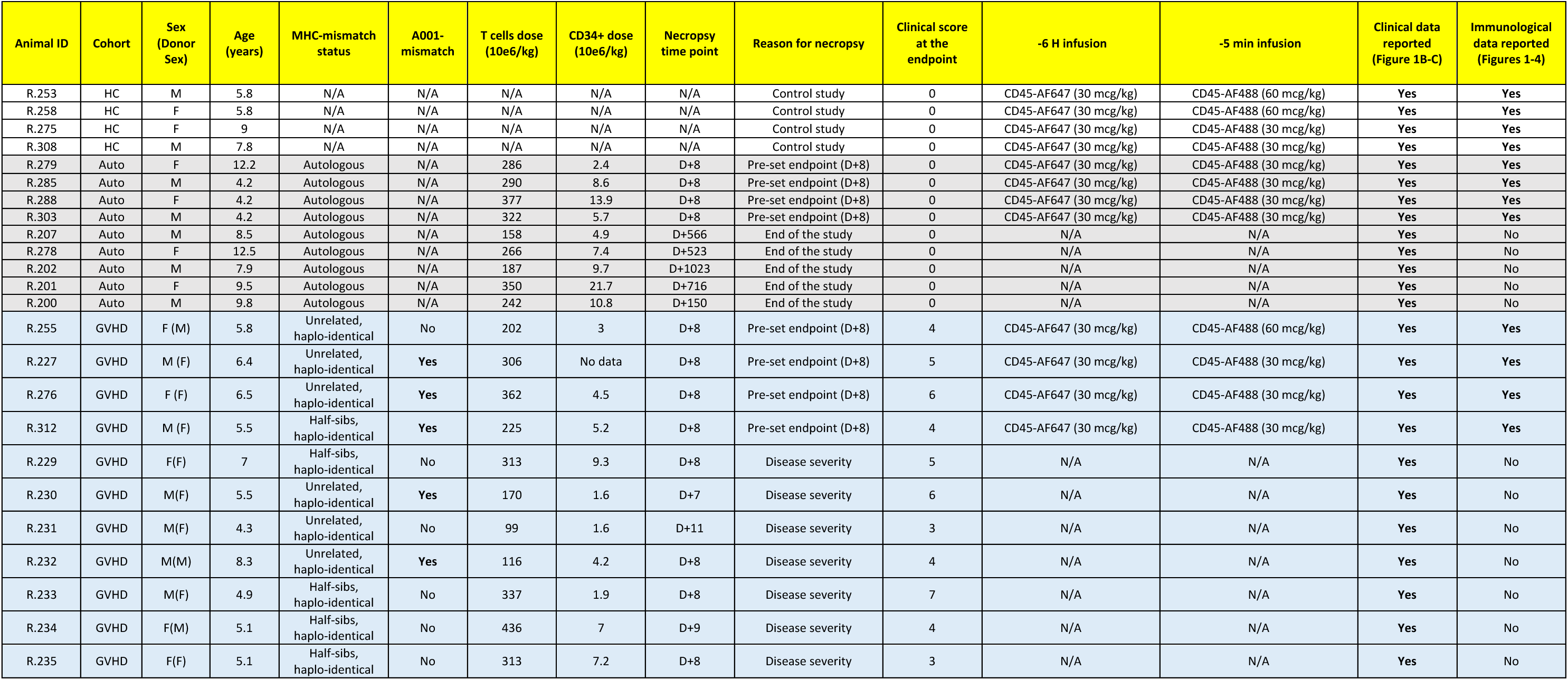
Transplant characteristics

**Table S2.** List of antibodies used, flow cytometry panels and flow-based definitions of lymphocyte populations.

| Laser | Filter | Mirror | Dye/Fluorochrome | InVivo labeling panel#1 | InVivo labeling panel#2 | InVivo labeling panel#3 |
| --- | --- | --- | --- | --- | --- | --- |
| UV | 379/28 |  | BUV395 | CD8 | CD8 | CD8 |
|  | 740/35 | 690LP | BUV737 | CD69 | a4b7 | PD-1 |
| Violet | 450/50 |  | BV421, V450 | CCR7 | CCR7 | Granzyme B |
|  | 525/50 | 505LP | LIVE/DEAD AQUA | LIVE/DEAD | LIVE/DEAD | LIVE/DEAD |
|  | 610/20 | 600LP | BV605 | CD95 | CD95 | CCR5 |
|  | 710/50 | 690LP | BV711 | CD45RA | Ki67 | CXCR3 |
|  | 780/60 | 750LP | BV785 | CD4 | CD4 | CD4 |
| Blue | 530/30 | 505LP | FITC | CD45-AF488 | CD45-AF488 | CD45-AF488 |
|  | 695/40 | 630LP | PerCP-Cy5.5 | CD14/CD20 | CD14/CD20 | CD14/CD20 |
| Yellow/Green | 586/15 |  | PE, AlexaFluor555 | MAMU*A001 | MAMU*A001 | MAMU*A001 |
|  | 780/60 | 750LP | PE-Cy7 | CD103 | CD45RA | CD137 |
| Red | 670/30 |  | APC, AlexaFluor647 | CD45-AF647 | CD45-AF647 | CD45-AF647 |
|  | 780/60 | 750LP | APC-Cy7 | CD3 | CD3 | CD3 |

Alternative flow panels for experiments where in vivo CD45-AF555 Abs were used:
| Laser | Filter | Mirror | Dye/Fluorochrome | InVivo labeling panel#1 | InVivo labeling panel#2 | InVivo labeling panel#3 |
| --- | --- | --- | --- | --- | --- | --- |
| UV | 379/28 |  | BUV395 | CD8 | CD8 | CD8 |
|  | 740/35 | 690LP | BUV737 | CD69 | a4b7 | PD-1 |
| Violet | 450/50 |  | BV421, V450 | CCR7 | CCR7 | Granzyme B |
|  | 525/50 | 505LP | LIVE/DEAD AQUA | LIVE/DEAD | LIVE/DEAD | LIVE/DEAD |
|  | 610/20 | 600LP | BV605 | CD95 | CD95 | CCR5 |
|  | 710/50 | 690LP | BV711 | CD45RA | Ki67 | CXCR3 |
|  | 780/60 | 750LP | BV785 | CD4 | CD4 | CD4 |
| Blue | 530/30 | 505LP | FITC | CD45-AF488 | CD45-AF488 | CD45-AF488 |
|  | 695/40 | 630LP | PerCP-Cy5.5 | CD14/CD20 | CD14/CD20 | CD14/CD20 |
| Yellow/Green | 586/15 |  | PE, AlexaFluor555 | CD45-AF555 | CD45-AF555 | CD45-AF555 |
|  | 780/60 | 750LP | PE-Cy7 | CD103 | CD45RA | CD137 |
| Red | 670/30 |  | APC, AlexaFluor647 | CD45-AF647 | CD45-AF647 | CD45-AF647 |
|  | 780/60 | 750LP | APC-Cy7 | CD3 | CD3 | CD3 |

**List of used antibodies**
| Antibody | Fluorochrome | Clone | Company | Catalogue # | Marker Type |
| --- | --- | --- | --- | --- | --- |
| CD3 | APC-Cy7 | SP34-2 | BD Biosciences | 557757 | Extracellular |
| CD4 | BV786 | L200 | BD Biosciences | 563914 | Extracellular |
| CD8 | BUV395 | RPA-T8 | BD Biosciences | 563795 | Extracellular |
| CD14 | PerCP-Cy5.5 | M5E2 | BD Biosciences | 550787 | Extracellular |
| CD20 | PerCP-Cy5.5 | 2H7 | eBioscience | 45-0209-42 | Extracellular |
| CD45 | Unconjugated* | MB4-6D6 | Miltenyi | Custom order | In vivo |
| CD45RA | PE-Cy7 | 2H4LDH11LDB9 | Beckman Coulter | B10821 | Extracellular |
| CD45RA | BV711 | 5H9 | BD Biosciences | 740806 | Extracellular |
| CD69 | BUV737 | FN50 | BD Biosciences | 612817 | Extracellular |
| CD95 | BrilliantViolet605 | DX2 | Biolegend | 305628 | Extracellular |
| CD103 | PE-Cy7 | B-Ly7 | eBioscience | 25-1038-42 | Extracellular |
| CD183 (CXCR3) | BV711 | 1C6 | BD Biosciences | 563156 | Extracellular |
| CD195 (CCR5) | BV605 | 3A9 | BD Biosciences | 742912 | Extracellular |
| CD197 (CCR7) | BrilliantViolet421 | G043H7 | Biolegend | 353208 | Extracellular |
| CD297 (PD-1) | BUV737 | EH12.1 | BD Biosciences | 612791 | Extracellular |
| a4b7 | Biotin | ACT1 | HIV AIDS Reagent Program | 11718 | Extracellular |
| Granzyme B | BV421 | GB11 | BD Biosciences | 563389 | Intracellular |
| Ki67 | BV711 | B56 | BD Biosciences | 563755 | Intracellular |
| MAMU*A001 | PE | N/A | NHP Resorce Reagent | PR-1102 | Extracellular |

\*Custom conjugation with AlexaFluor488, AlexaFluor555, AlexaFluor647, Biotin

| Phenotypic definition of T cell subsets: |  |
| --- | --- |
| T cells | CD3+CD14-CD20- lymphocytes |
| CD4 T cells | CD4+CD8-CD3+CD14-CD20- lymphocytes |
| CD8 T cells | CD4-CD8+CD3+CD14-CD20- lymphocytes |
| Compartment-1 CD4 T cells | CD45AF488+CD4+CD8-CD3+CD14-CD20- lymphocytes |
| Compartment-1 CD8 T cells | CD45AF488+CD4-CD8+CD3+CD14-CD20- lymphocytes |
| Compartment-2 CD4 T cells | CD45AF647+CD45AF488-CD4+CD8-CD3+CD14-CD20- lymphocytes |
| Compartment-2 CD8 T cells | CD45AF647+CD45AF488-CD4-CD8+CD3+CD14-CD20- lymphocytes |
| Compartment-3 CD4 T cells | CD45AF647-CD45AF488-CD4+CD8-CD3+CD14-CD20- lymphocytes |
| Compartment-3 CD8 T cells | CD45AF647-CD45AF488-CD4-CD8+CD3+CD14-CD20- lymphocytes |
| Naïve T cells (Tn) | CD45RA+CCR7+CD95- CD4 or CD8 T cells |
| T memory stem cells (Tscm) | CD45RA+CCR7+CD95+ CD4 or CD8 T cells |
| Central memory T cells | CD45RA-CCR7+ CD4 or CD8 T cells |
| Effector memory T cell (Tem) | CD45RA-CCR7- CD4 or CD8 T cells |
| Effector memory T cell with CD45RA re-expression (Temra) | CD45RA+CCR7- CD4 or CD8 T cells |

**Table S3.** scRNA-Seq sample information and QC metrics.

| Sample ID | Animal ID | Experimental cohort | Type of sample | Sequencing Batch | Estimated Number of Cells | Mean Reads per Cell | Median Genes per Cell | Number of Reads (x10<sup>6</sup>) | Valid Barcodes | Sequencing Saturation | Q30 Bases in Barcode | Q30 Bases in RNA Read | Q30 Bases in Sample Index | Q30 Bases in UMI | Reads Mapped to Genome | Reads Mapped Confidently to Genome | Reads Mapped Confidently to Intergenic Regions | Reads Mapped Confidently to Intronic Regions | Reads Mapped Confidently to Exonic Regions | Reads Mapped Confidently to Transcriptome | Reads Mapped Antisense to Gene | Fraction Reads in Cells | Total Genes Detected | Median UMI Counts per Cell | Mean Genes Per Cell | Cells Retained After QC and Filtering | 10X Genomics Chromium 3' Chemistry | Sequencer |
| --- | --- | --- | --- | --- | --- | --- | --- | --- | --- | --- | --- | --- | --- | --- | --- | --- | --- | --- | --- | --- | --- | --- | --- | --- | --- | --- | --- | --- |
| NHP_40_Host | R.276 | GVHD | FACS-purified Host cells | 1 | 5021 | 23337 | 707 | 117 | 0.97 | 0.64 | 0.92 | 0.88 | 0.91 | 0.92 | 0.92 | 0.69 | 0.09 | 0.179 | 0.422 | 0.385 | 0.008 | 0.791 | 17063 | 1815 | 748.5 | 2177 | V2 | NovaSeq S2 |
| NHP_41_Donor | R.276 | GVHD | FACS-purified Donor cells | 1 | 3191 | 32552 | 1103 | 104 | 0.97 | 0.62 | 0.93 | 0.89 | 0.88 | 0.92 | 0.92 | 0.81 | 0.098 | 0.234 | 0.478 | 0.431 | 0.017 | 0.88 | 16056 | 3298 | 1206.9 | 2388 | V2 | NovaSeq S2 |
| NHP_42_HC | R.253 | HC | FACS-purified healthy control cells | 1 | 5903 | 21270 | 707 | 126 | 0.98 | 0.64 | 0.93 | 0.88 | 0.91 | 0.92 | 0.93 | 0.81 | 0.101 | 0.16 | 0.552 | 0.506 | 0.017 | 0.804 | 15910 | 2136 | 843.7 | 5159 | V2 | NovaSeq S2 |
| NHP_45_Donor | R.227 | GVHD | FACS-purified Donor cells | 1 | 1277 | 106207 | 967 | 136 | 0.98 | 0.86 | 0.93 | 0.88 | 0.90 | 0.92 | 0.93 | 0.78 | 0.091 | 0.087 | 0.597 | 0.531 | 0.01 | 0.944 | 13889 | 3418 | 997.9 | 887 | V2 | NovaSeq S2 |
| NHP_60_HC | R.265 | HC | FACS-purified healthy control cells | 2 | 3647 | 28294 | 904 | 103 | 0.97 | 0.56 | 0.94 | 0.90 | 0.89 | 0.93 | 0.94 | 0.82 | 0.092 | 0.319 | 0.409 | 0.342 | 0.011 | 0.882 | 17891 | 2330 | 941.3 | 1379 | V3 | NovaSeq S2 |
| NHP_61_HC | R.241 | HC | FACS-purified healthy control cells | 2 | 4642 | 22103 | 868 | 103 | 0.96 | 0.48 | 0.94 | 0.90 | 0.91 | 0.93 | 0.95 | 0.86 | 0.118 | 0.384 | 0.359 | 0.308 | 0.021 | 0.824 | 17690 | 2285 | 980.3 | 3780 | V3 | NovaSeq S2 |
| NHP_63_HC | R.216 | HC | FACS-purified healthy control cells | 2 | 5064 | 34667 | 871 | 176 | 0.96 | 0.63 | 0.93 | 0.91 | 0.89 | 0.93 | 0.94 | 0.83 | 0.1 | 0.352 | 0.381 | 0.321 | 0.014 | 0.841 | 18739 | 2166 | 974.0 | 2633 | V3 | NovaSeq S2 |
| NHP_64_GVHD | R.312 | GVHD | FACS-purified GVHD cells* | 2 | 3769 | 46784 | 1203 | 176 | 0.97 | 0.66 | 0.94 | 0.91 | 0.90 | 0.93 | 0.95 | 0.85 | 0.112 | 0.276 | 0.459 | 0.387 | 0.015 | 0.887 | 18122 | 3286 | 1232.4 | 2265 | V3 | NovaSeq S2 |
| NHP_65_GVHD | R.255 | GVHD | FACS-purified GVHD cells* | 2 | 2198 | 25218 | 932 | 55 | 0.96 | 0.39 | 0.93 | 0.90 | 0.89 | 0.93 | 0.95 | 0.82 | 0.107 | 0.24 | 0.475 | 0.39 | 0.014 | 0.859 | 15708 | 2271 | 1014.6 | 822 | V3 | NovaSeq S2 |

**Table S4.**
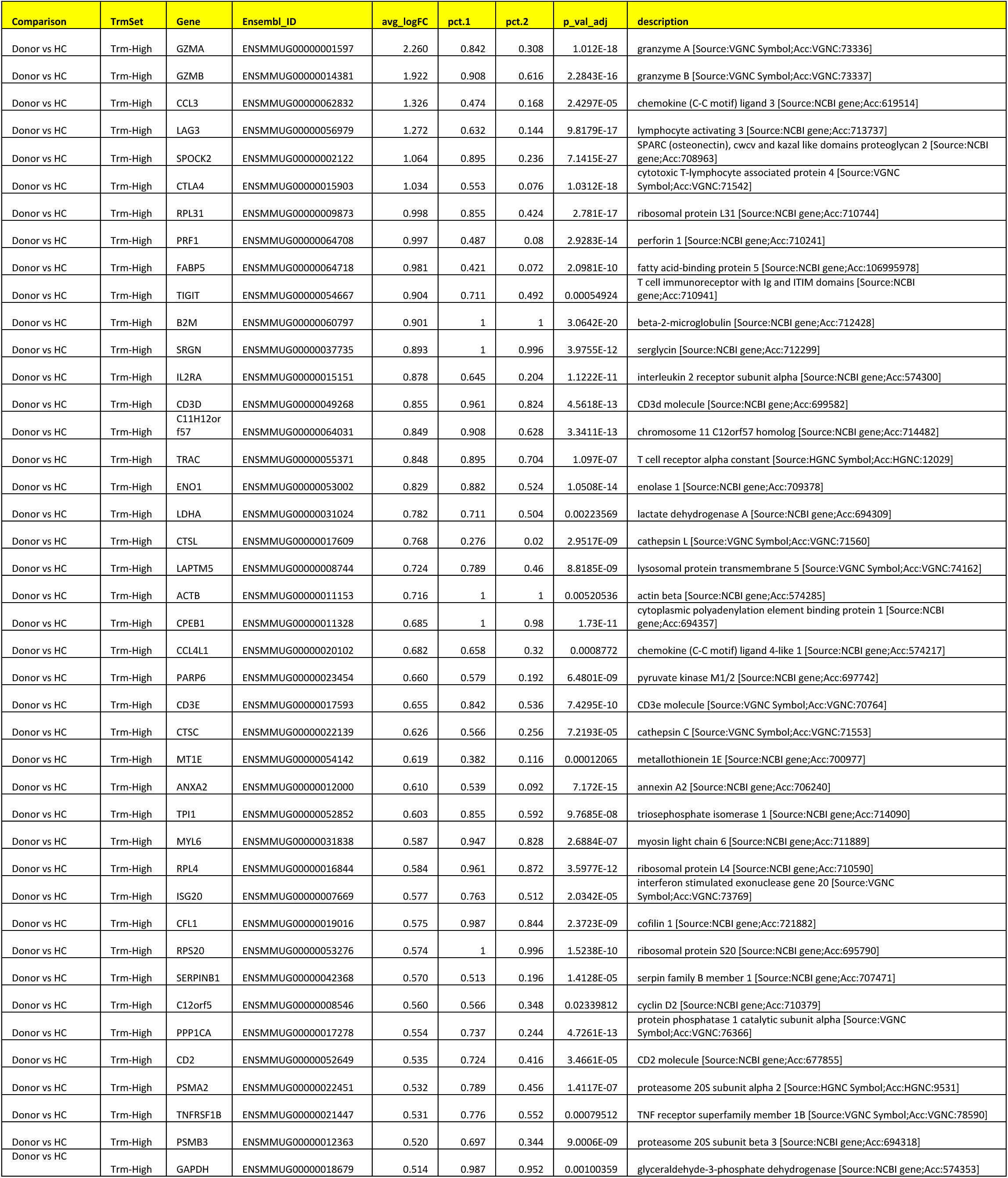
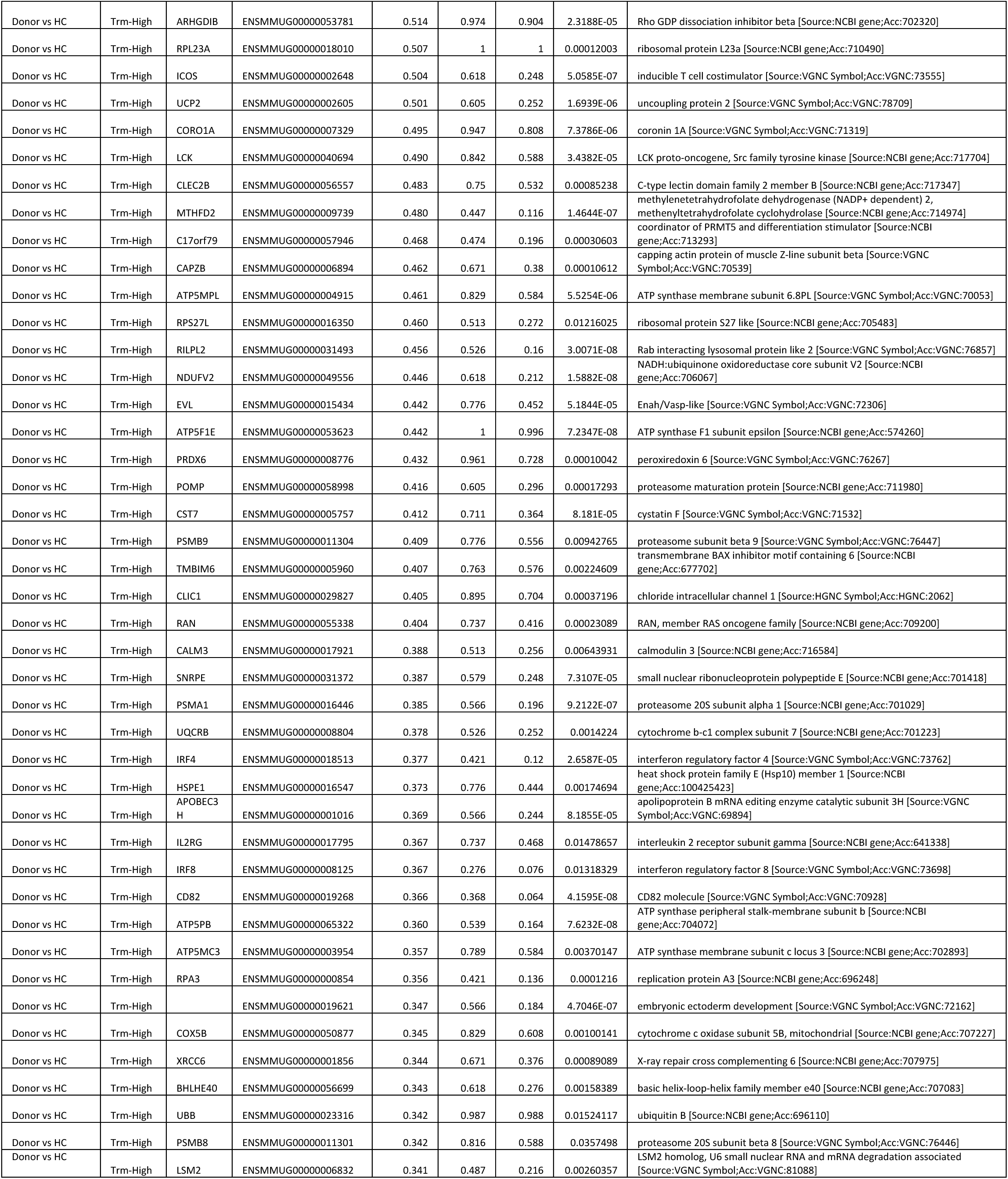
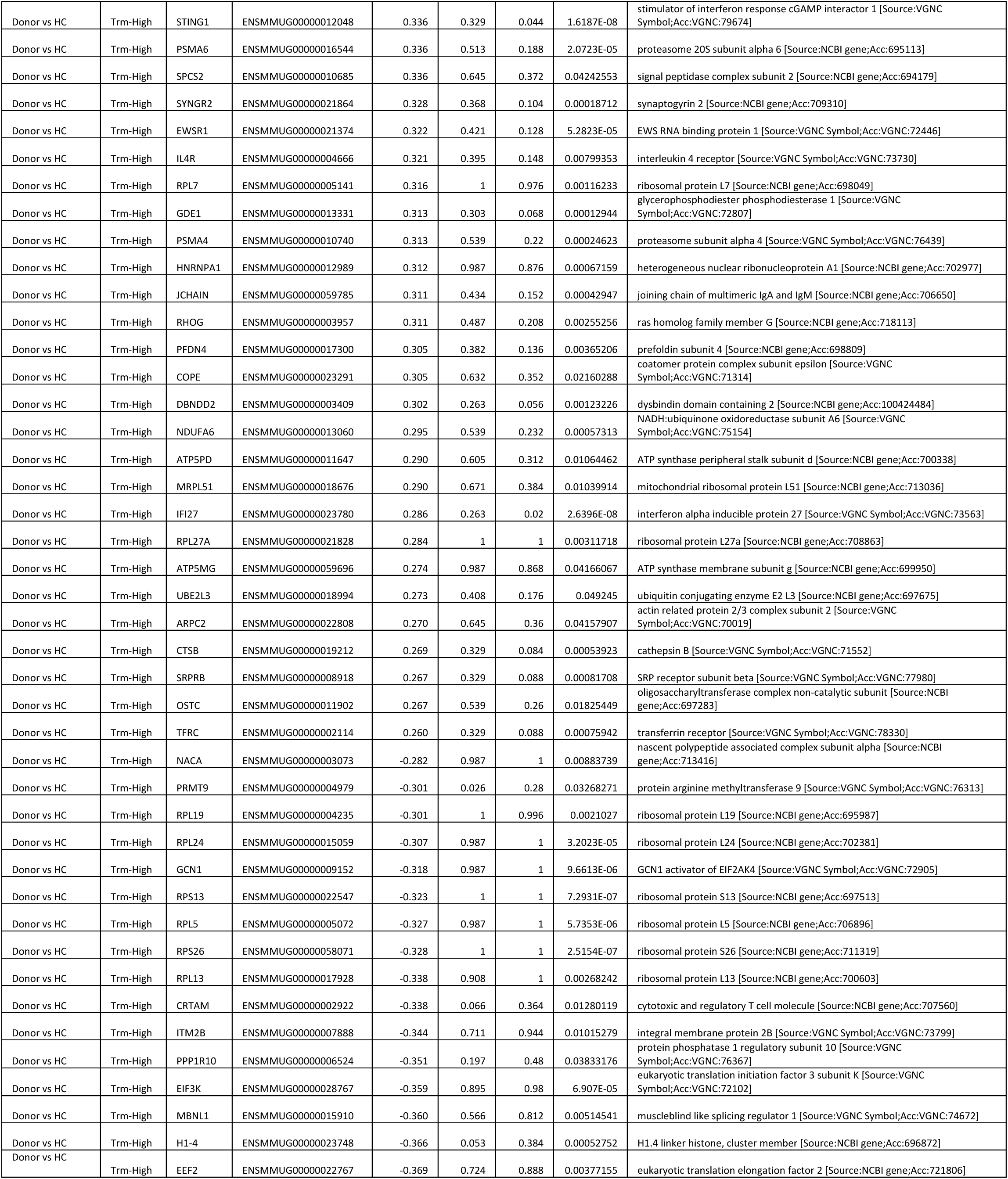
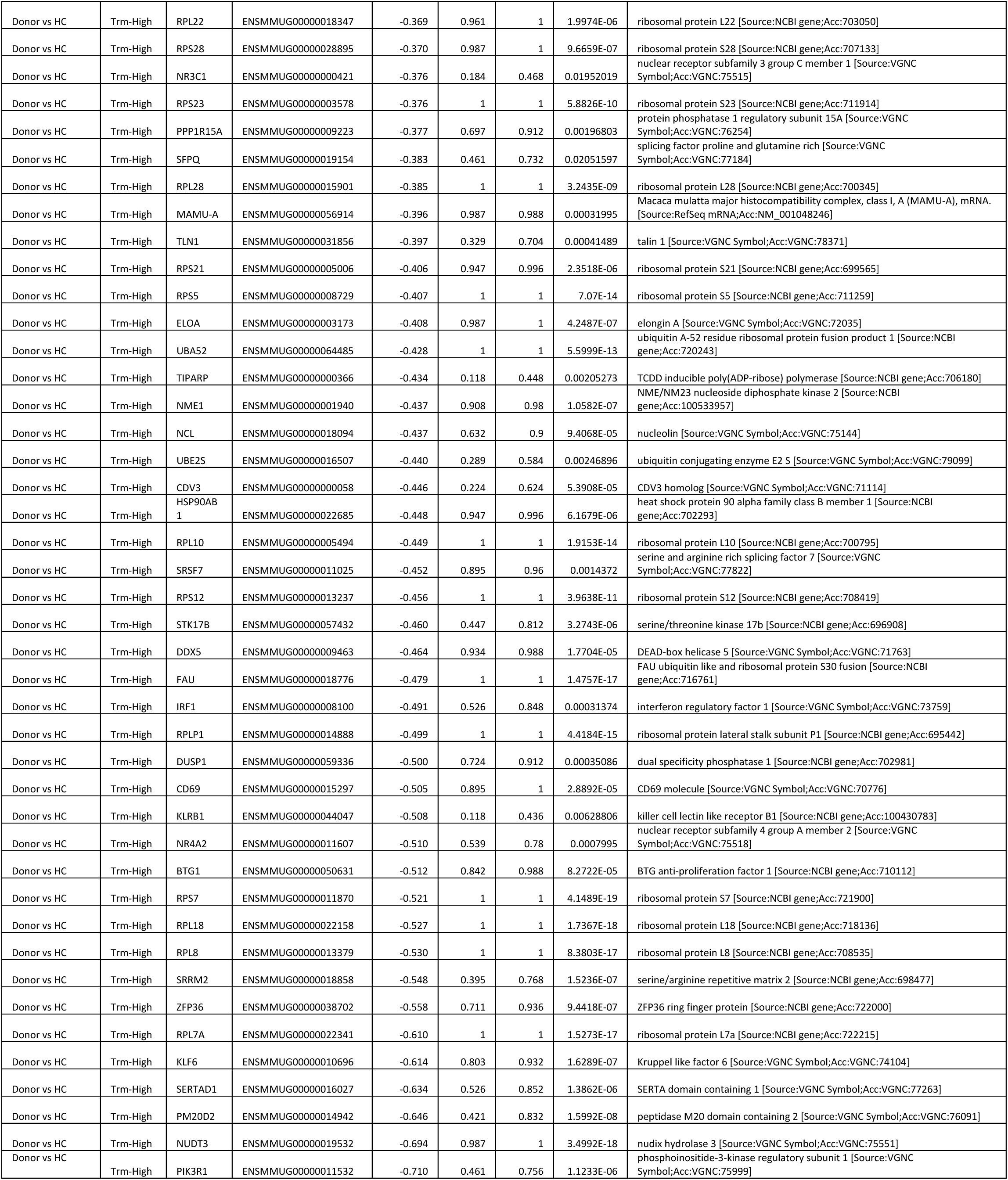
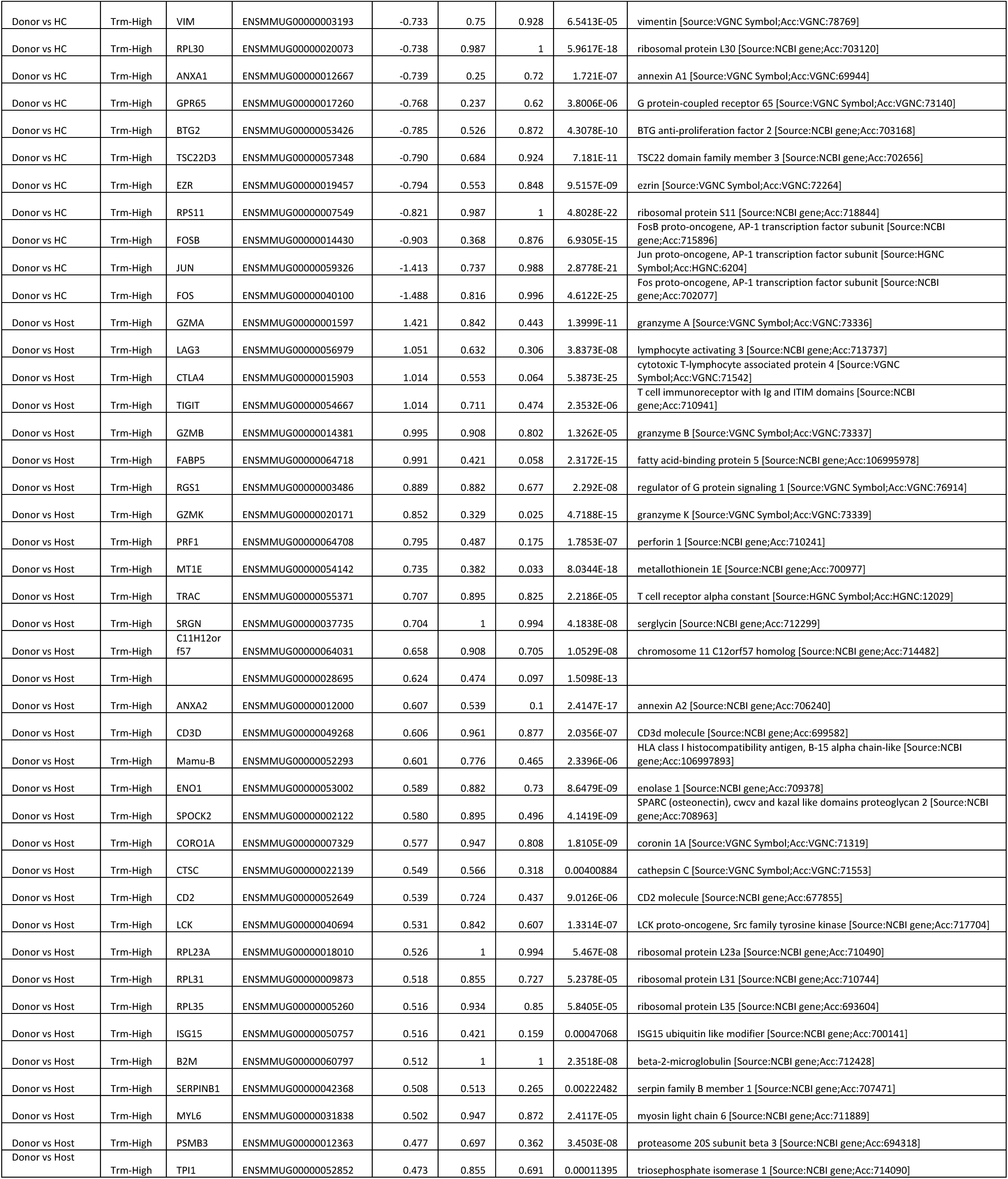
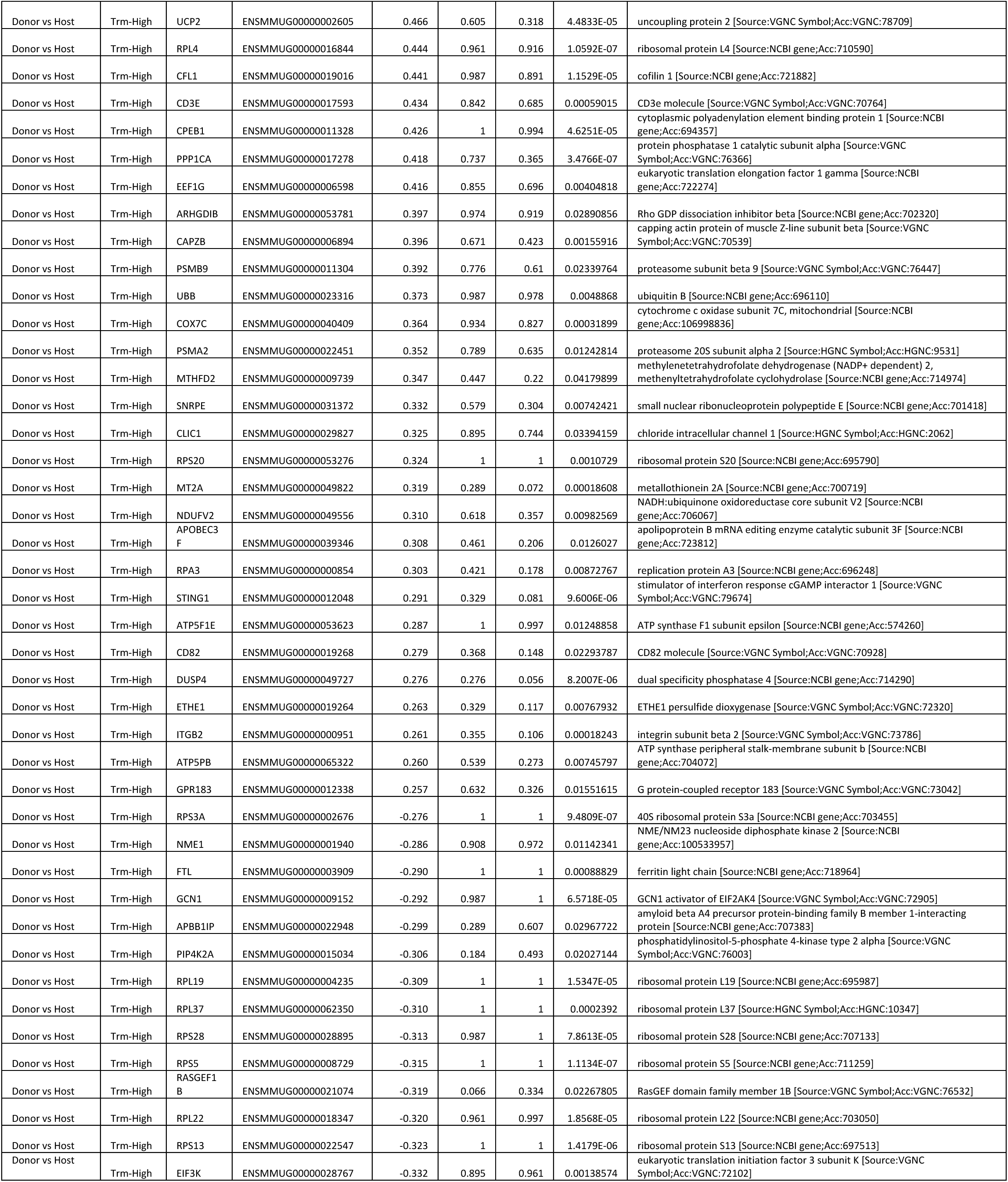
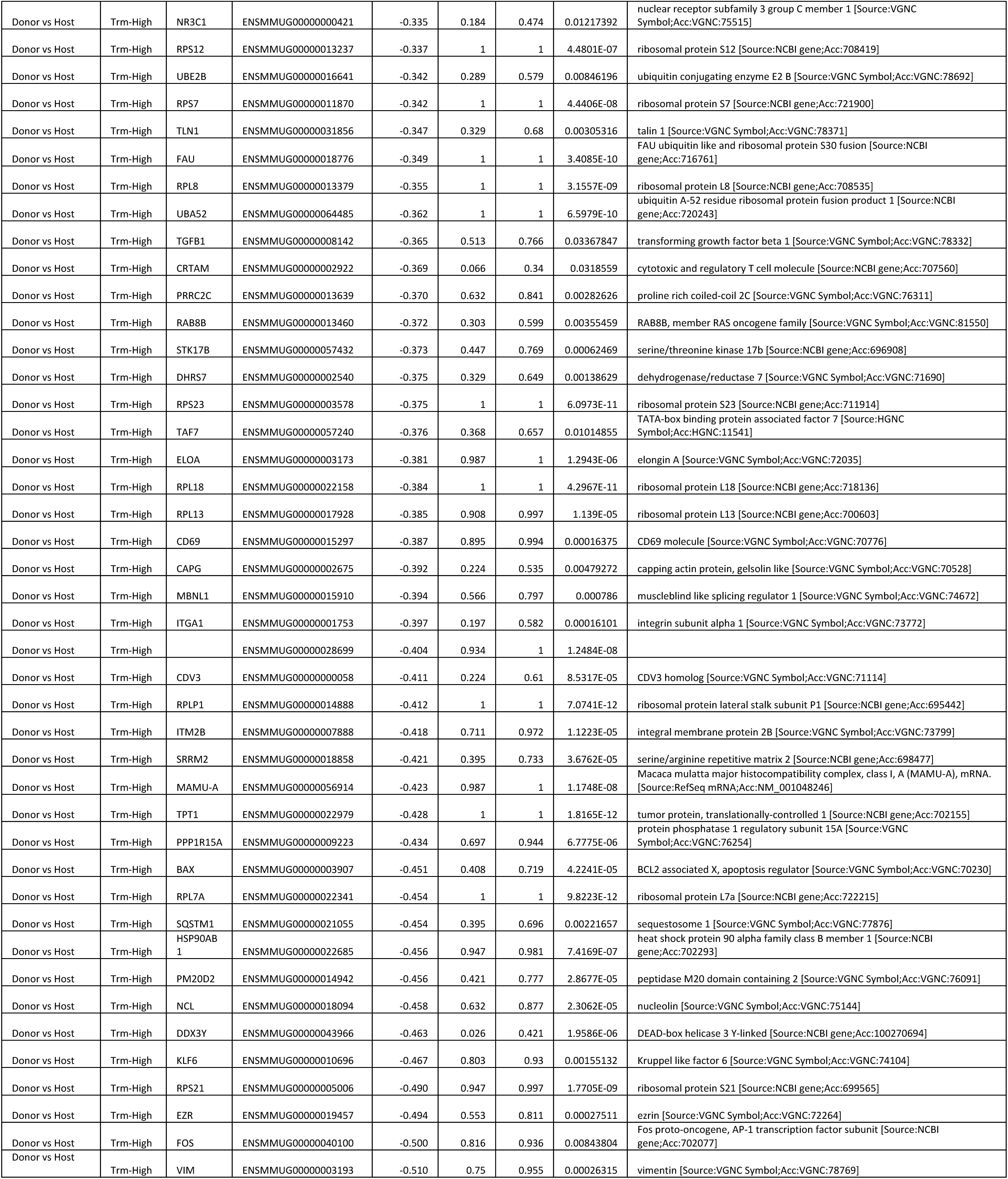
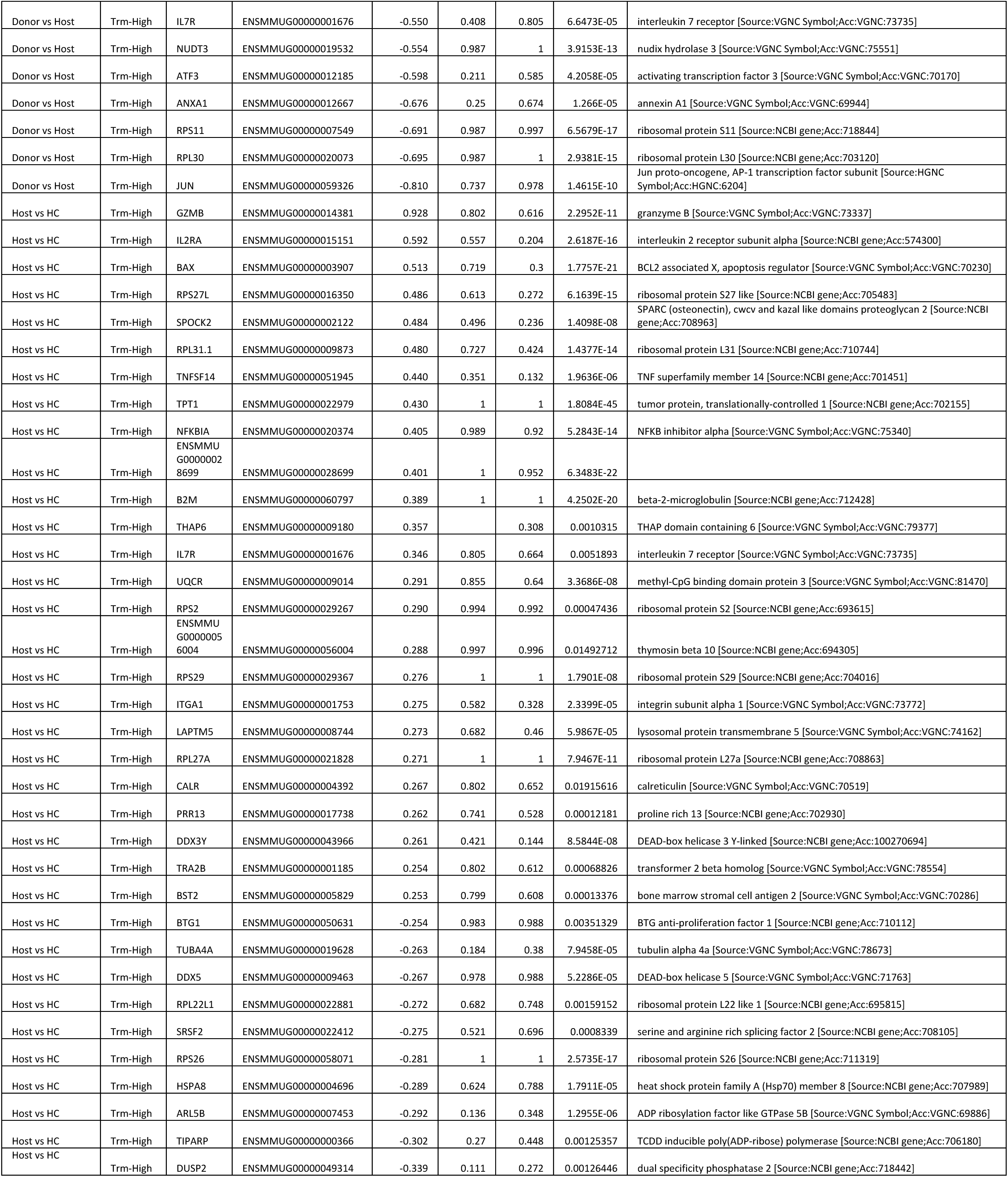
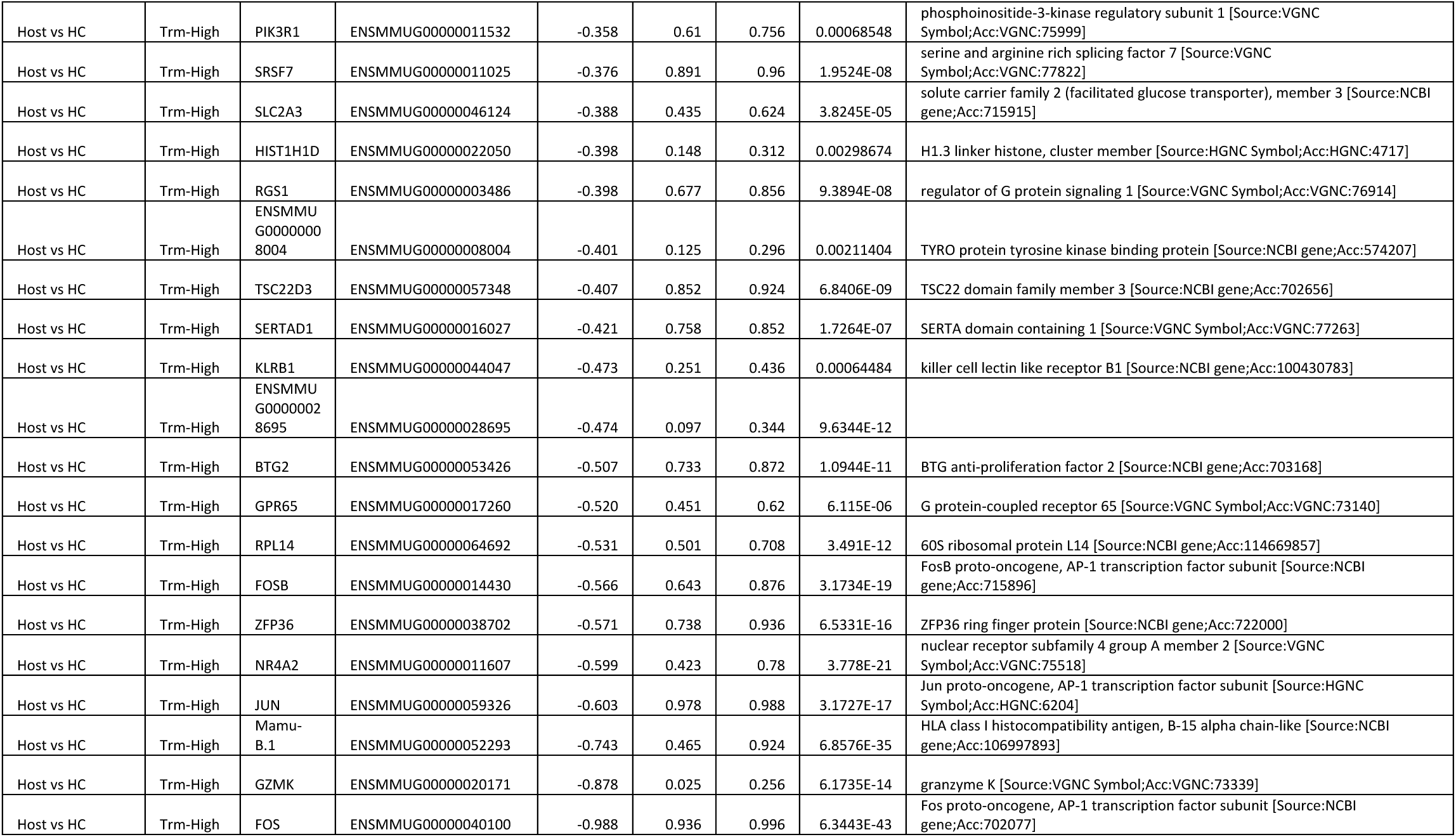
Differentially expressed (DE) genes between donor, host and healthy control CD8 T cells with high

**Table S5.**
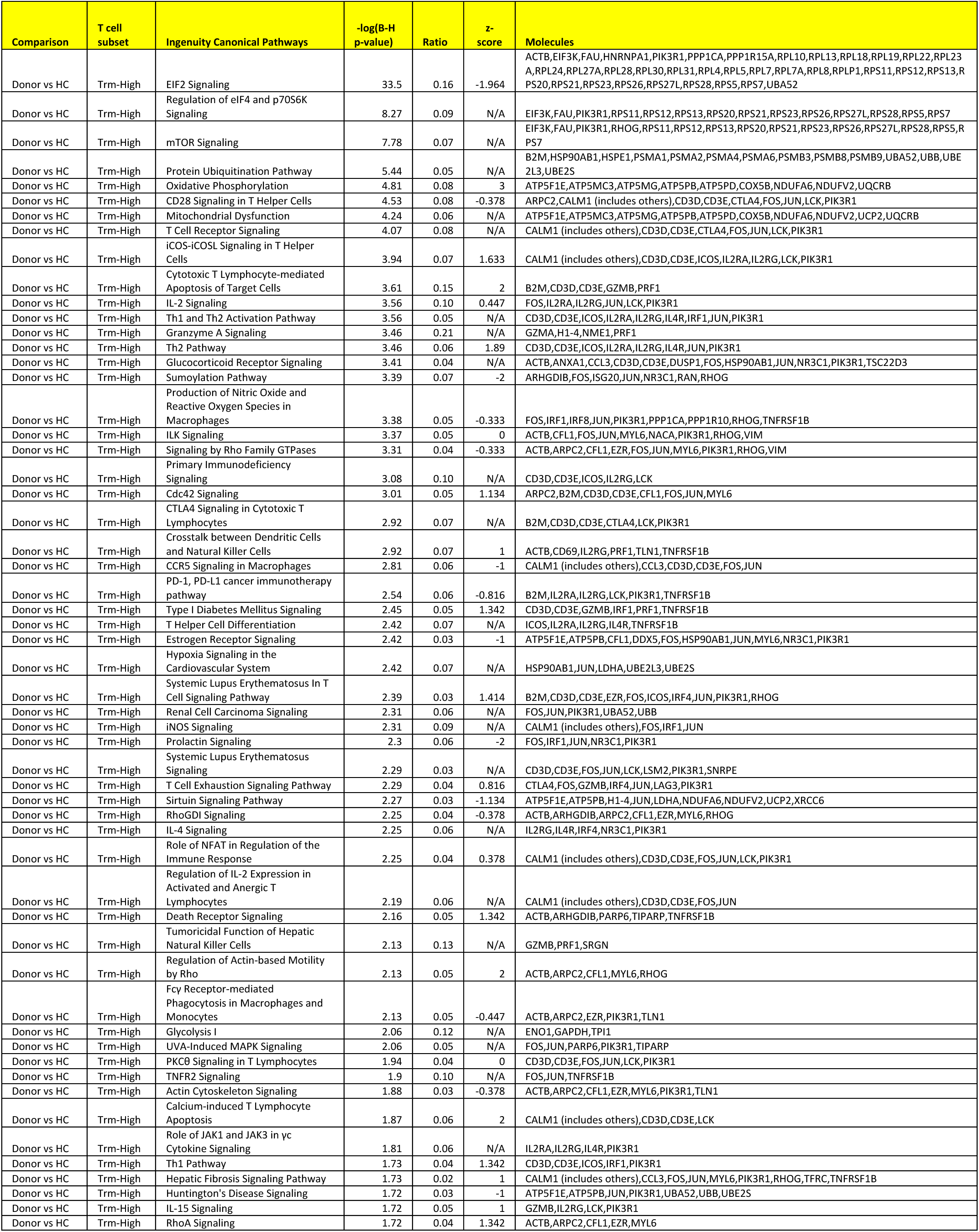
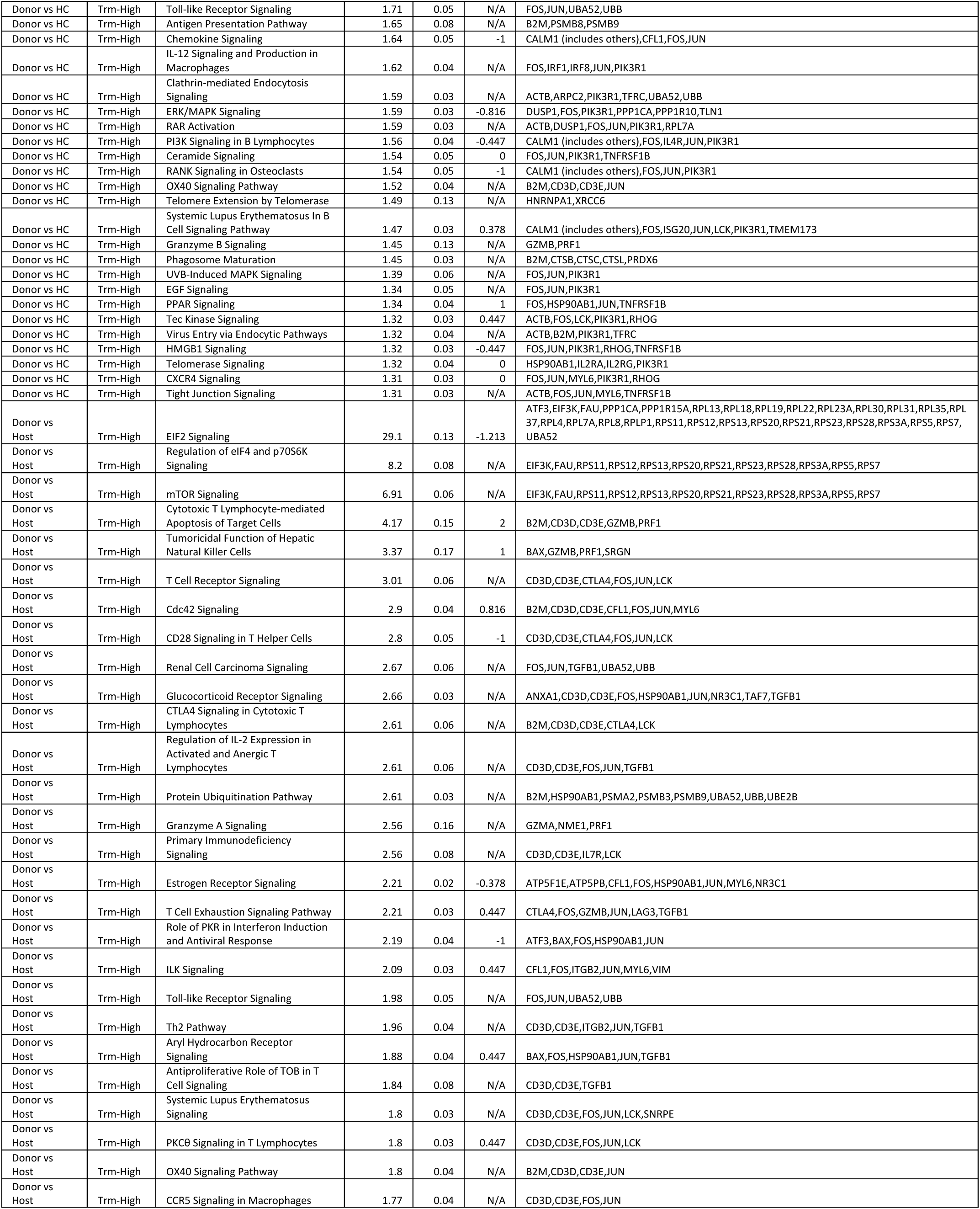
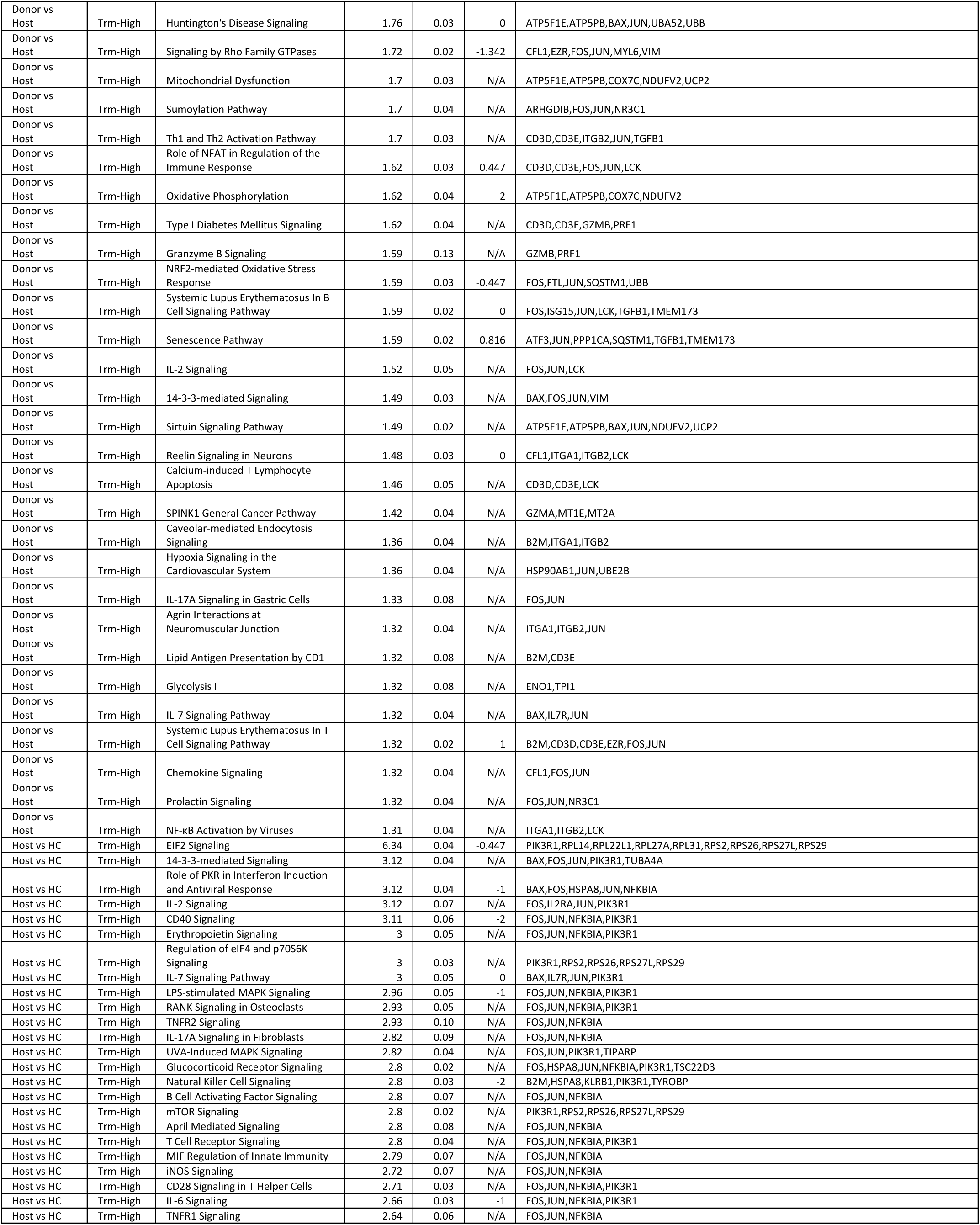
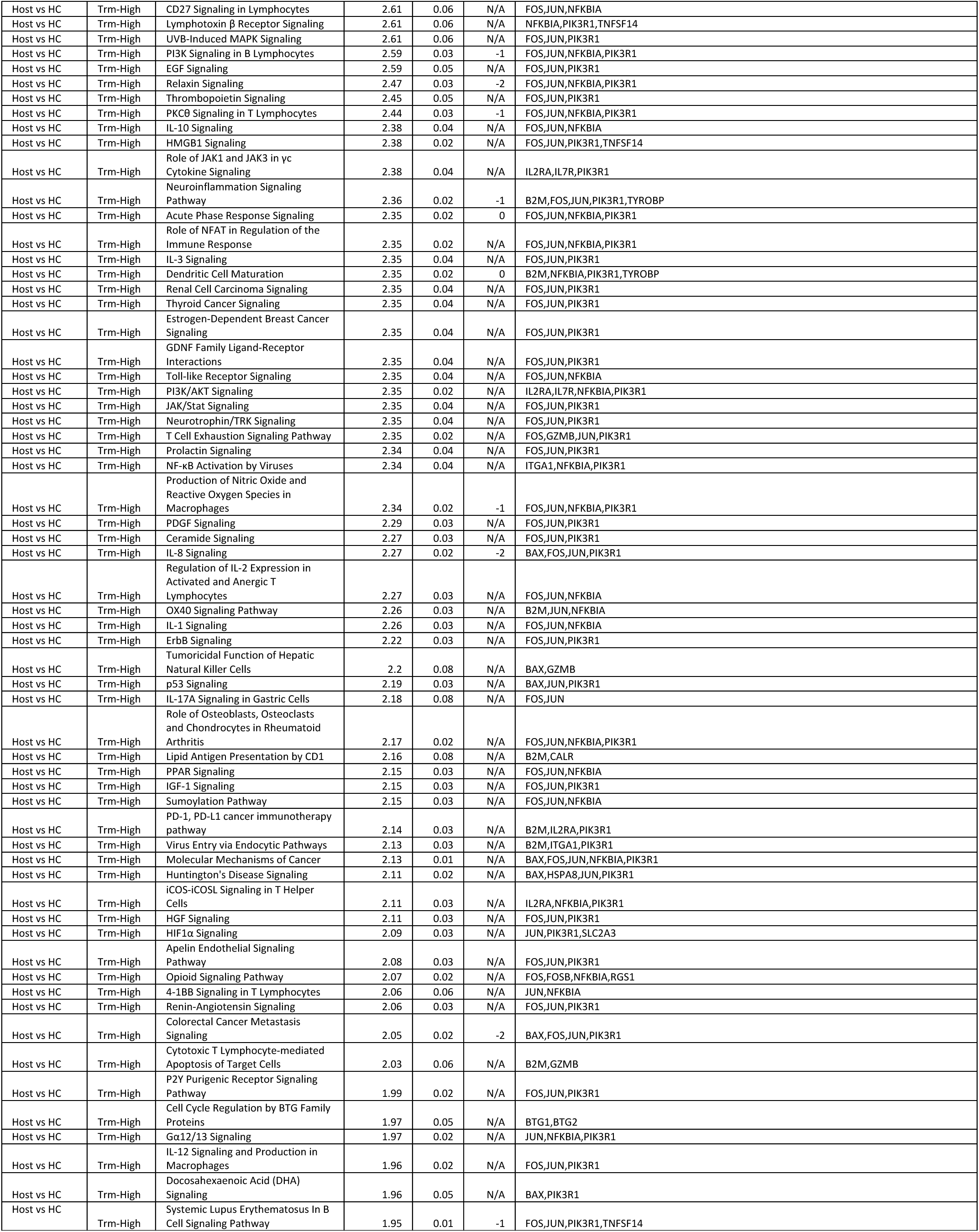
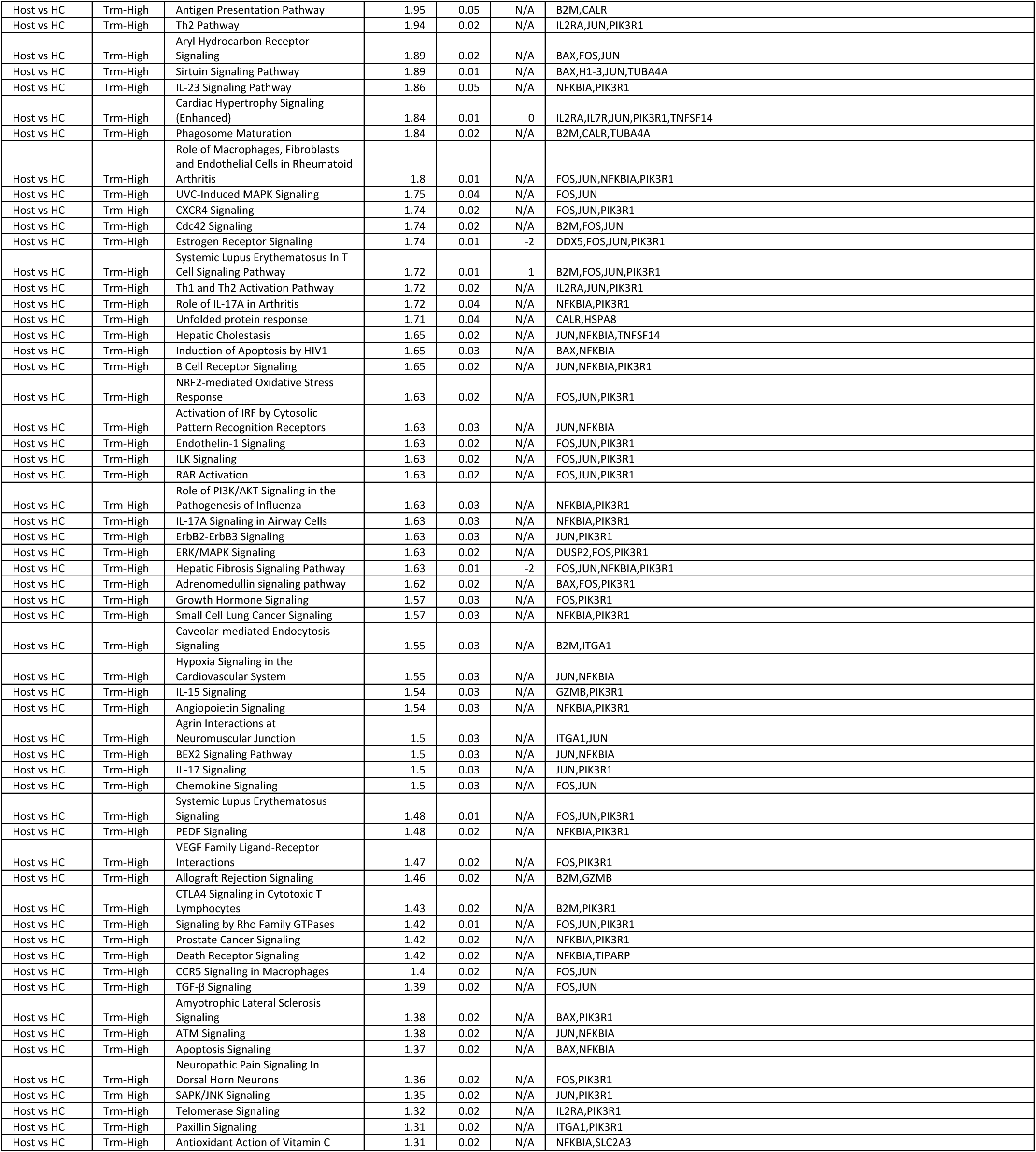
Pathway analysis on DE genes between donor, host and healthy control CD8 T cells with high enrichment score for T_RM_ signature.

**Table S6.**
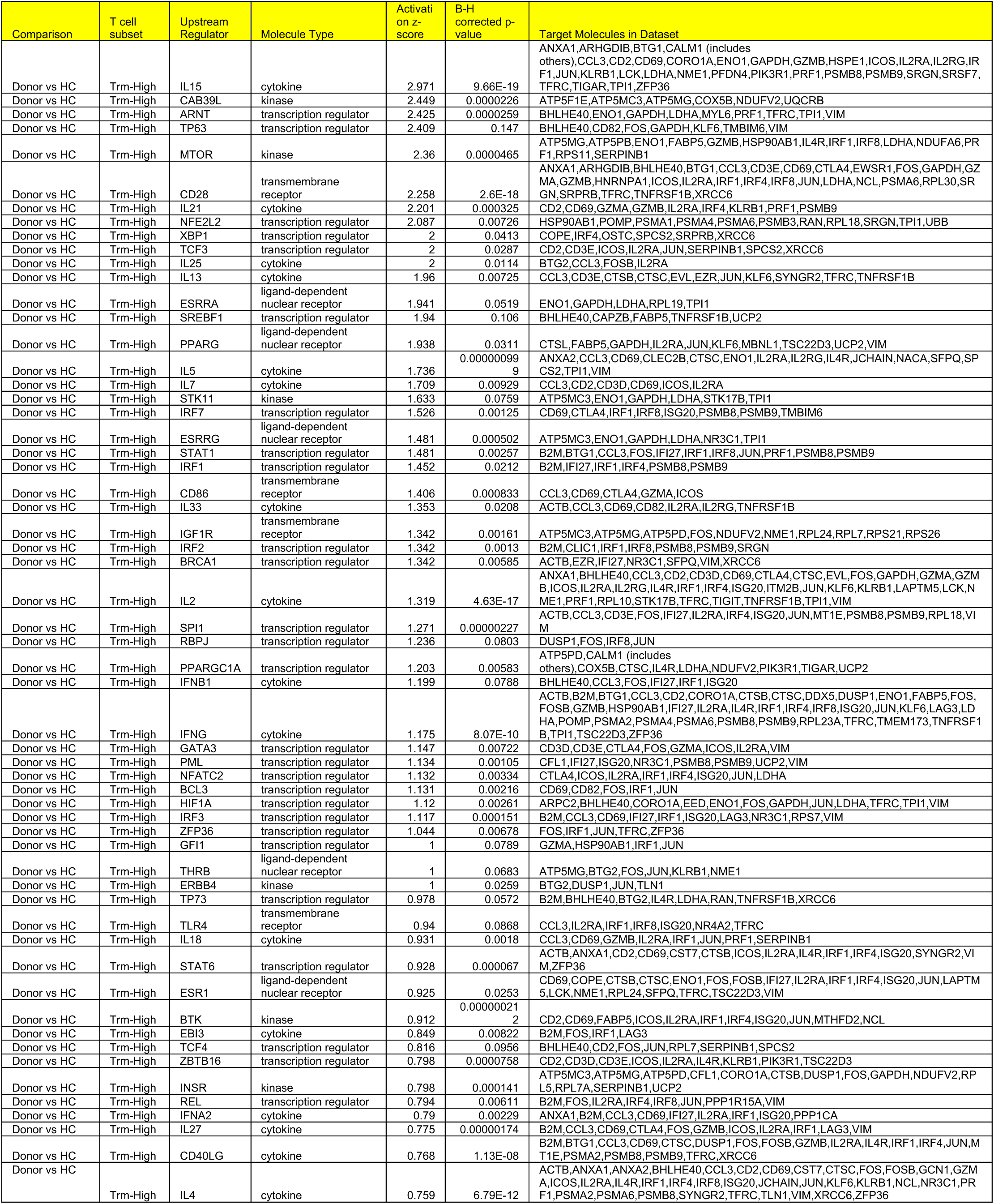
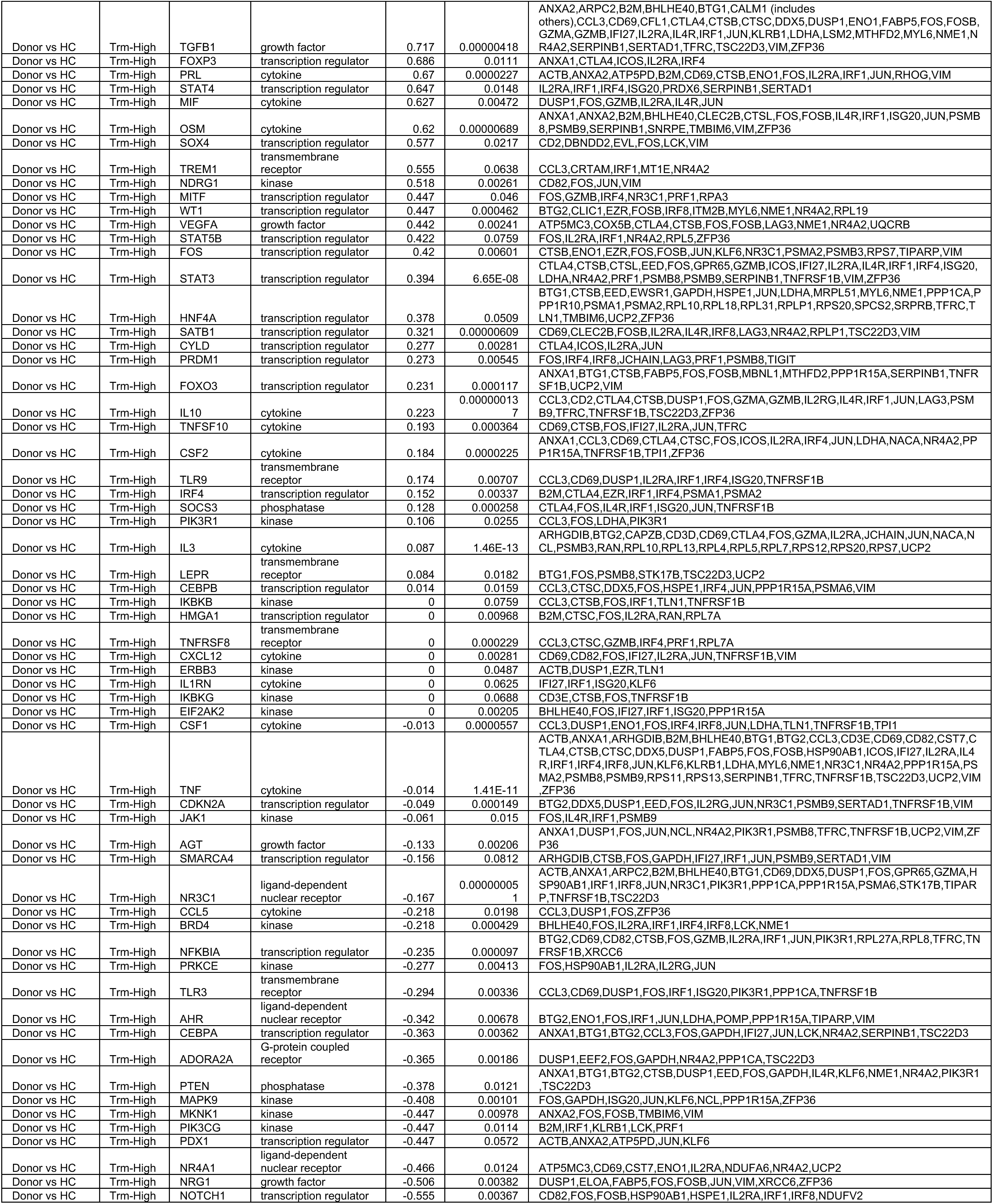
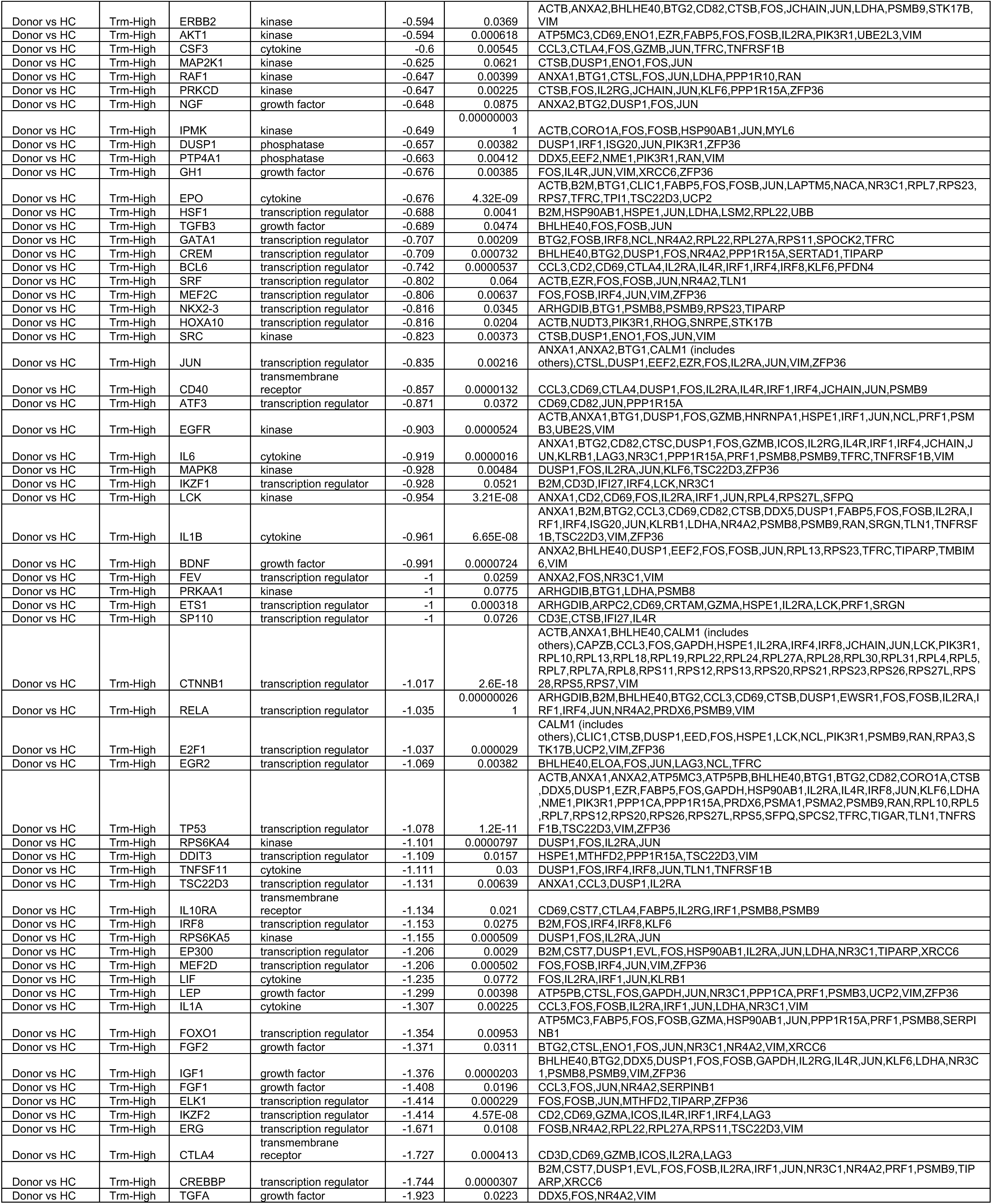
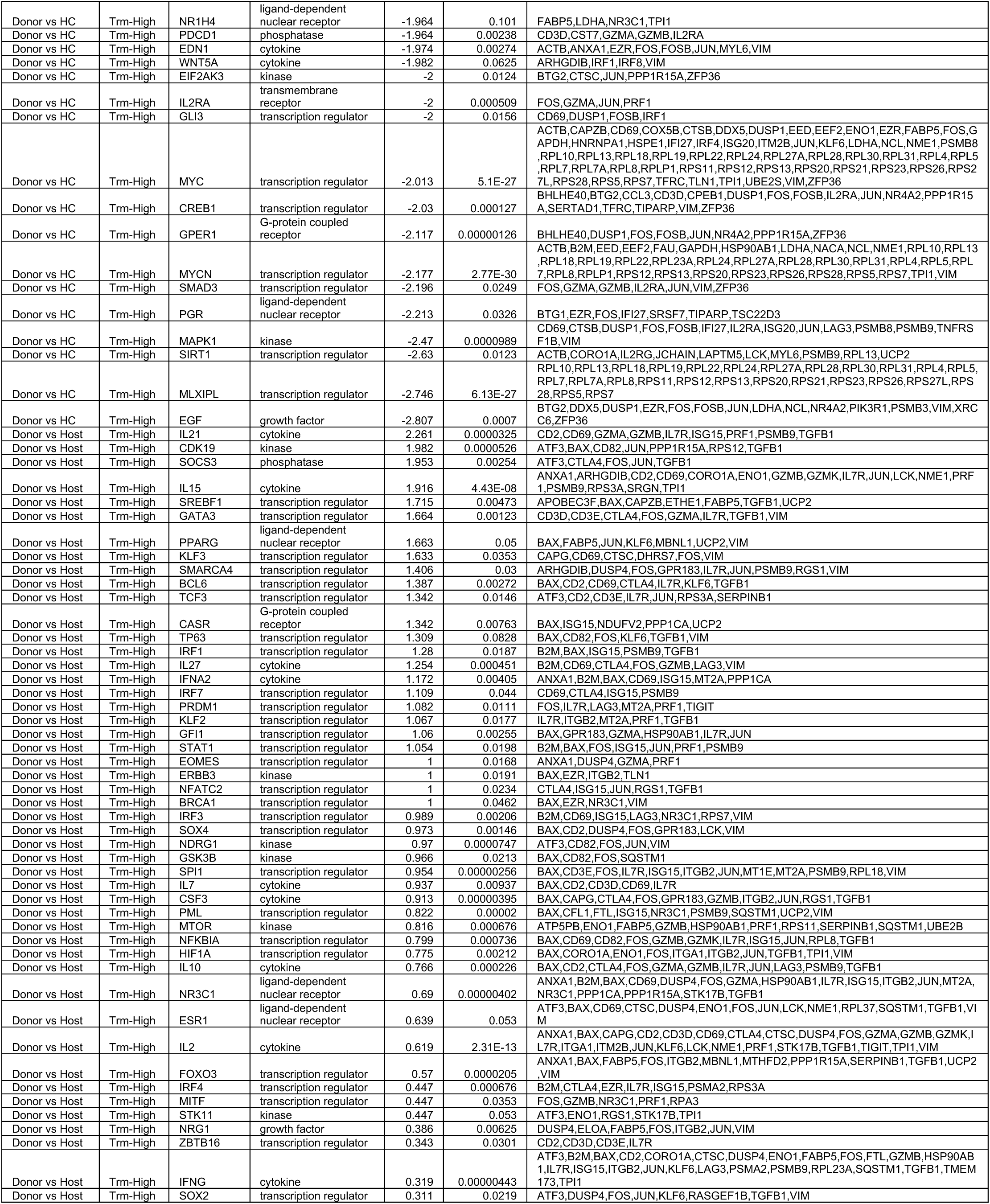
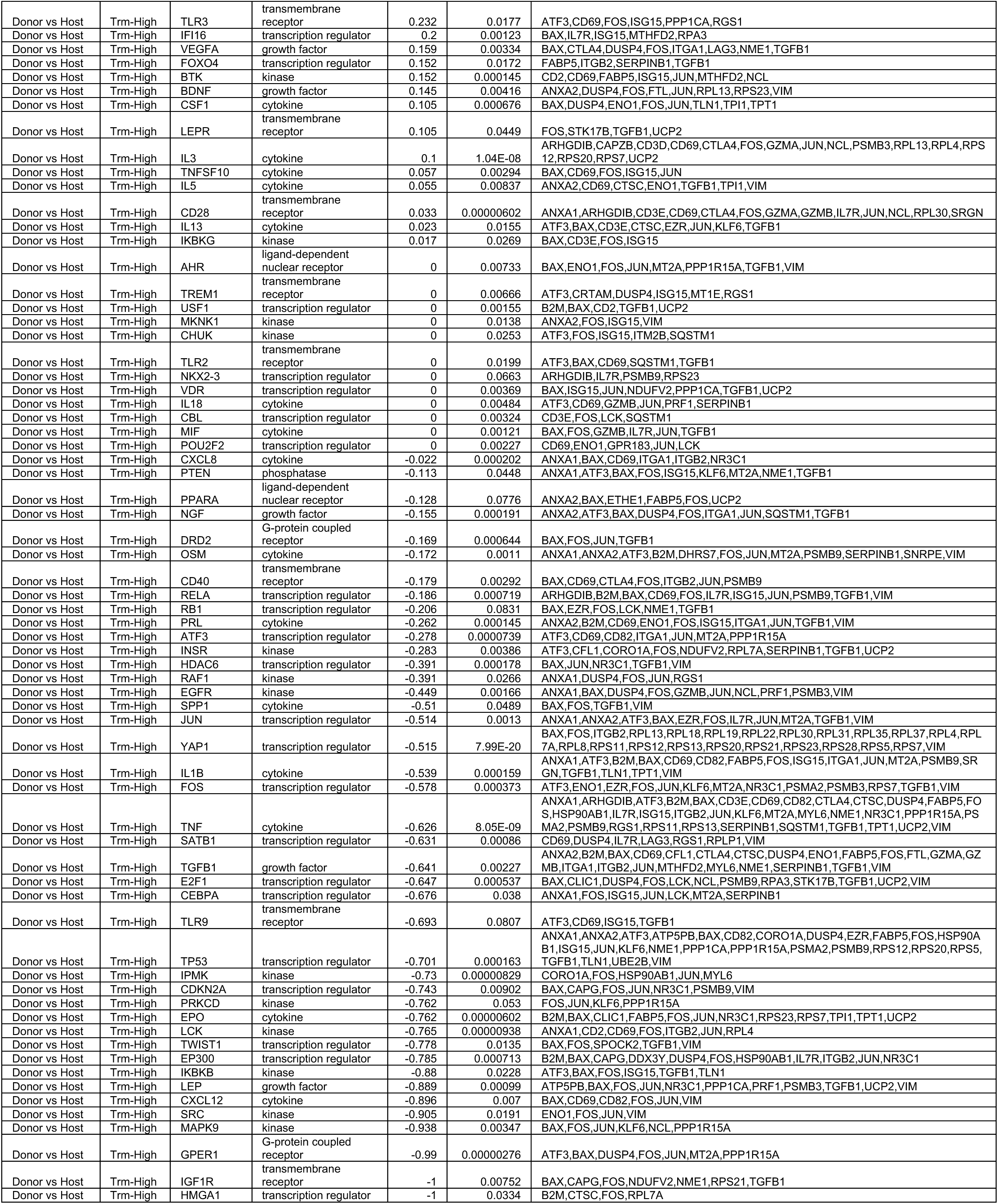
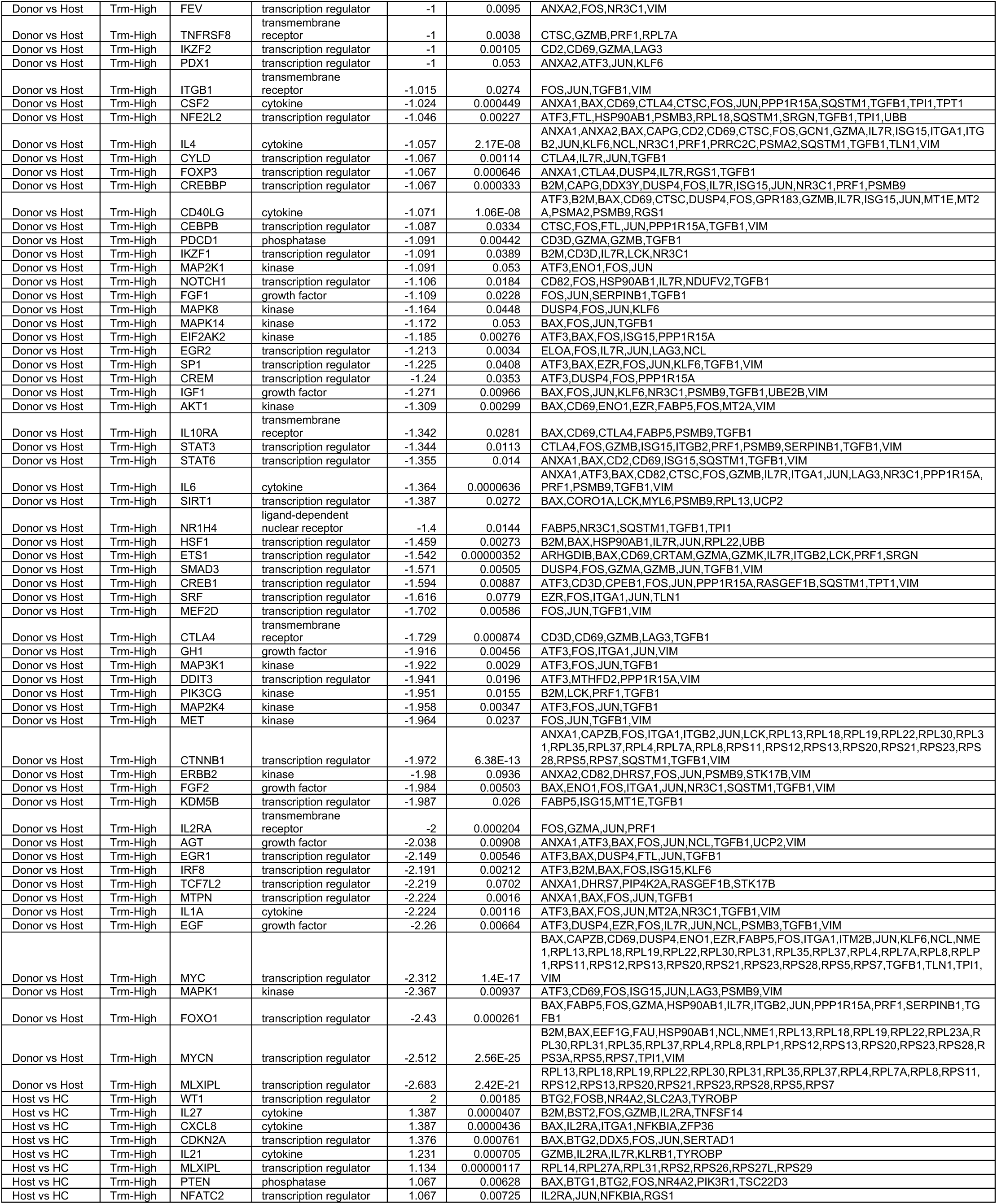
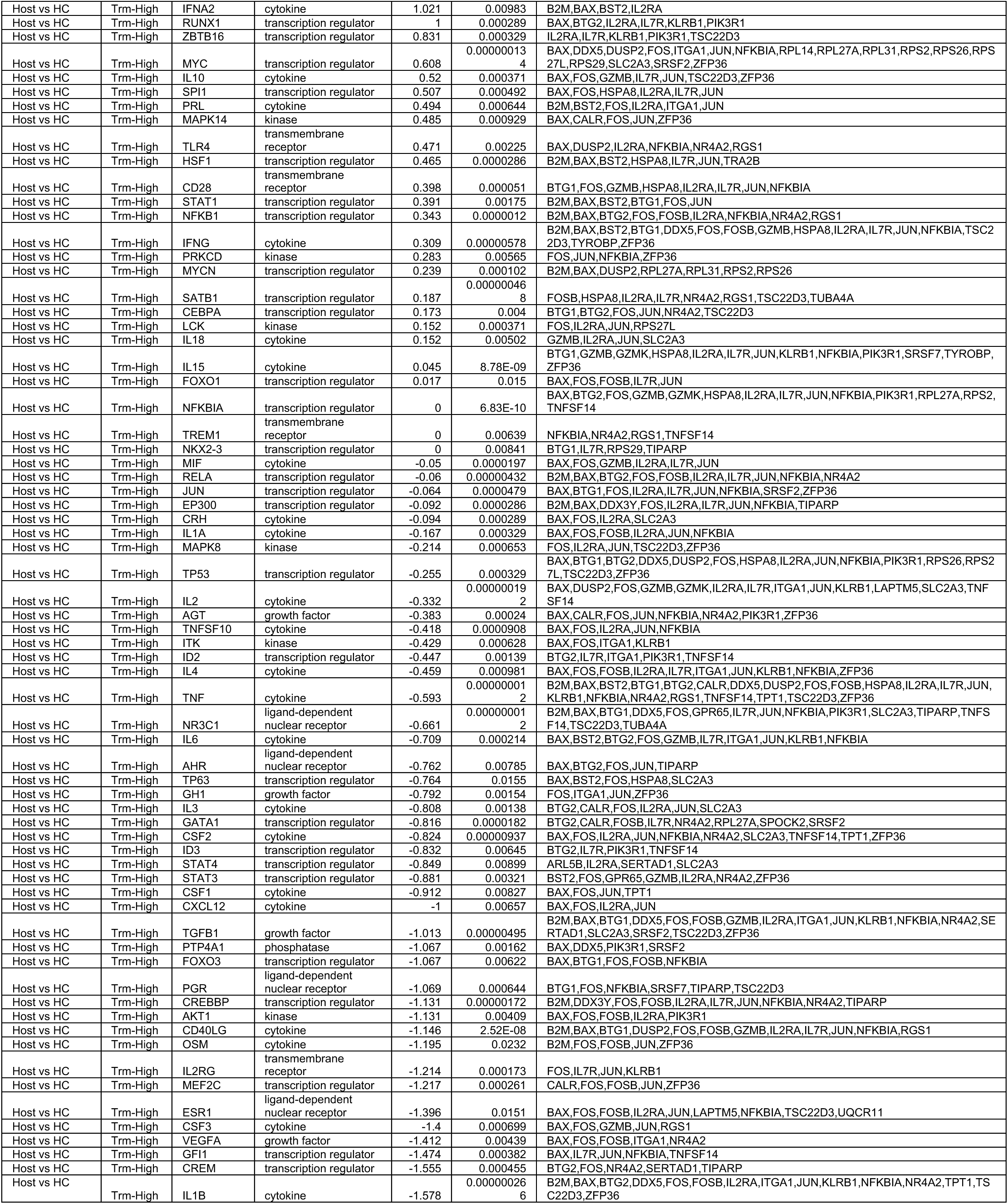
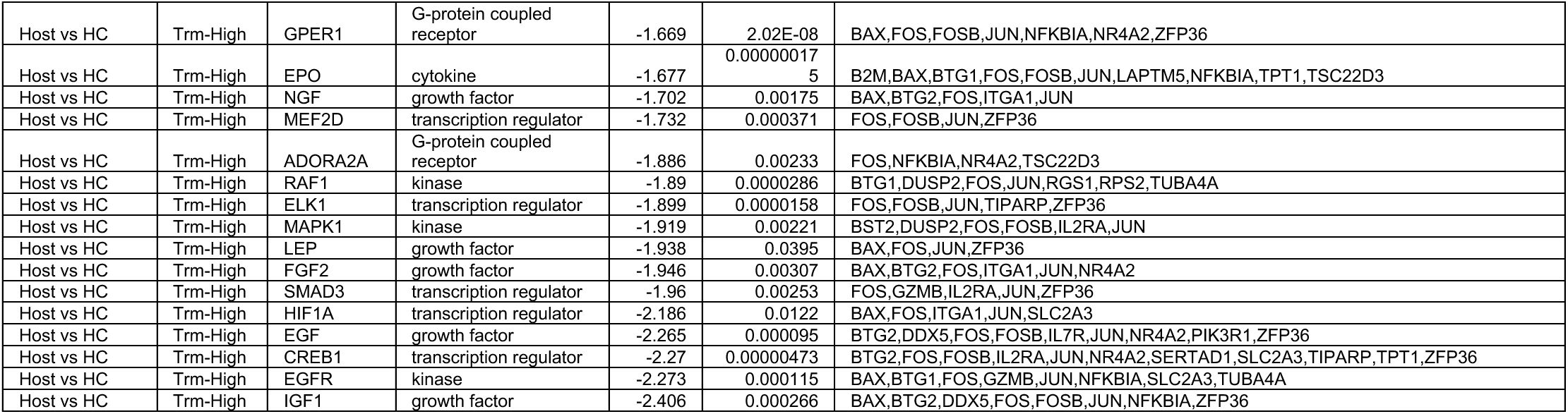
Predicted activated up-stream regulators in donor CD8 T cells with high enrichment score for T_RM_ signature in comparison with their healthy control and host counterparts.

**Table S7.**
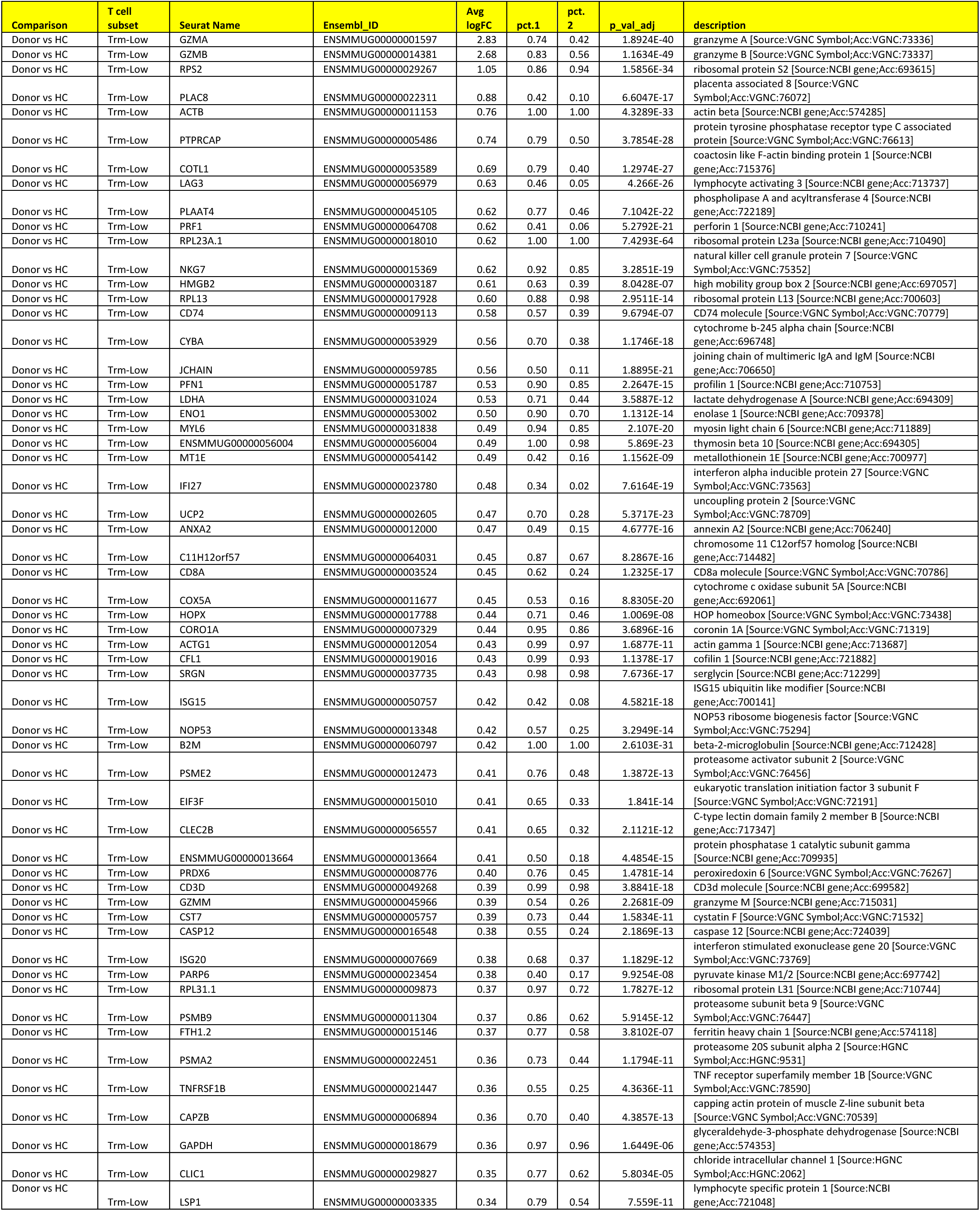
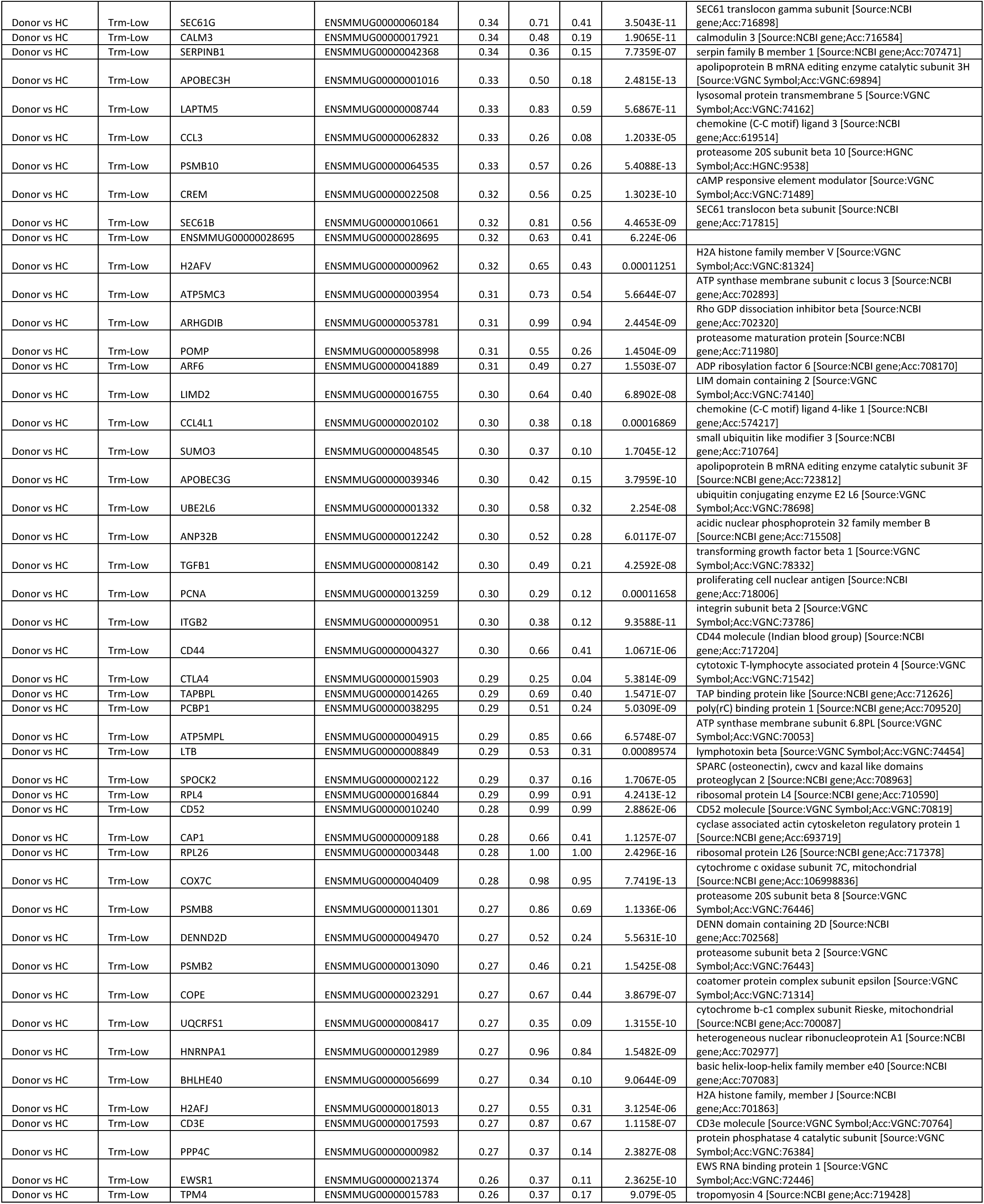
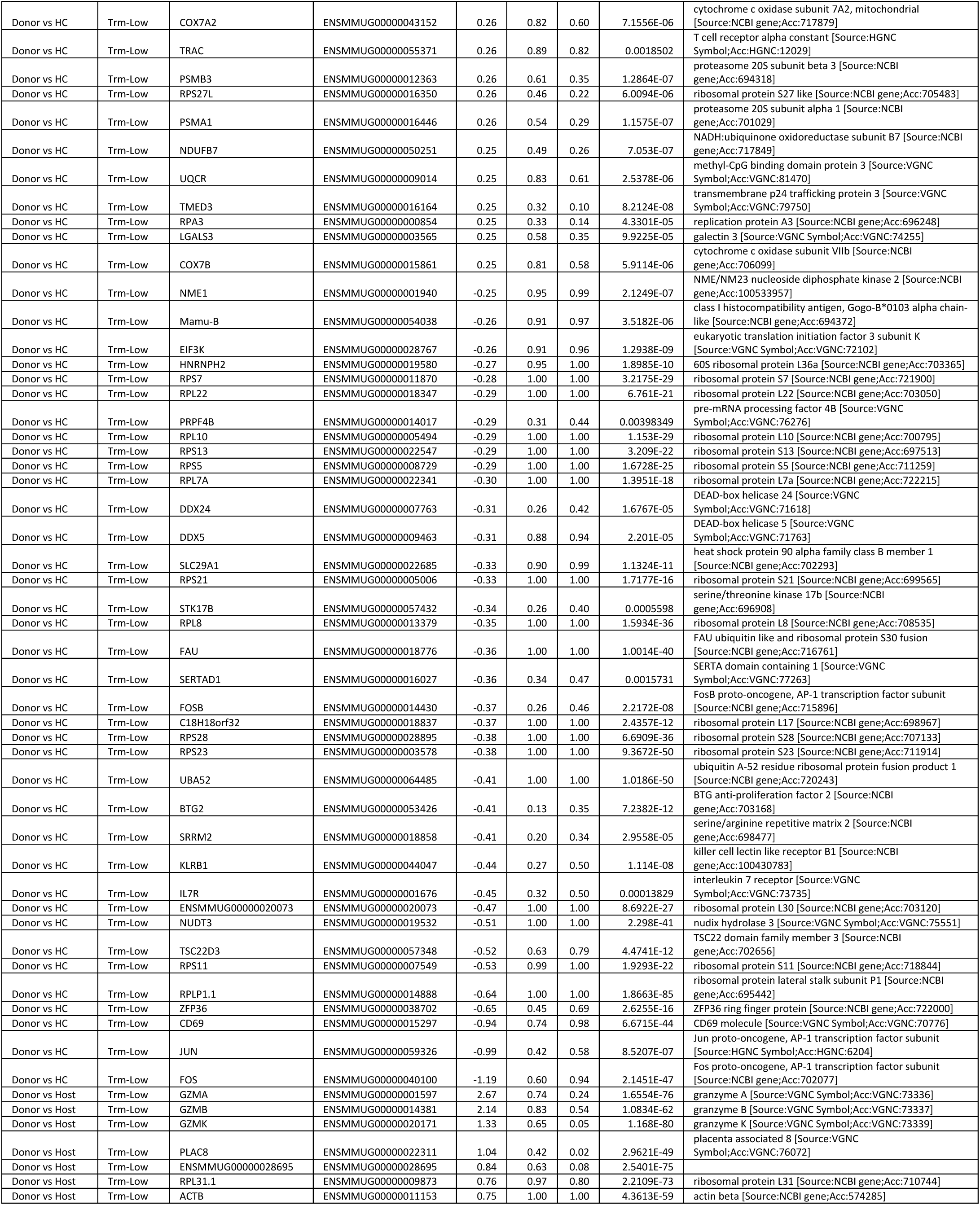
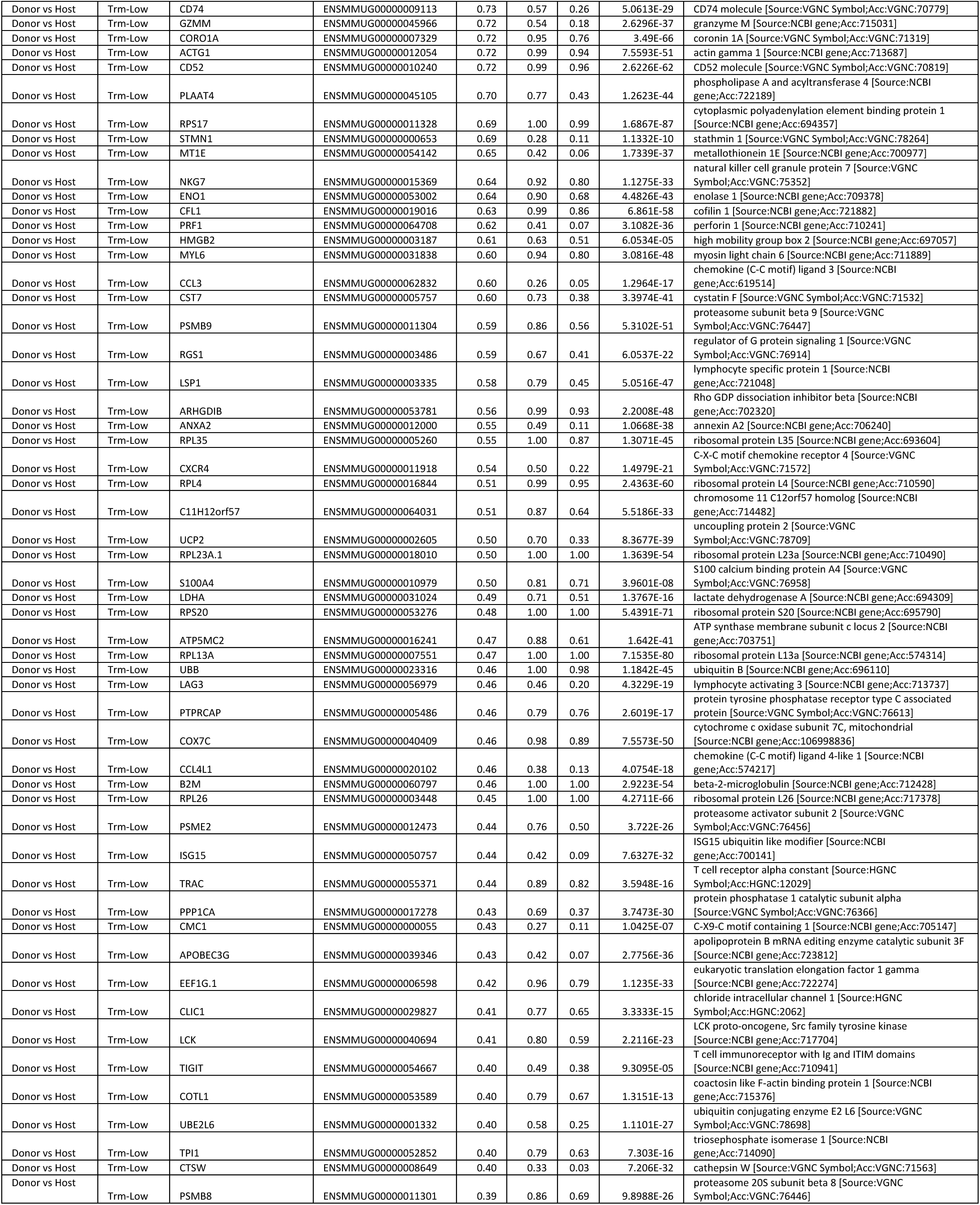
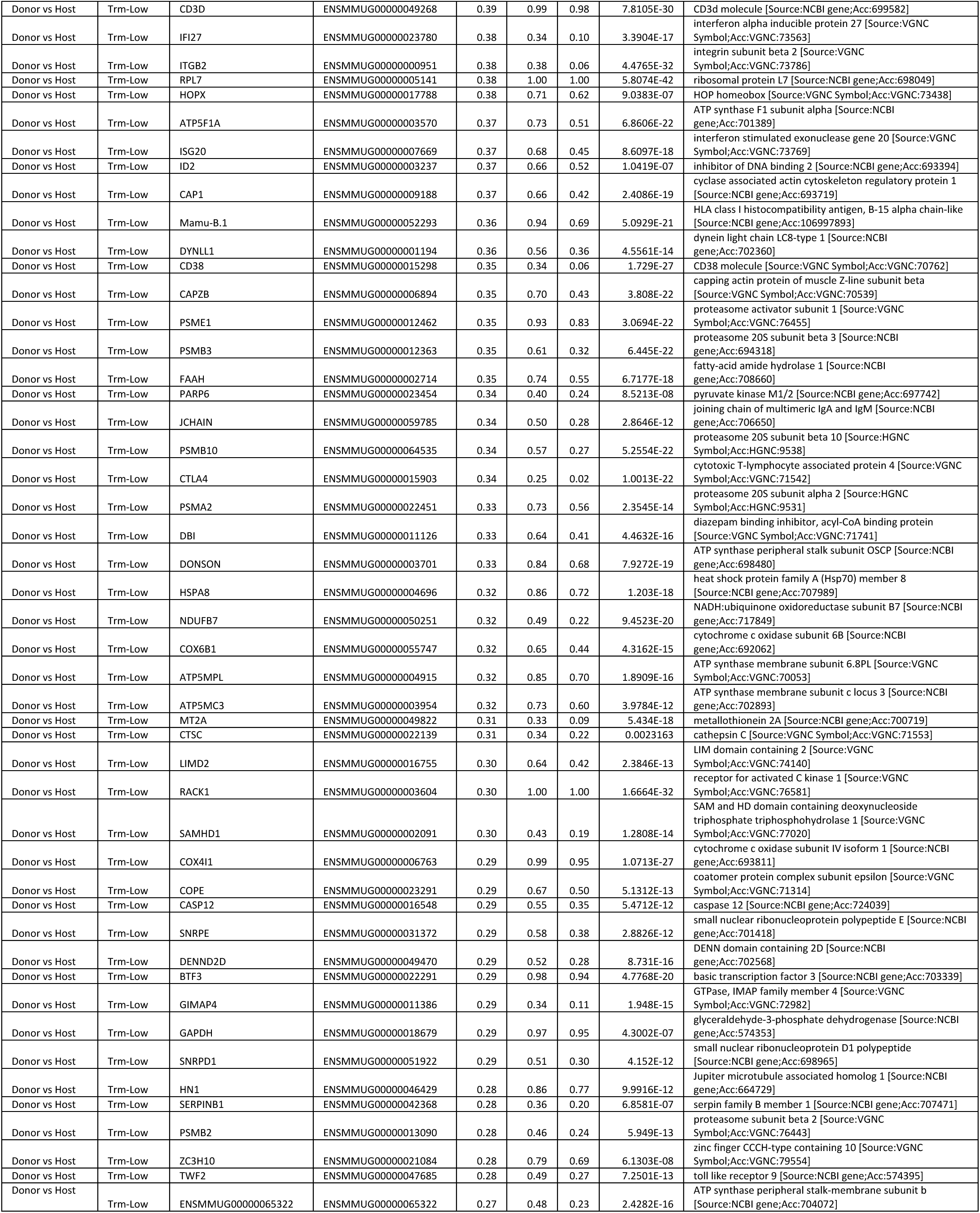
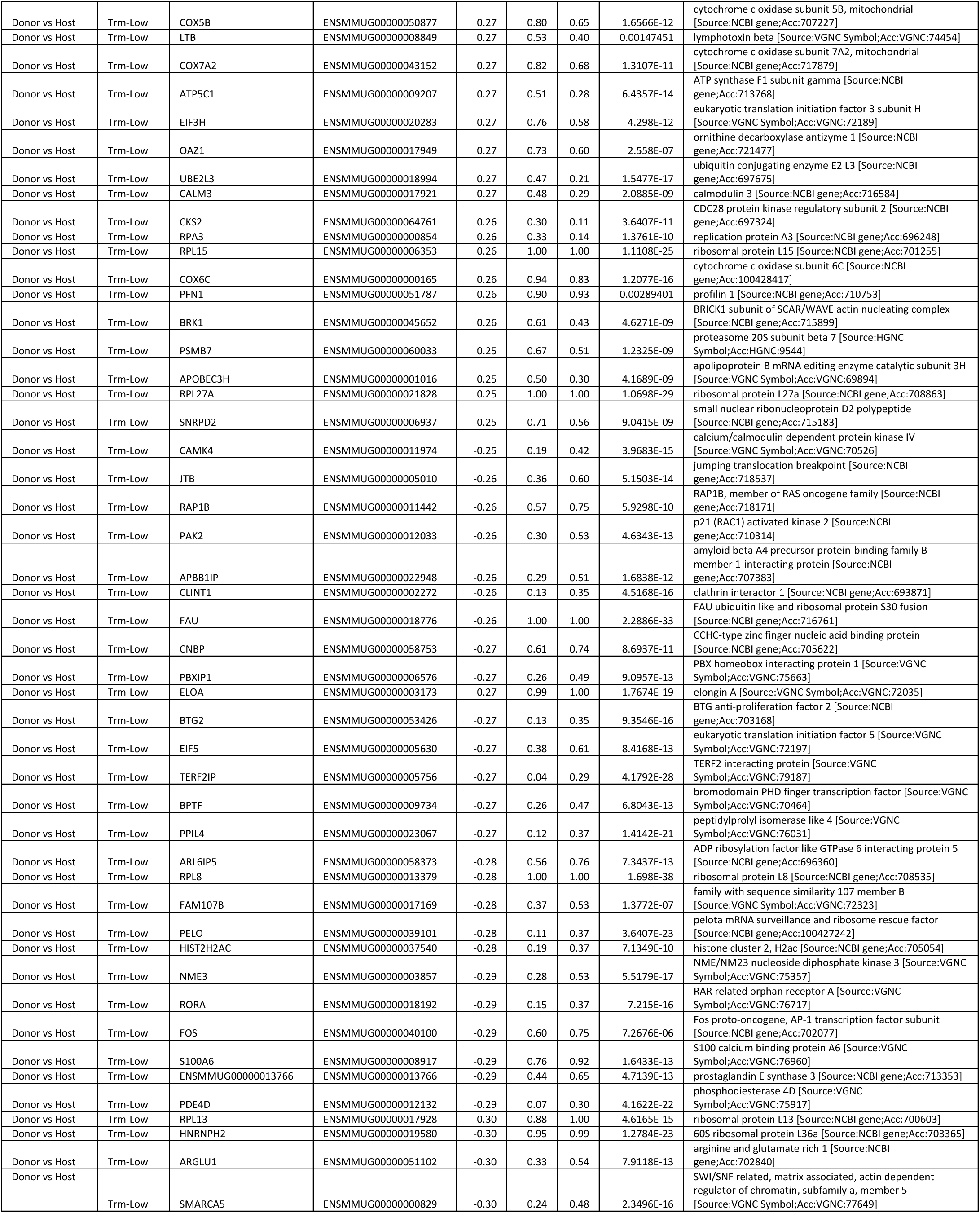
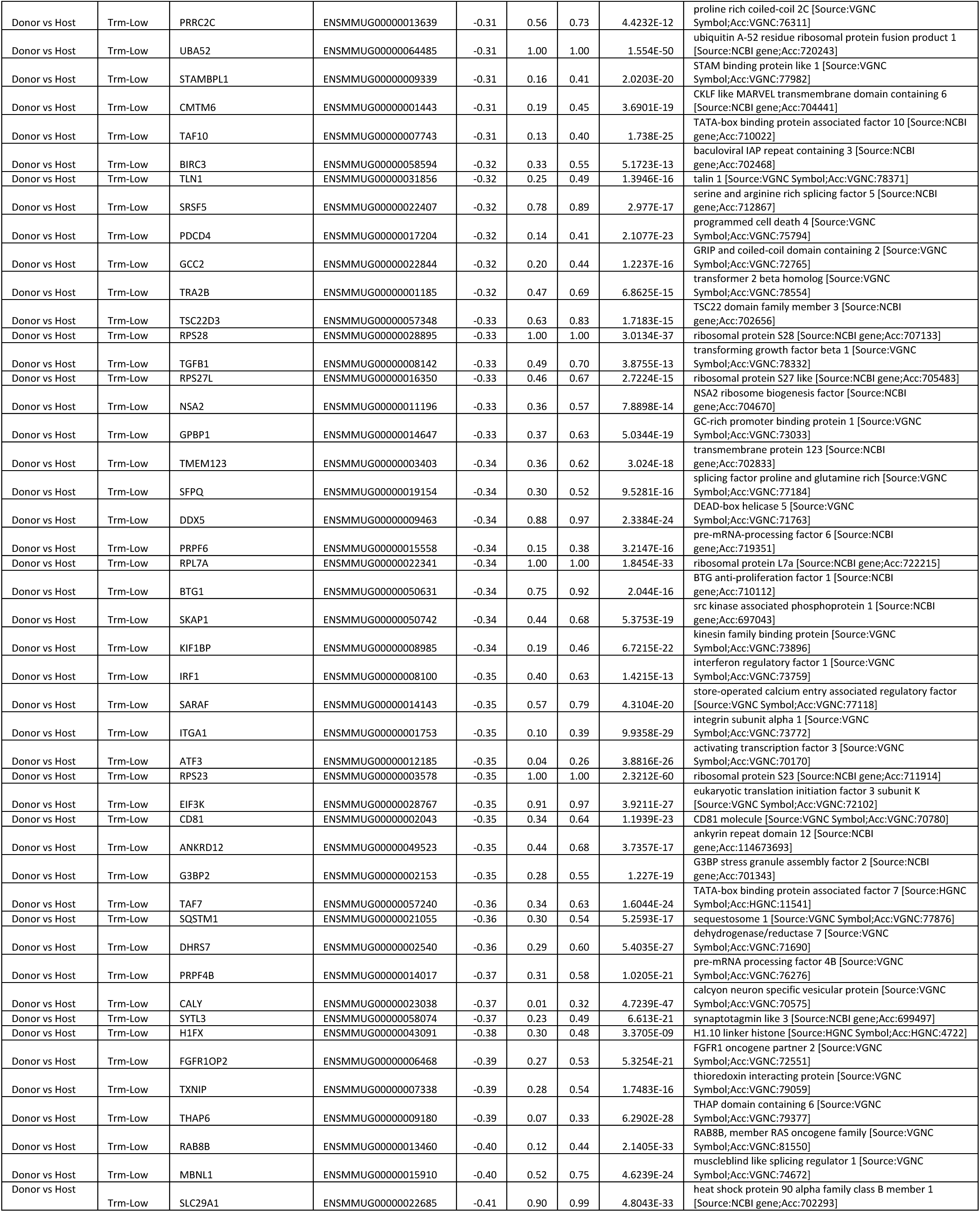
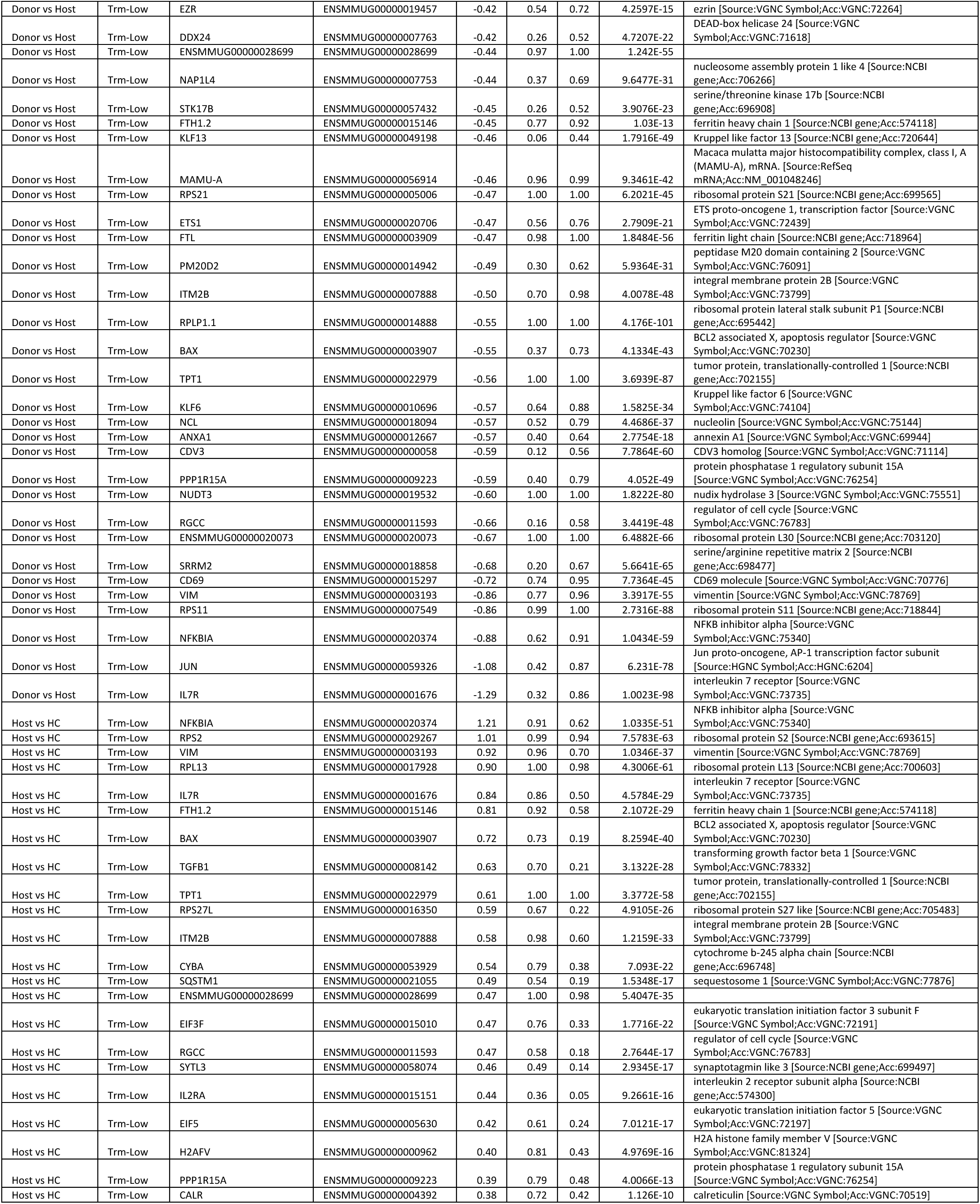
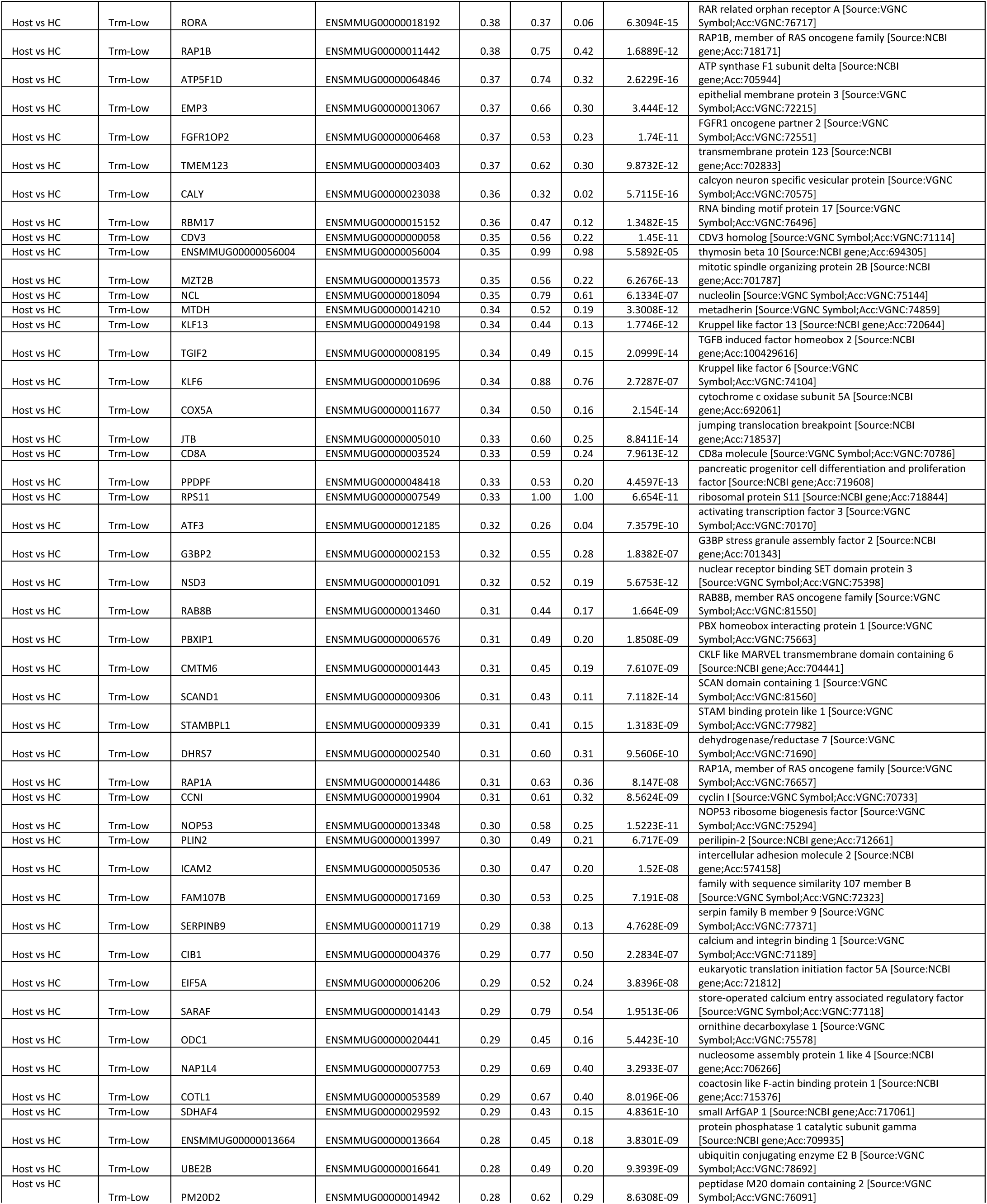
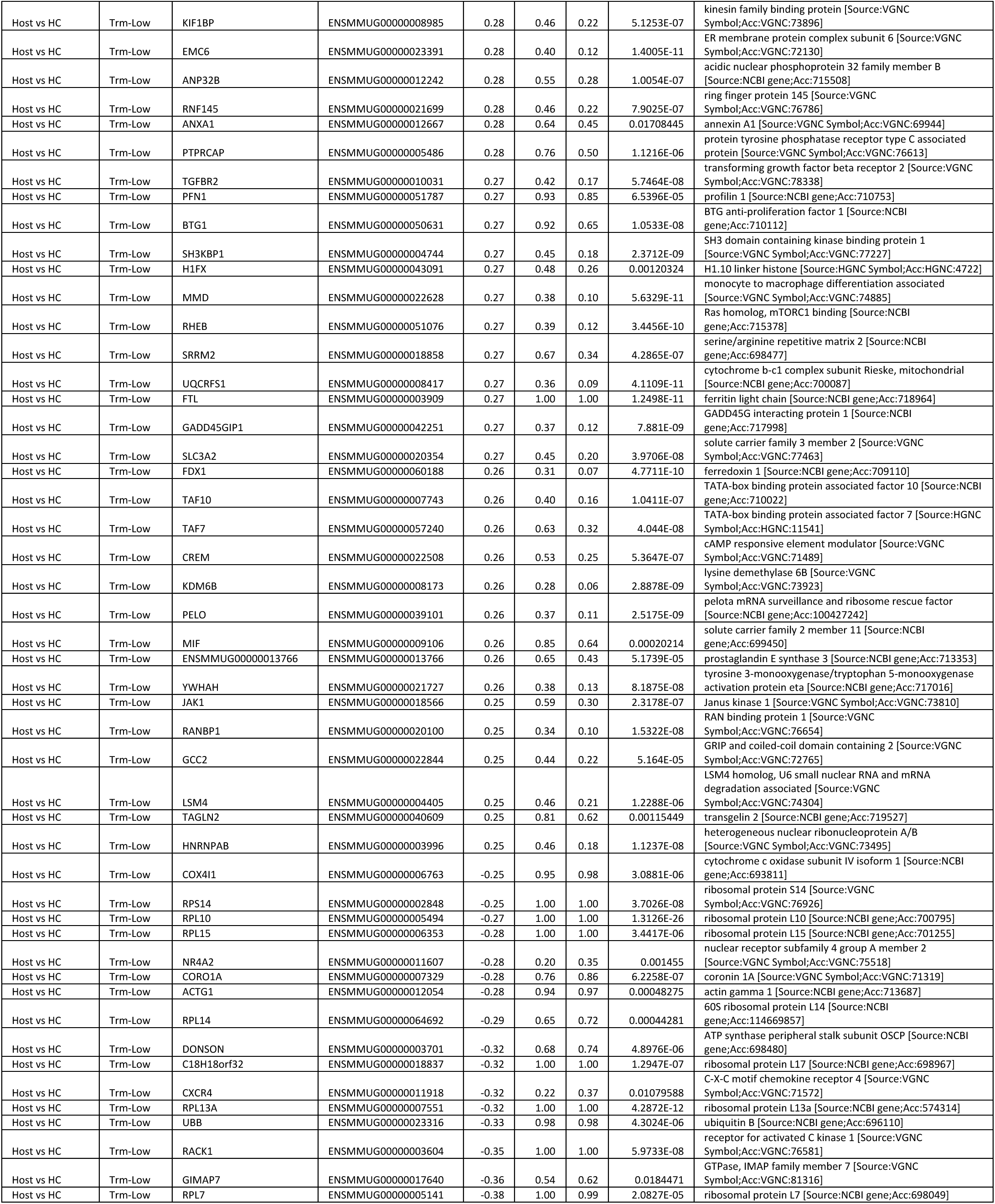
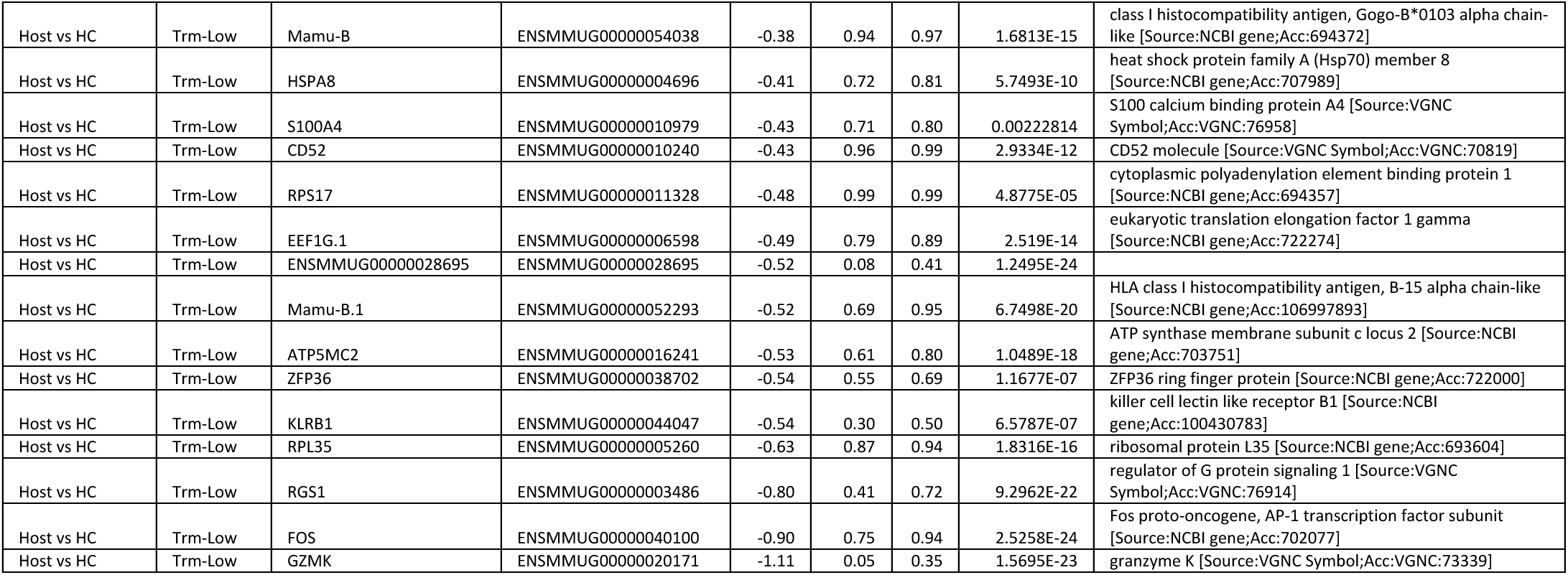
Differentially expressed (DE) genes between donor, host and healthy control CD8 T cells with low enrichment score for T_RM_ signature.

**Table S8.**
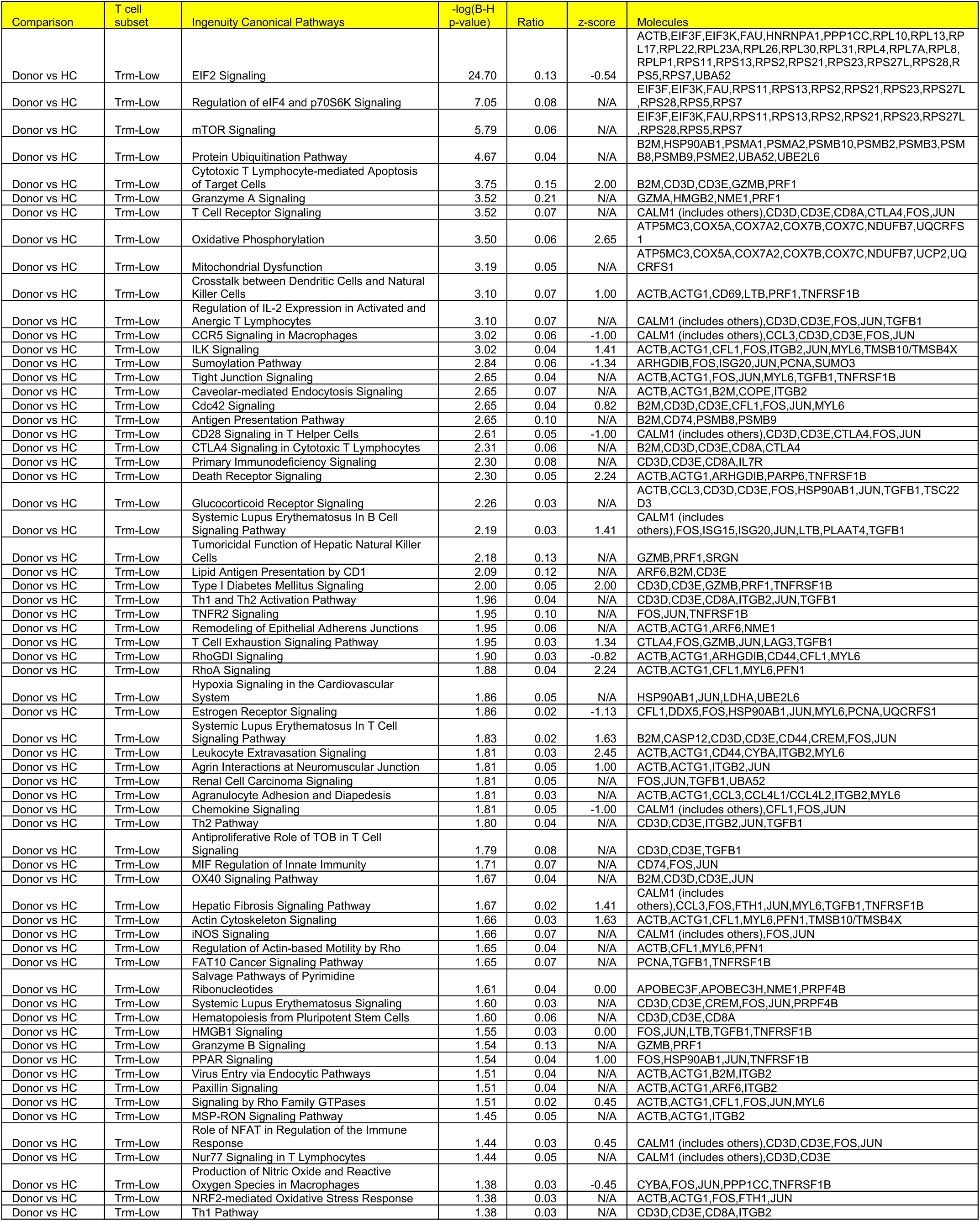
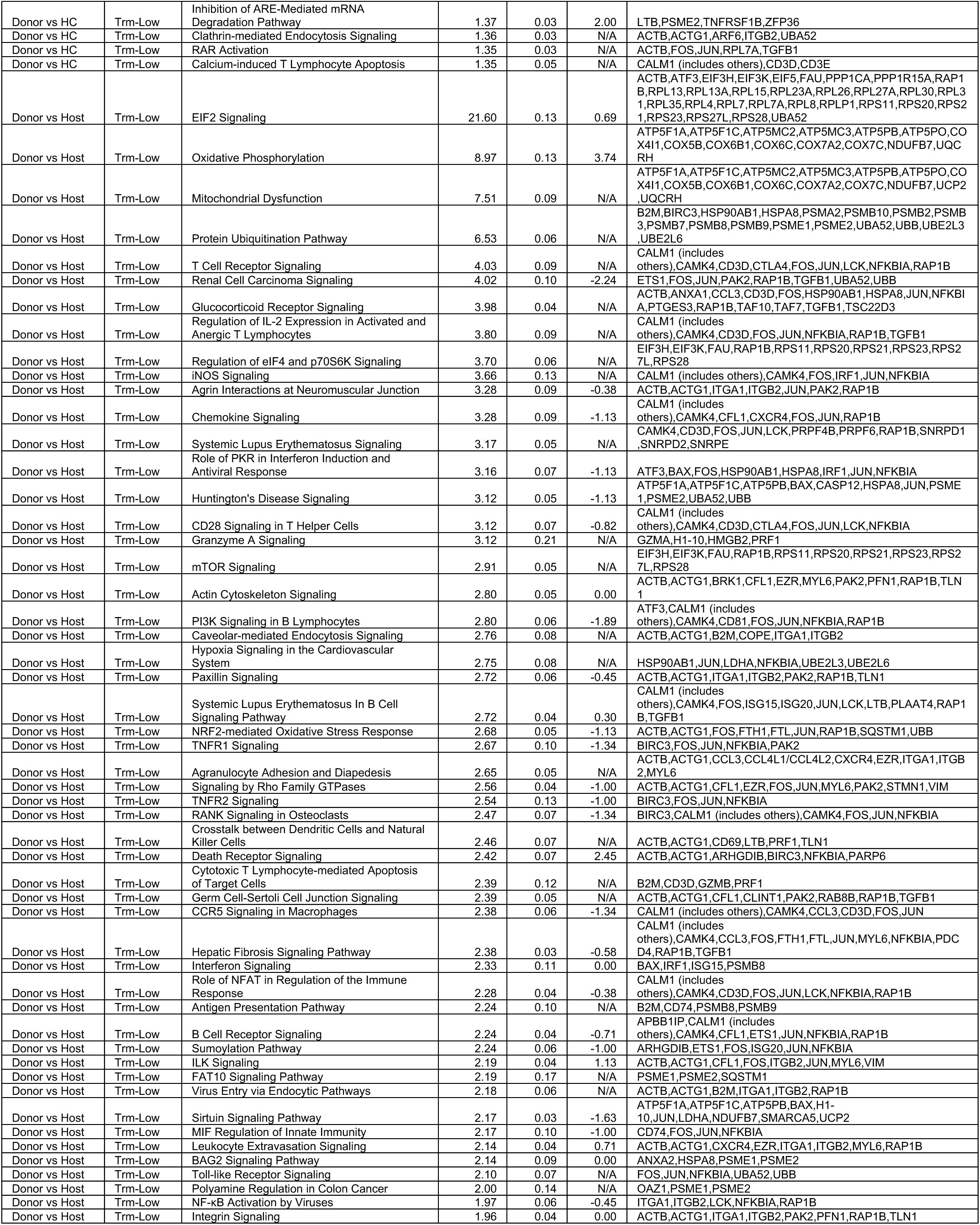
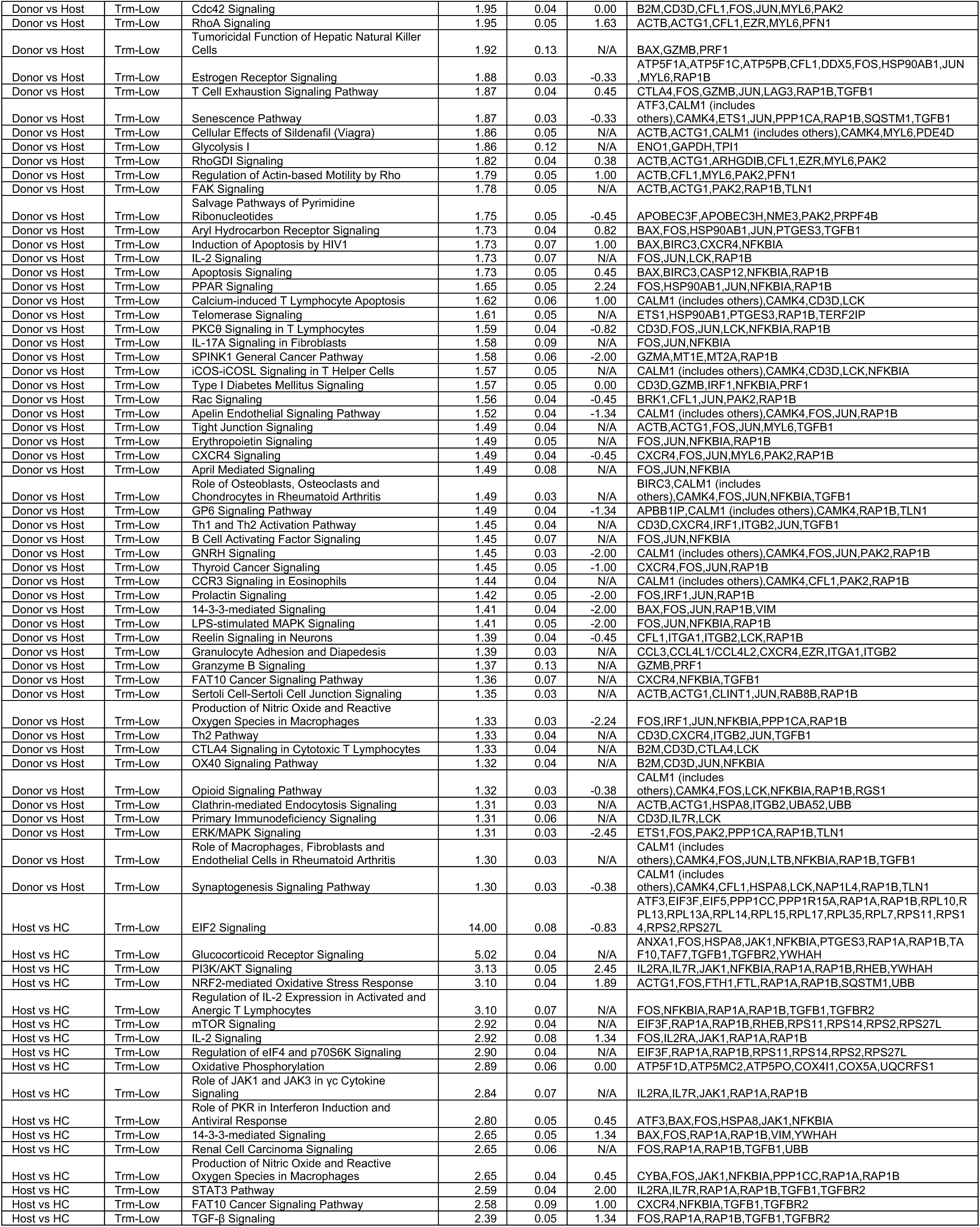
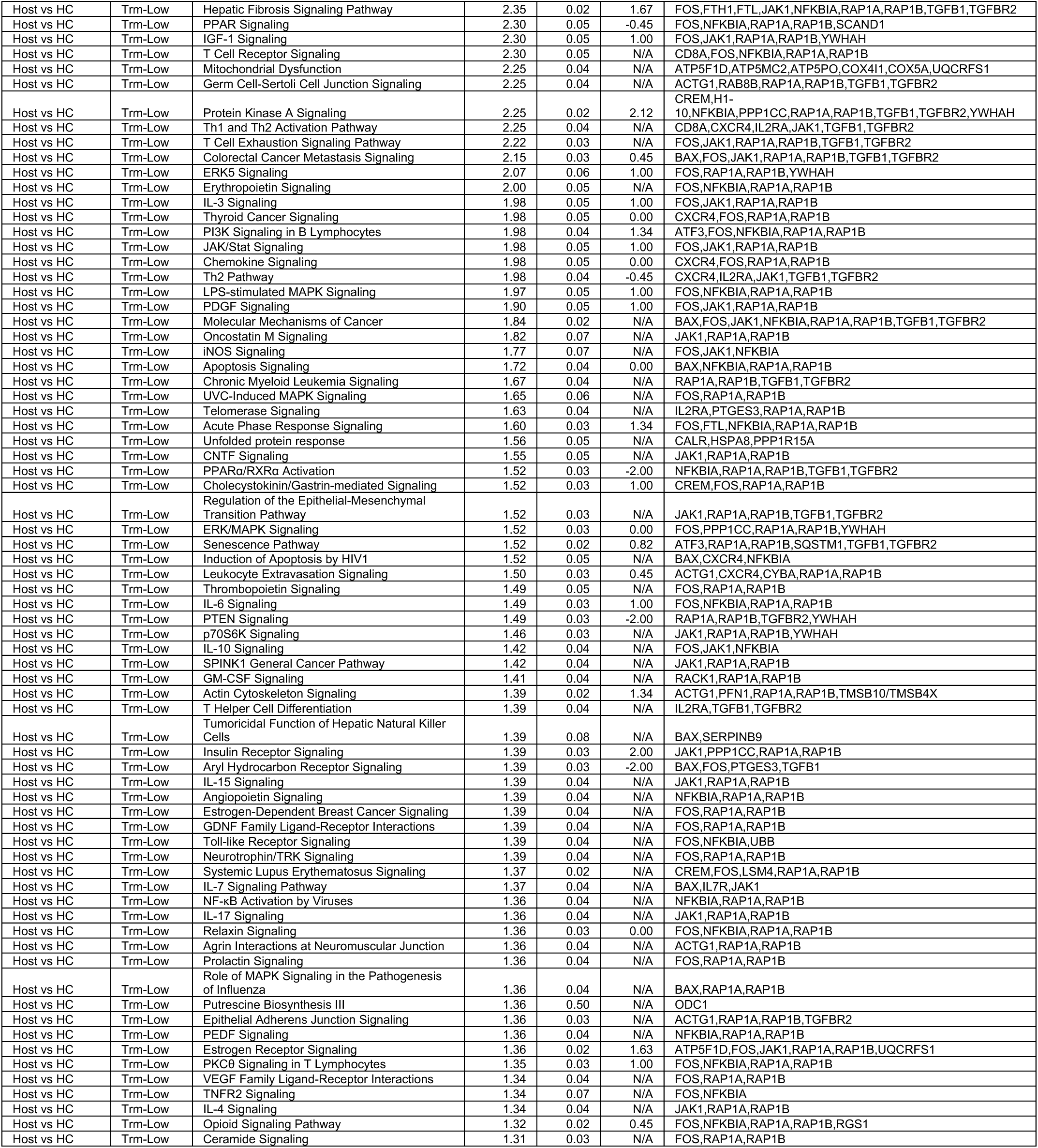
Pathway analysis on DE genes between donor, host and healthy control CD8 T cells with low enrichment score for T_RM_ signature.

**Table S9.**
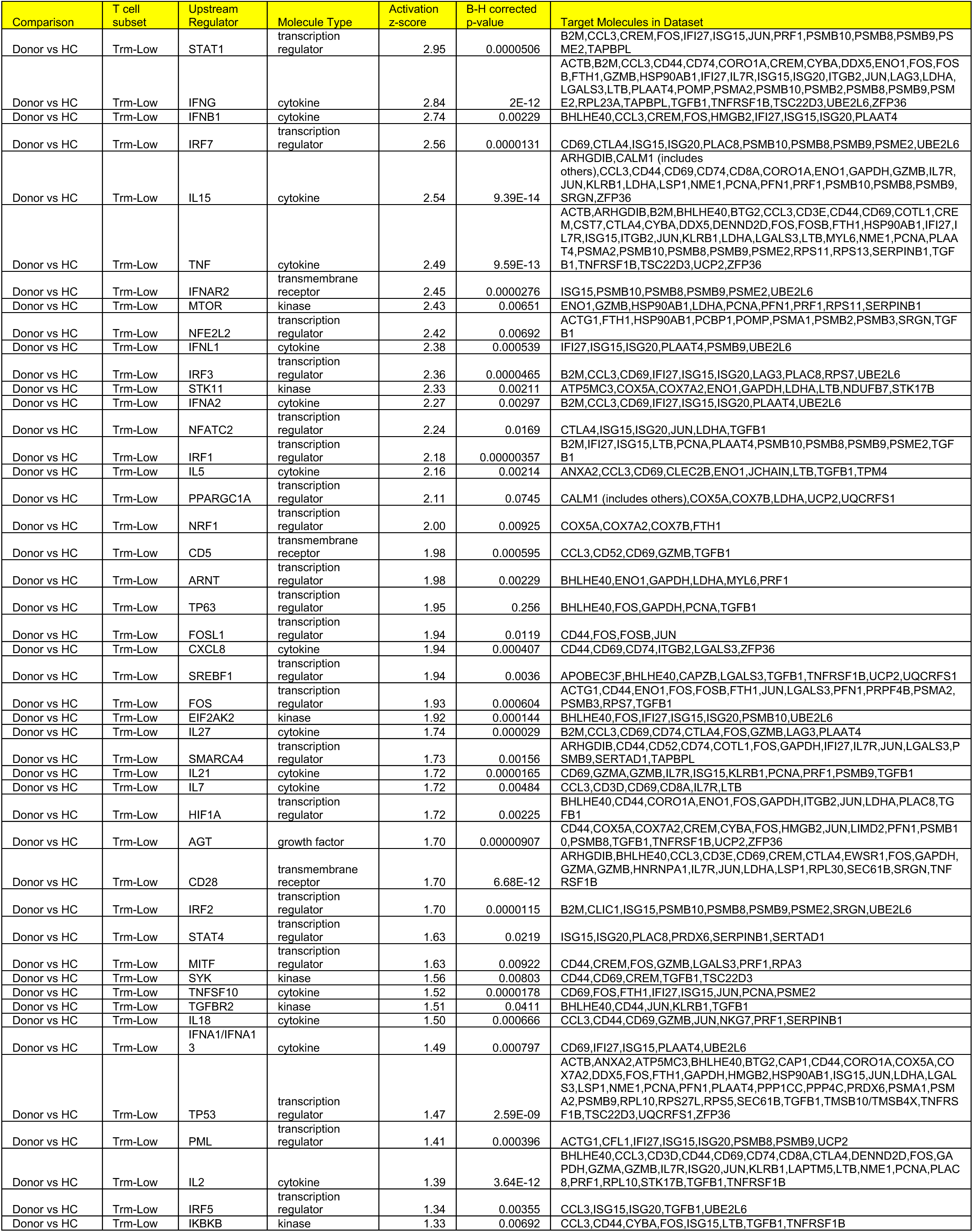
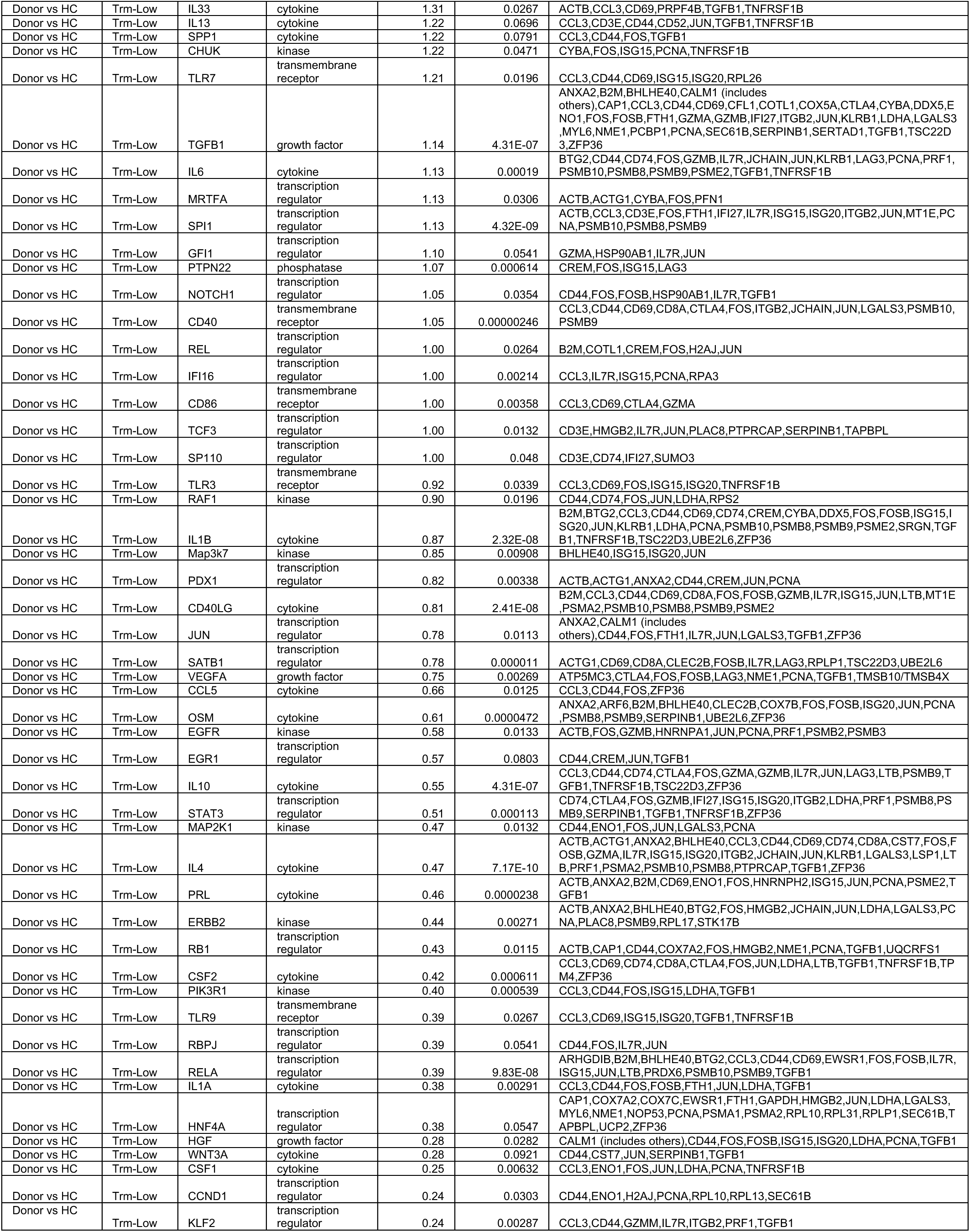
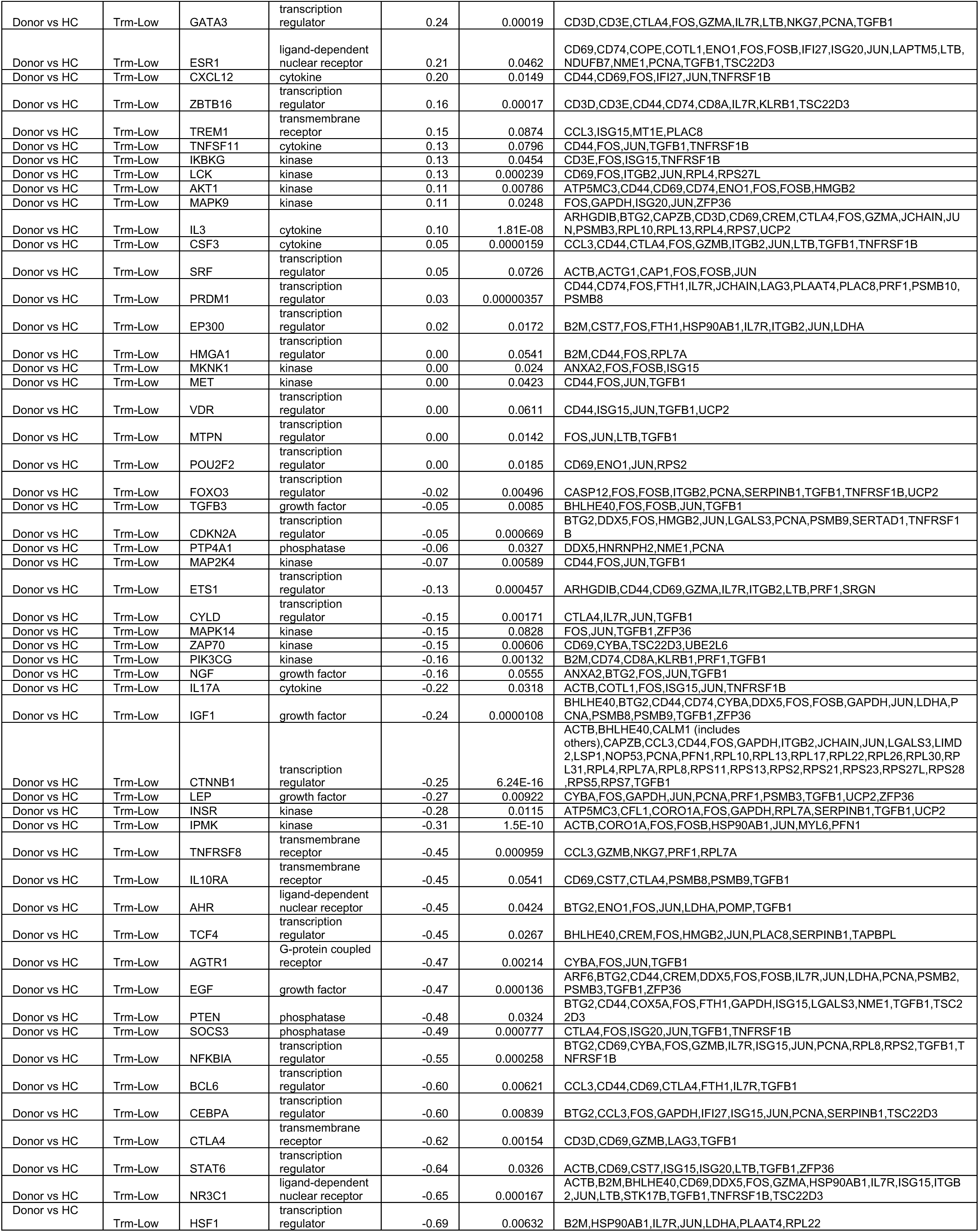
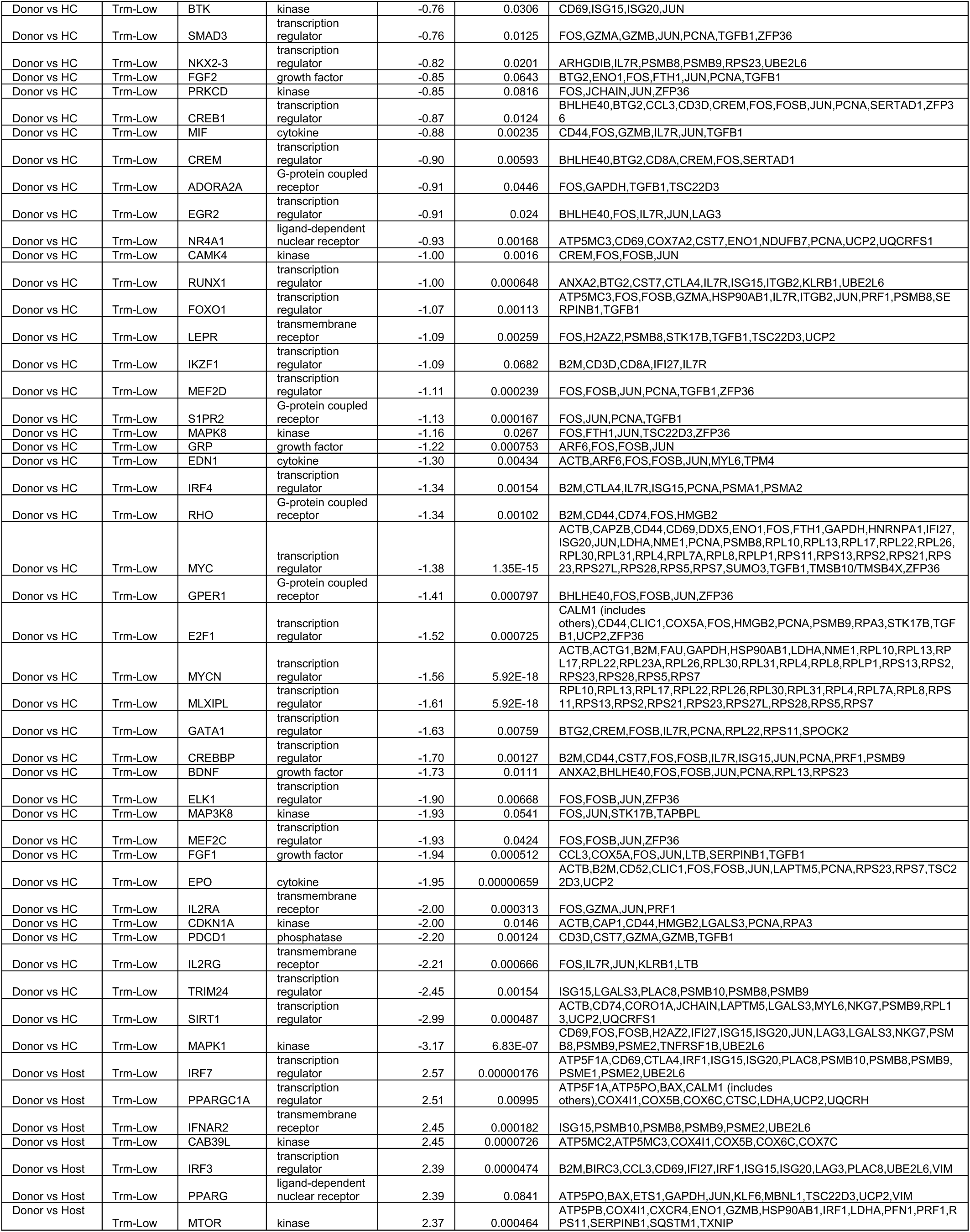
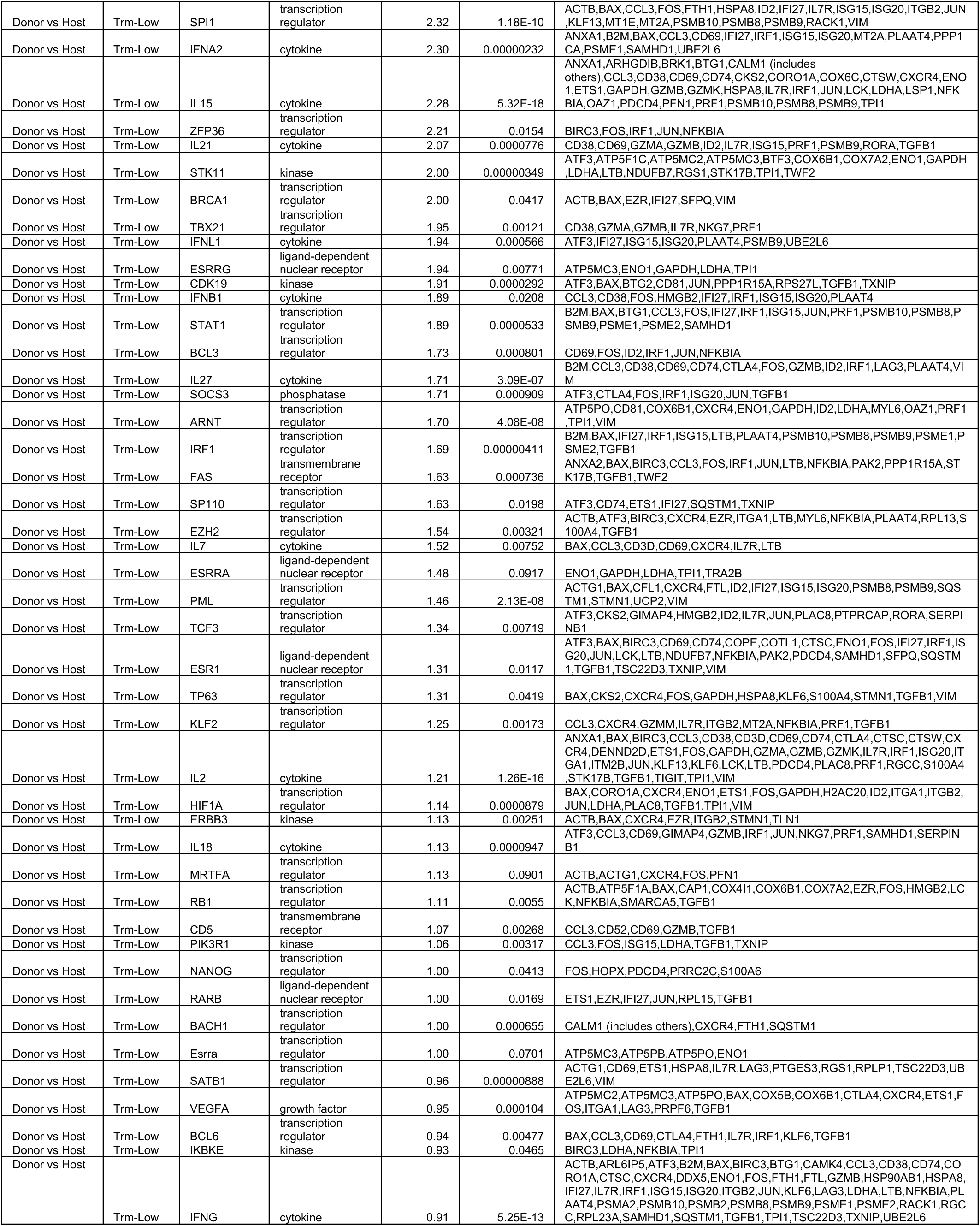
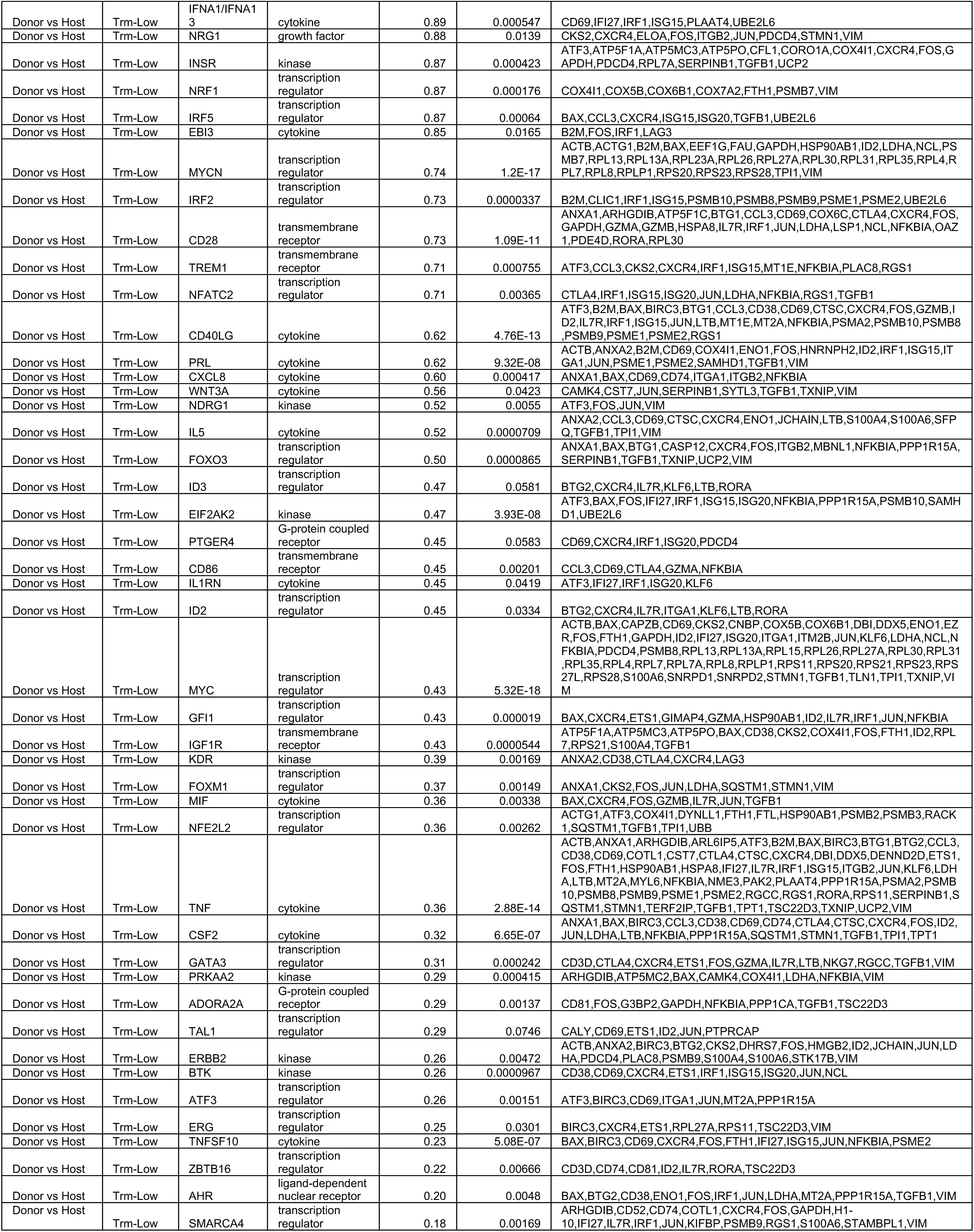
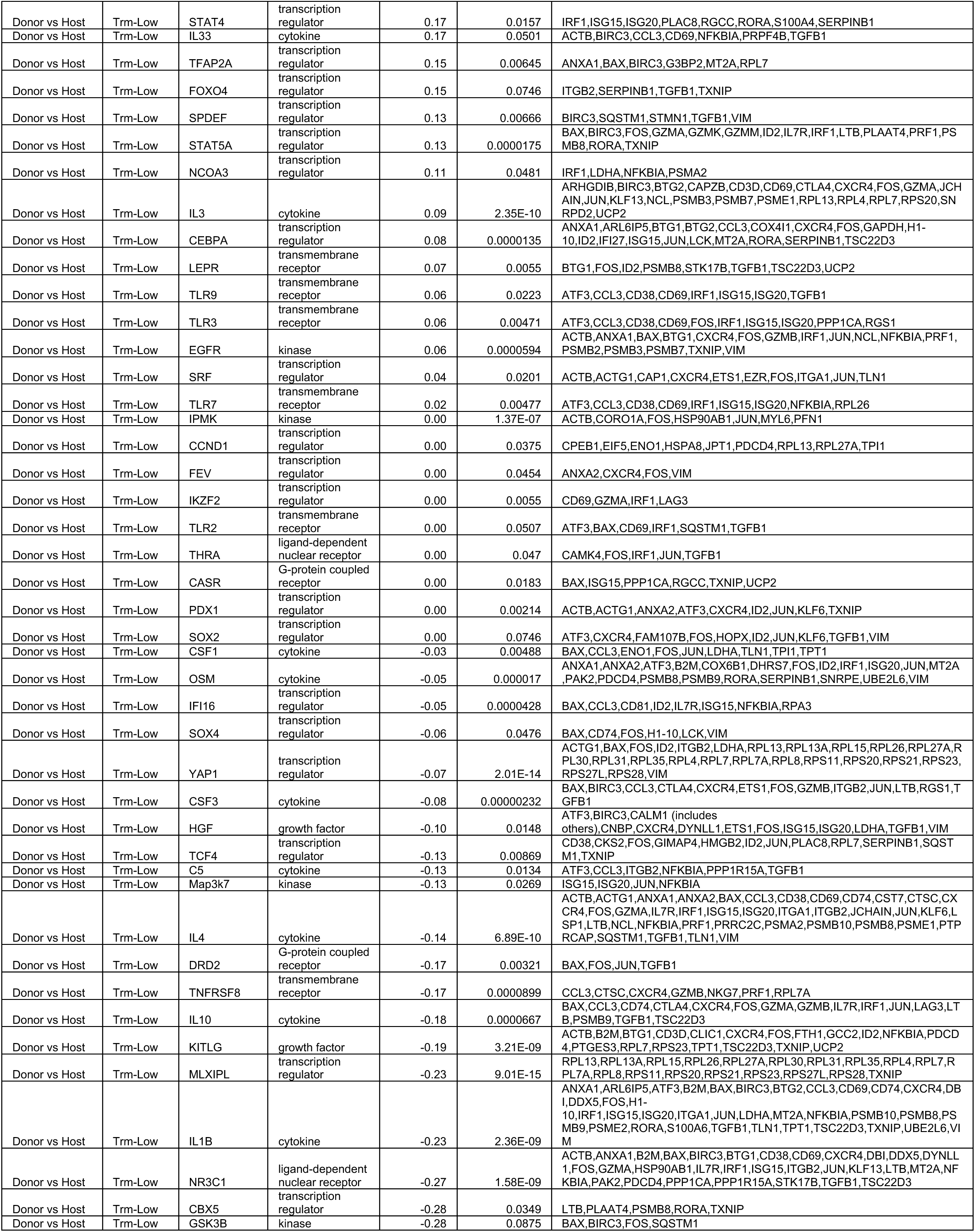
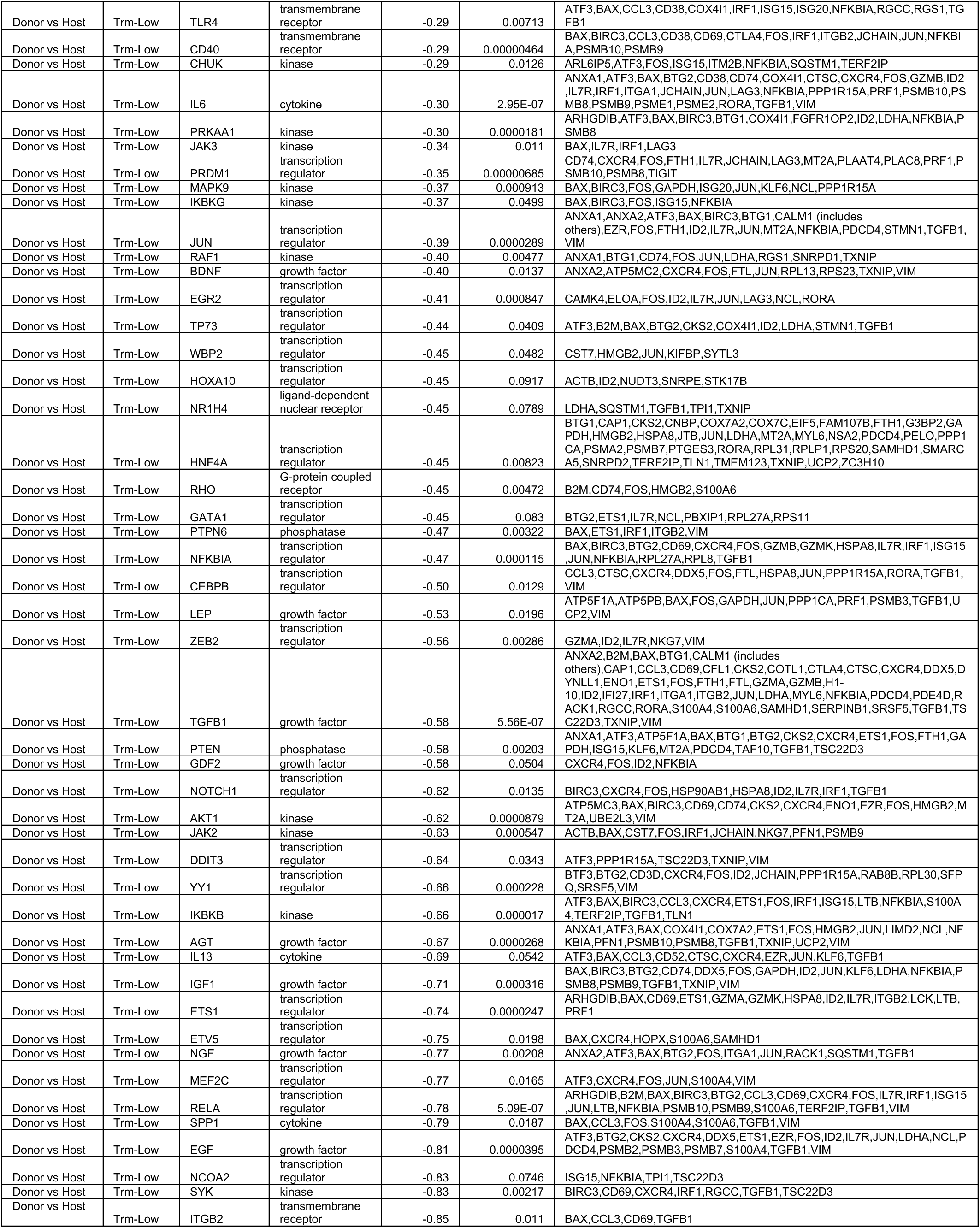
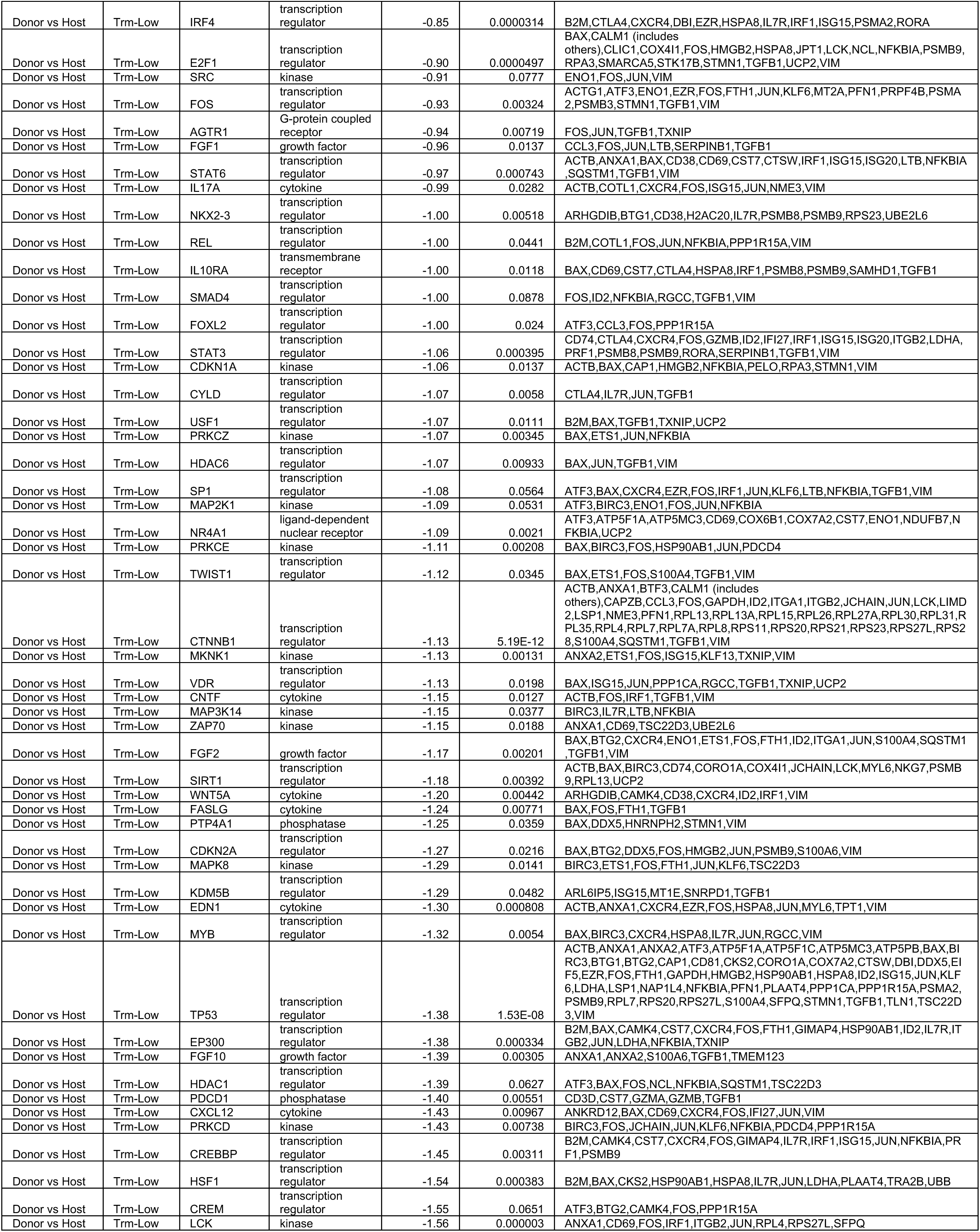
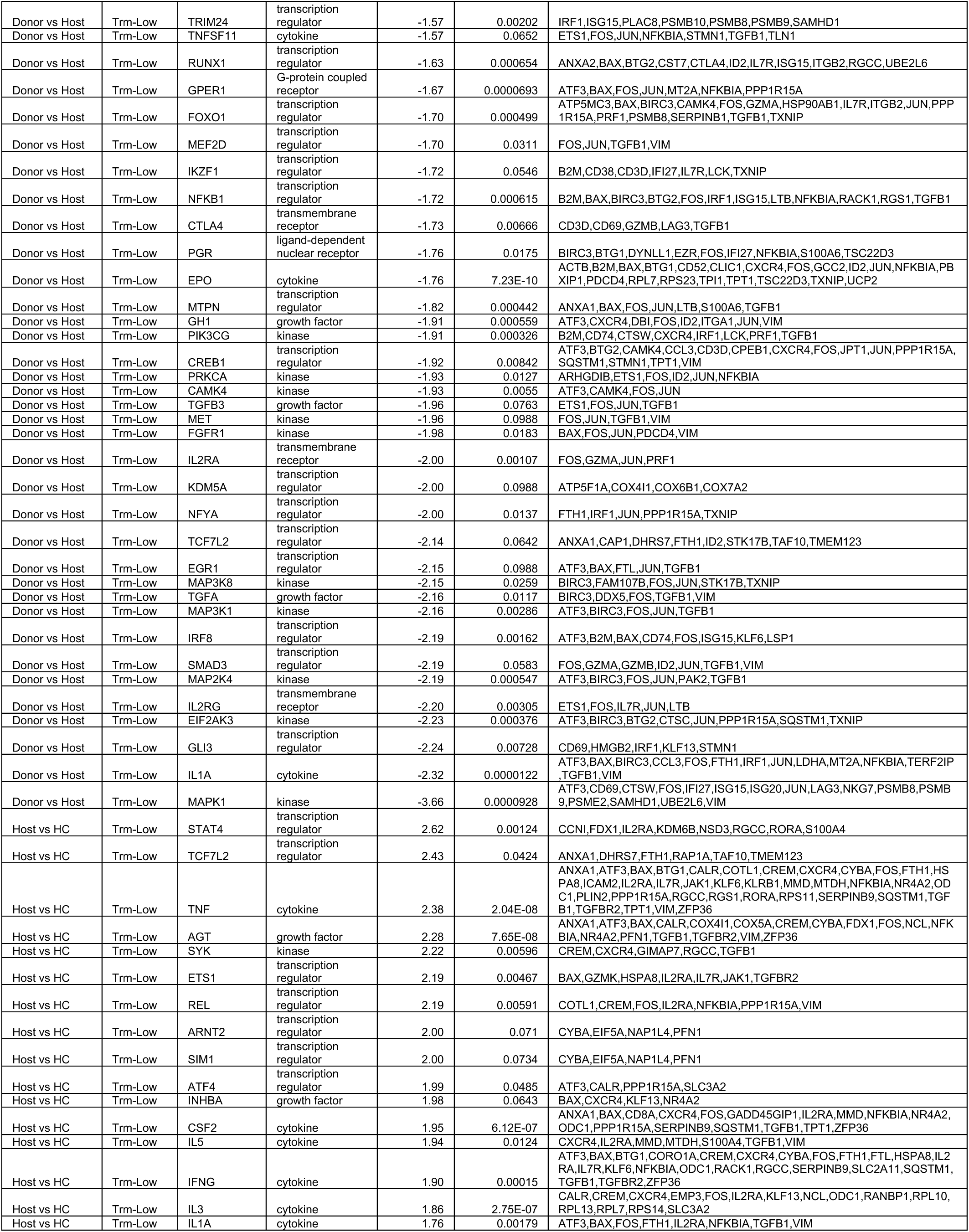
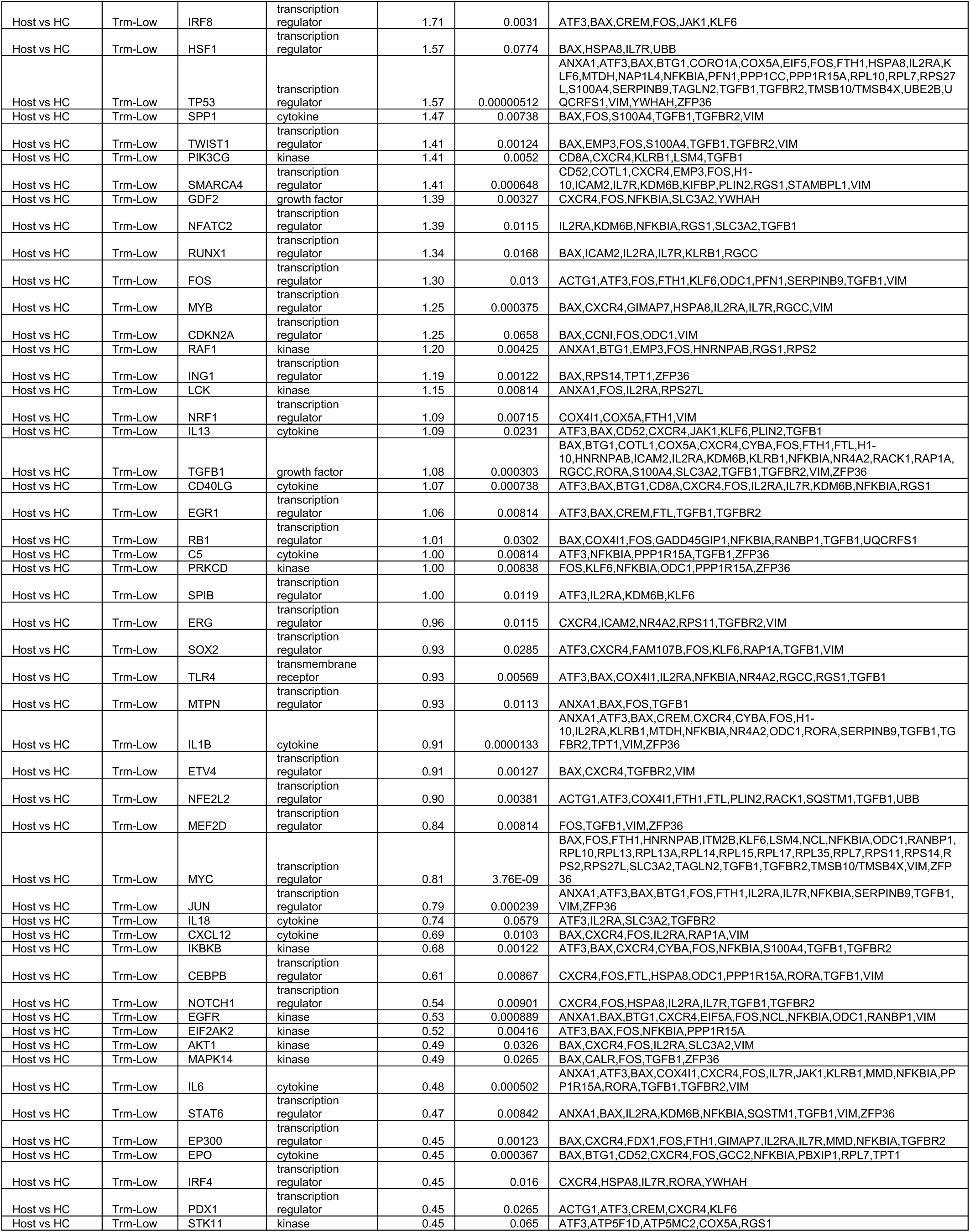
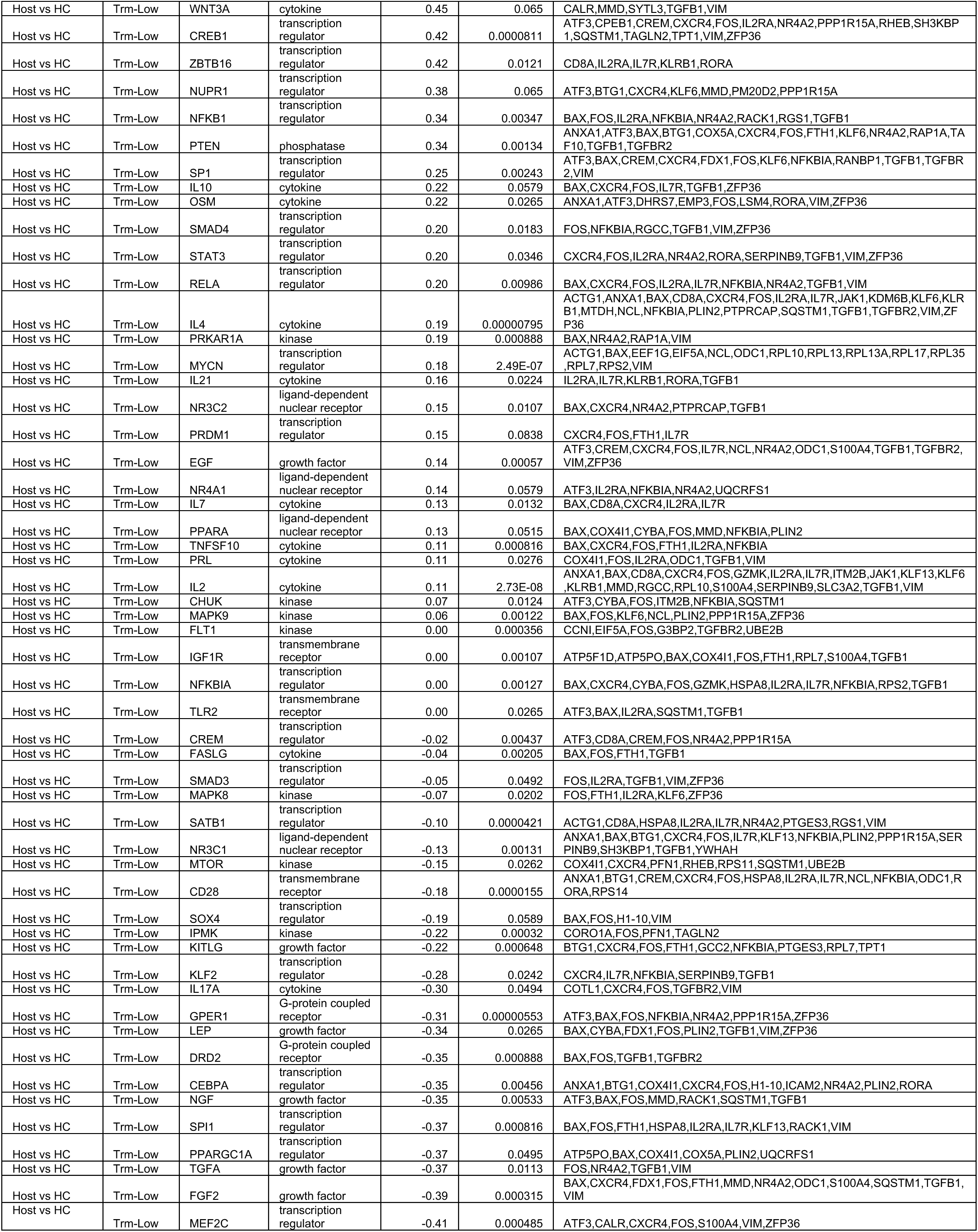
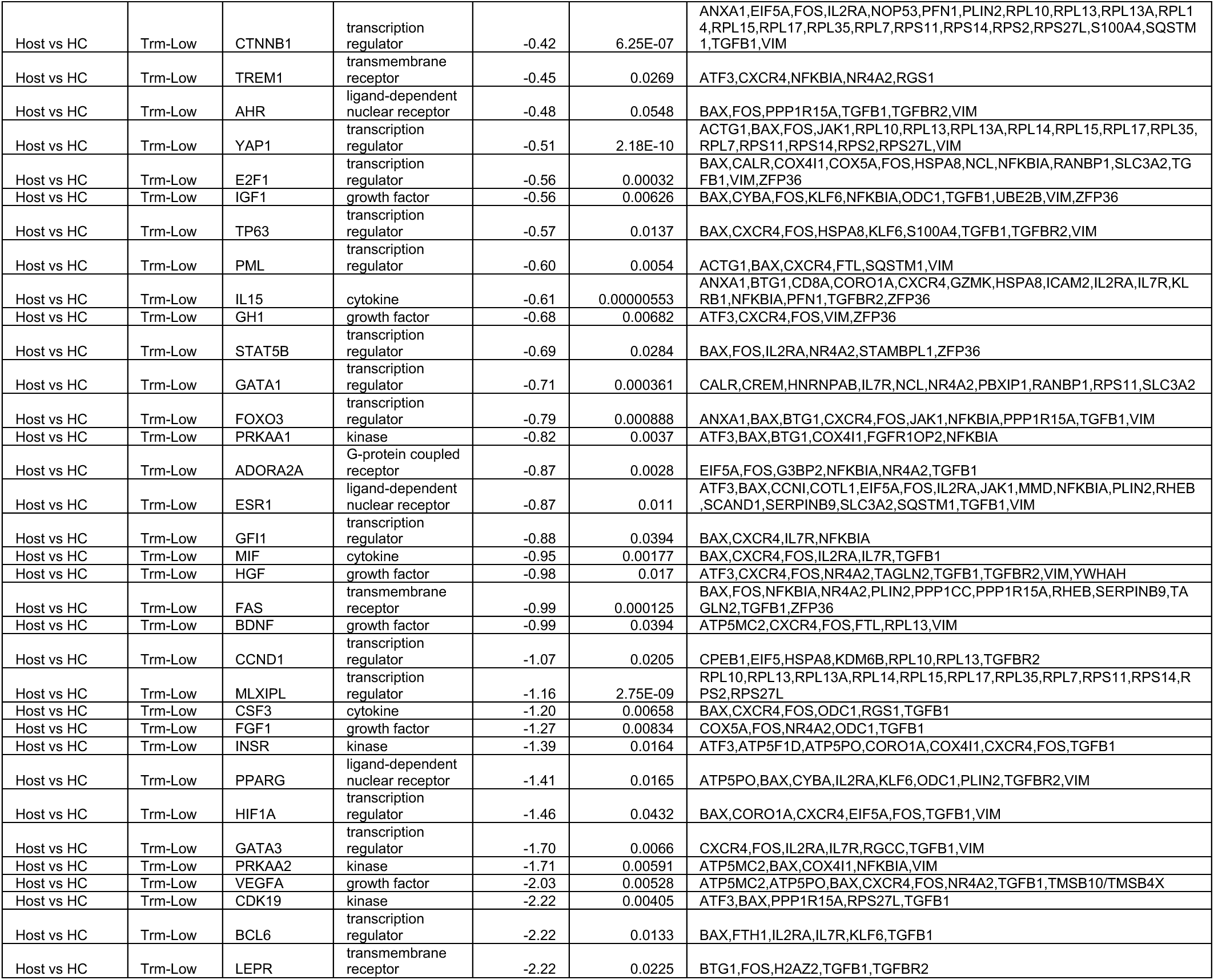
Predicted activated up-stream regulators in donor CD8 T cells with low enrichment score for T_RM_ signature in comparison with their healthy control and host counterparts.

**Table S10.** Gene signature of donor CD8 T cells with low enrichment score for T_RM_ signature, defined as shown in Figure 5B.

| Gene | Status |
| --- | --- |
| ACTB | Up-regulated |
| ACTG1 | Up-regulated |
| APOBEC3G | Up-regulated |
| APOBEC3H | Up-regulated |
| ATP5MC3 | Up-regulated |
| ATP5MPL | Up-regulated |
| BTG2 | Down-regulated |
| CALM3 | Up-regulated |
| CAP1 | Up-regulated |
| CASP12 | Up-regulated |
| CCL3 | Up-regulated |
| CCL4L1 | Up-regulated |
| CD52 | Up-regulated |
| CD74 | Up-regulated |
| COPE | Up-regulated |
| COTL1 | Up-regulated |
| COX7A2 | Up-regulated |
| COX7C | Up-regulated |
| CST7 | Up-regulated |
| DDX24 | Down-regulated |
| DDX5 | Down-regulated |
| DENND2D | Up-regulated |
| ENSMMUG00000028695 | Up-regulated |
| GAPDH | Up-regulated |
| GZMM | Up-regulated |
| HMGB2 | Up-regulated |
| HNRNPH2 | Down-regulated |
| HOPX | Up-regulated |
| IFI27 | Up-regulated |
| IL7R | Down-regulated |
| ISG15 | Up-regulated |
| ISG20 | Up-regulated |
| ITGB2 | Up-regulated |
| JCHAIN | Up-regulated |
| LDHA | Up-regulated |
| LIMD2 | Up-regulated |
| LSP1 | Up-regulated |
| LTB | Up-regulated |
| NDUFB7 | Up-regulated |
| NKG7 | Up-regulated |
| PARP6 | Up-regulated |
| PFN1 | Up-regulated |
| PLAAT4 | Up-regulated |
| PLAC8 | Up-regulated |
| PRPF4B | Down-regulated |
| PSMB10 | Up-regulated |
| PSMB2 | Up-regulated |
| PSMB8 | Up-regulated |
| PSME2 | Up-regulated |
| PTPRCAP | Up-regulated |
| RPL26 | Up-regulated |
| TSC22D3 | Down-regulated |
| UBE2L6 | Up-regulated |

**Table S11.**
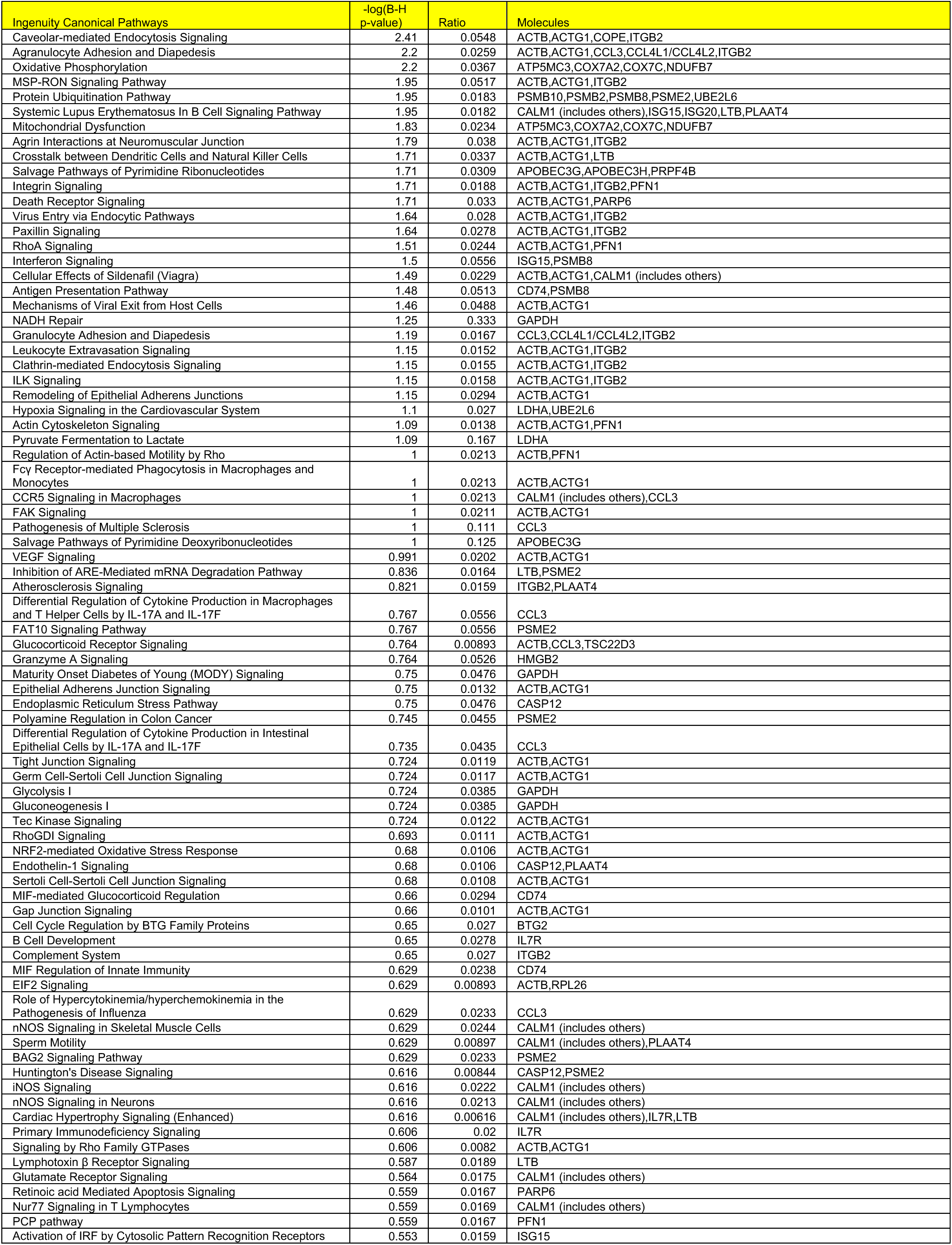
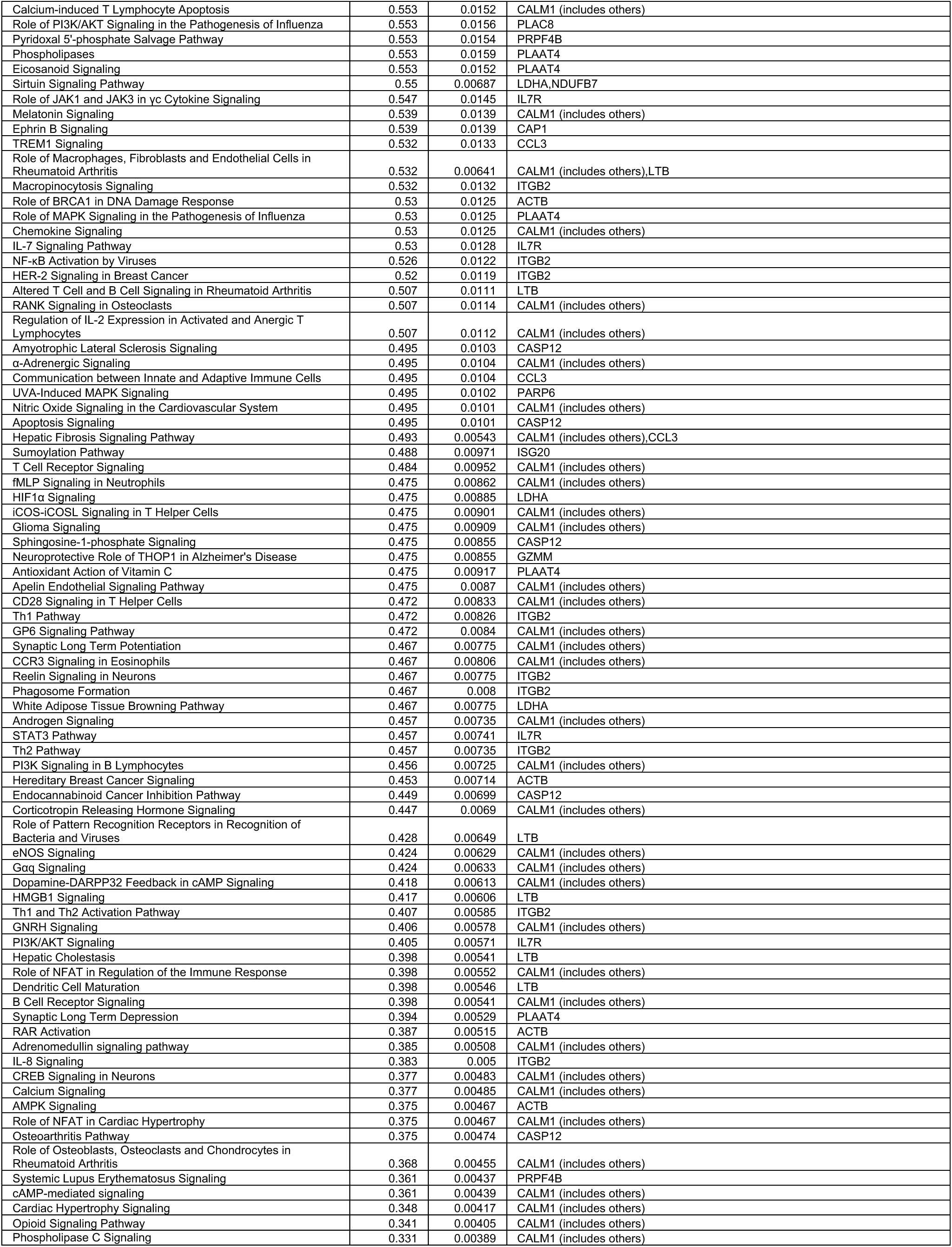
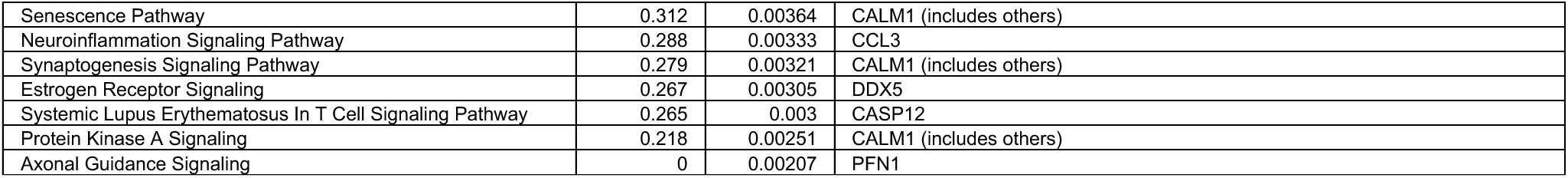
Pathway analysis on the gene signature of donor CD8 T cells with low enrichment score for T_RM_ signature.

**Table S12.**
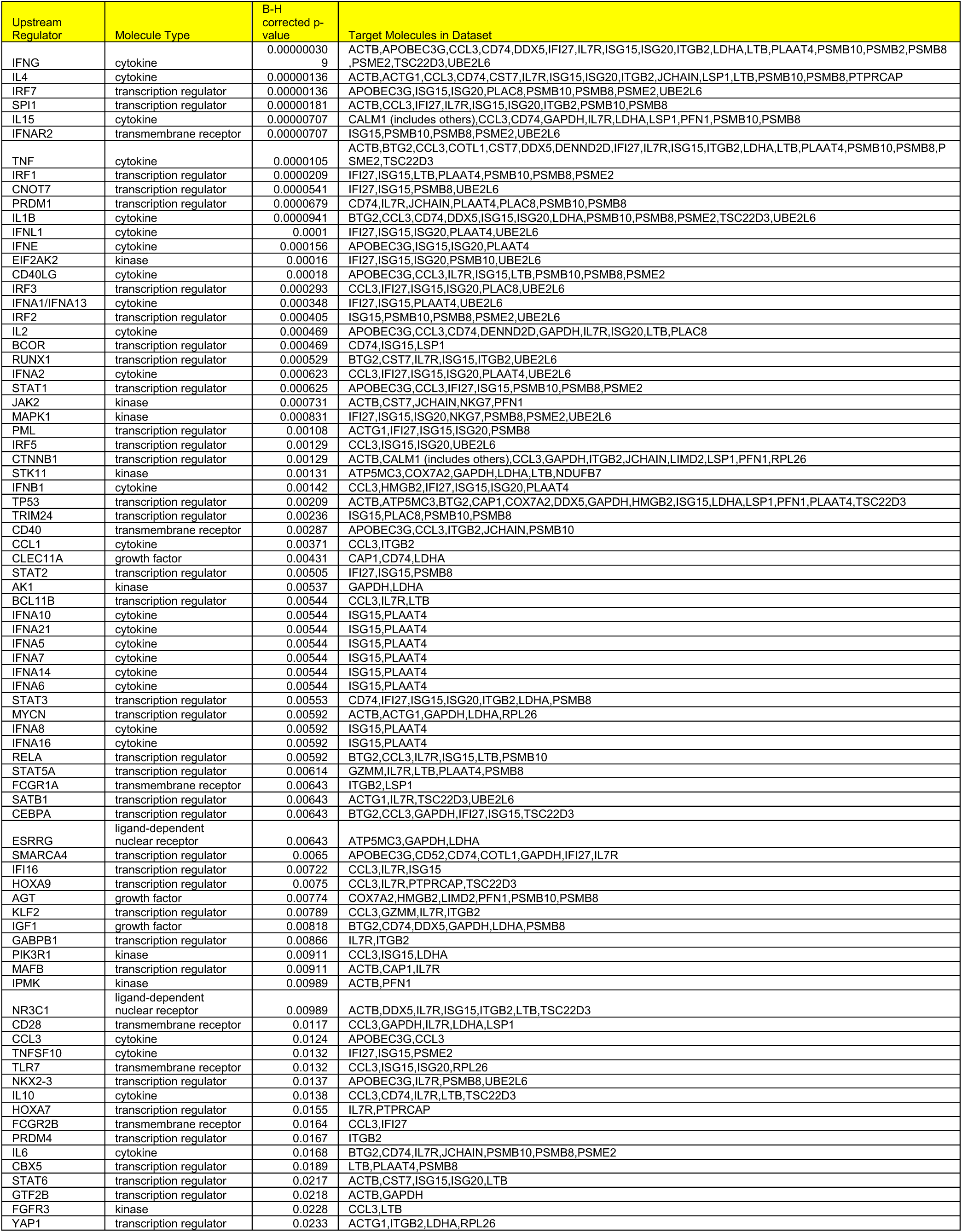
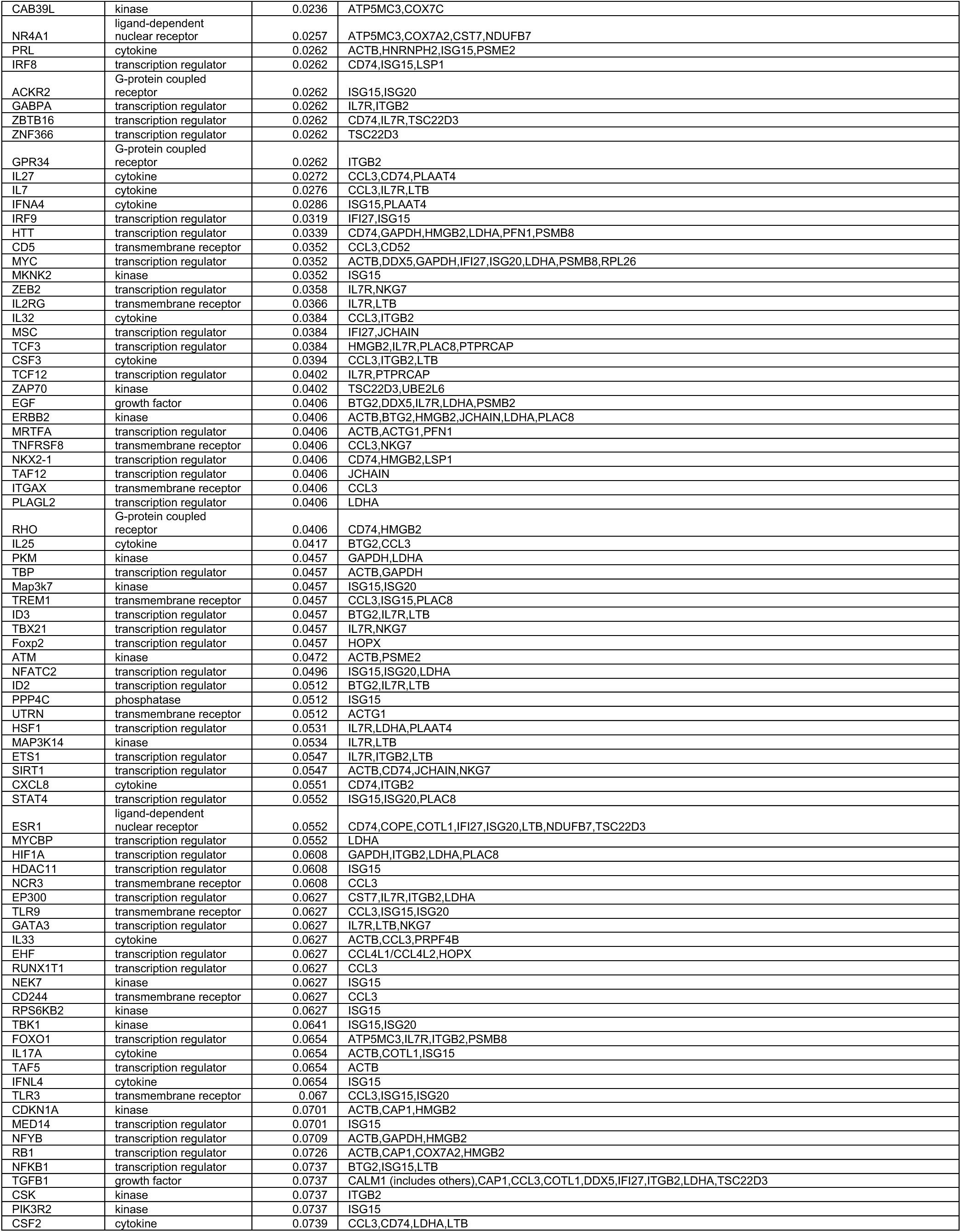
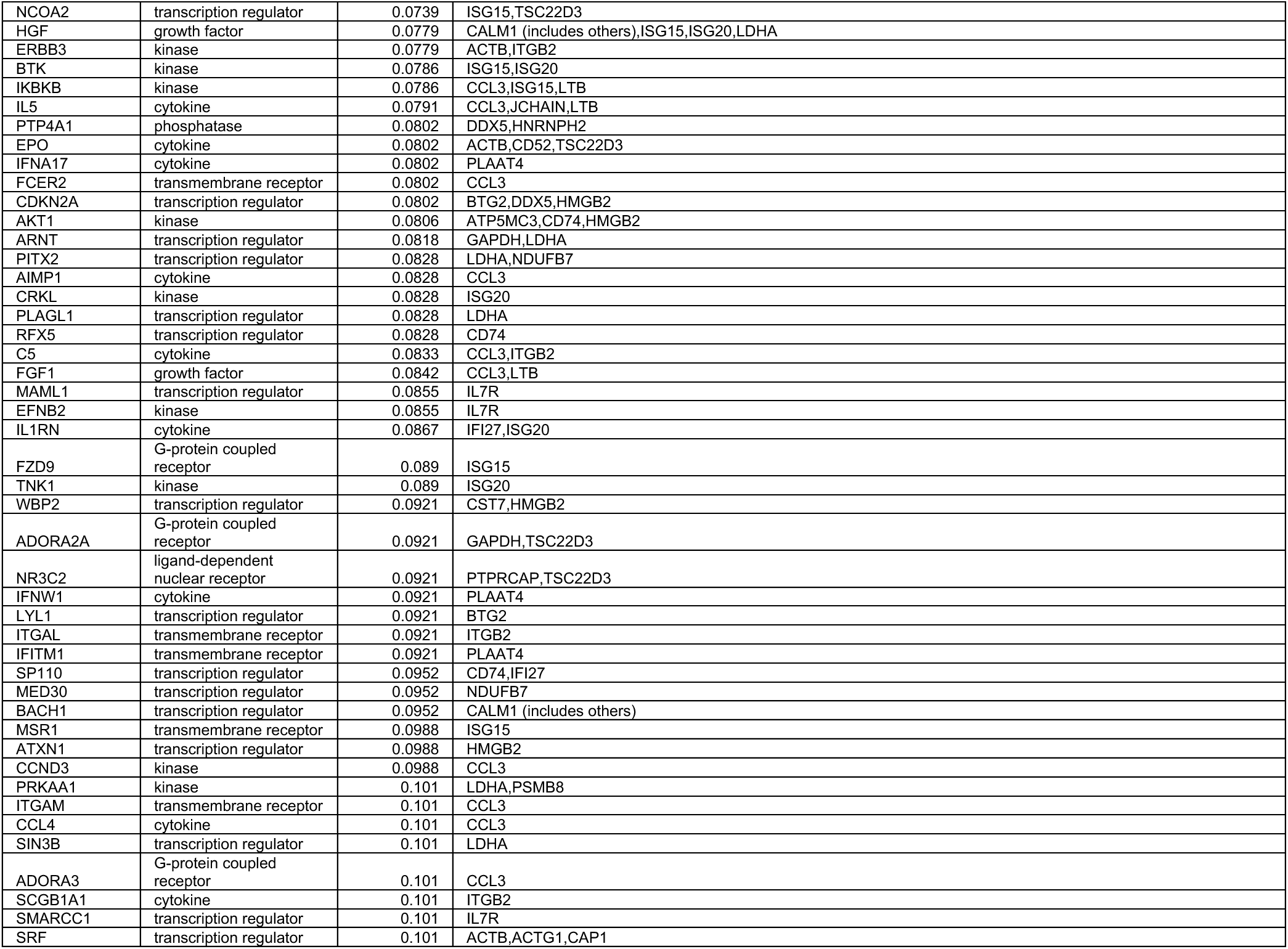
Predicted activated up-stream regulators based on the gene signature of donor CD8 T cells with low enrichment score for T_RM_ signature.

**Table S13.**
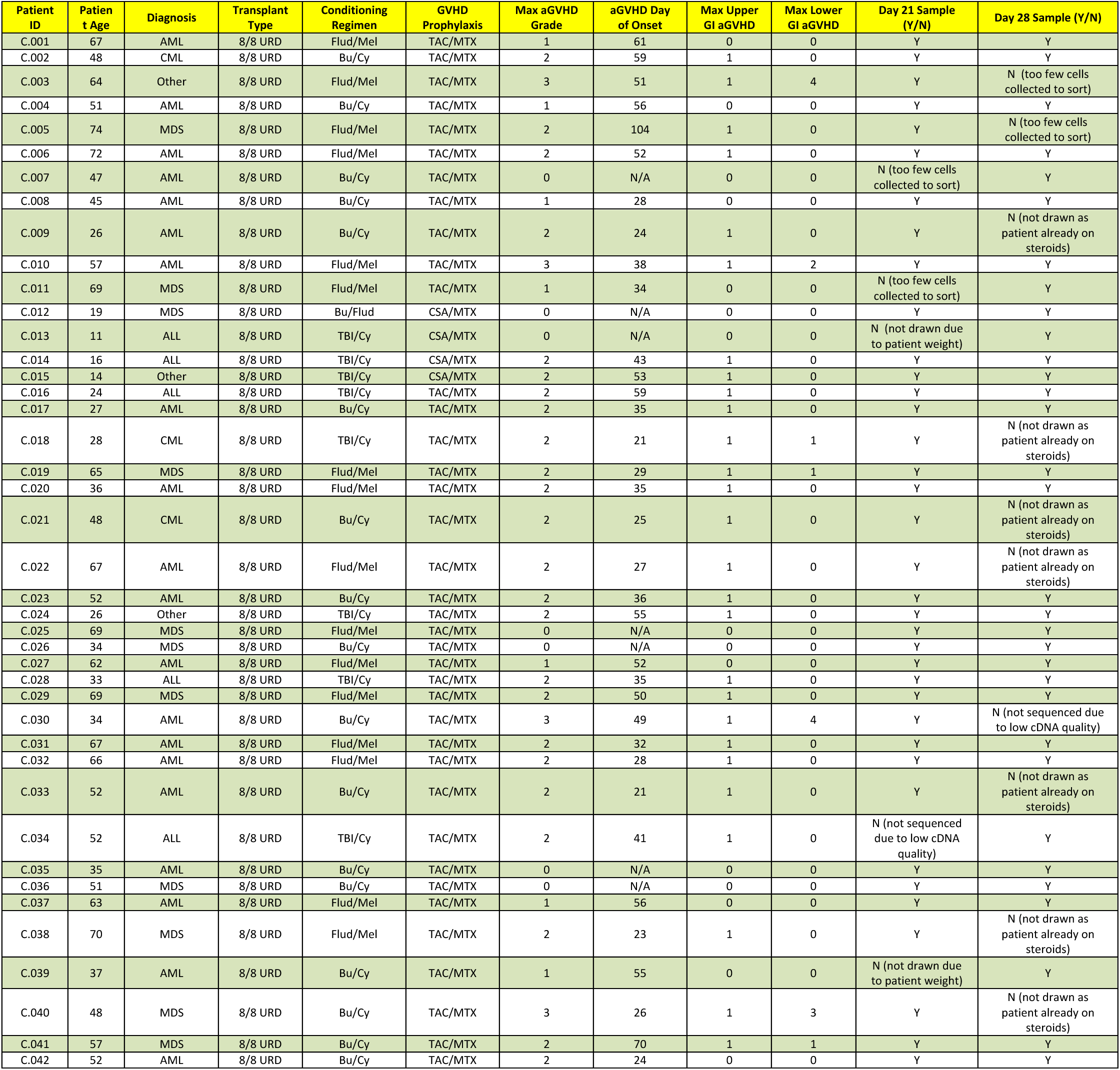
Clinical characteristics of patients from the ABA2 clinical trial (NCT01743131), included in the current study.

**Table S14.**
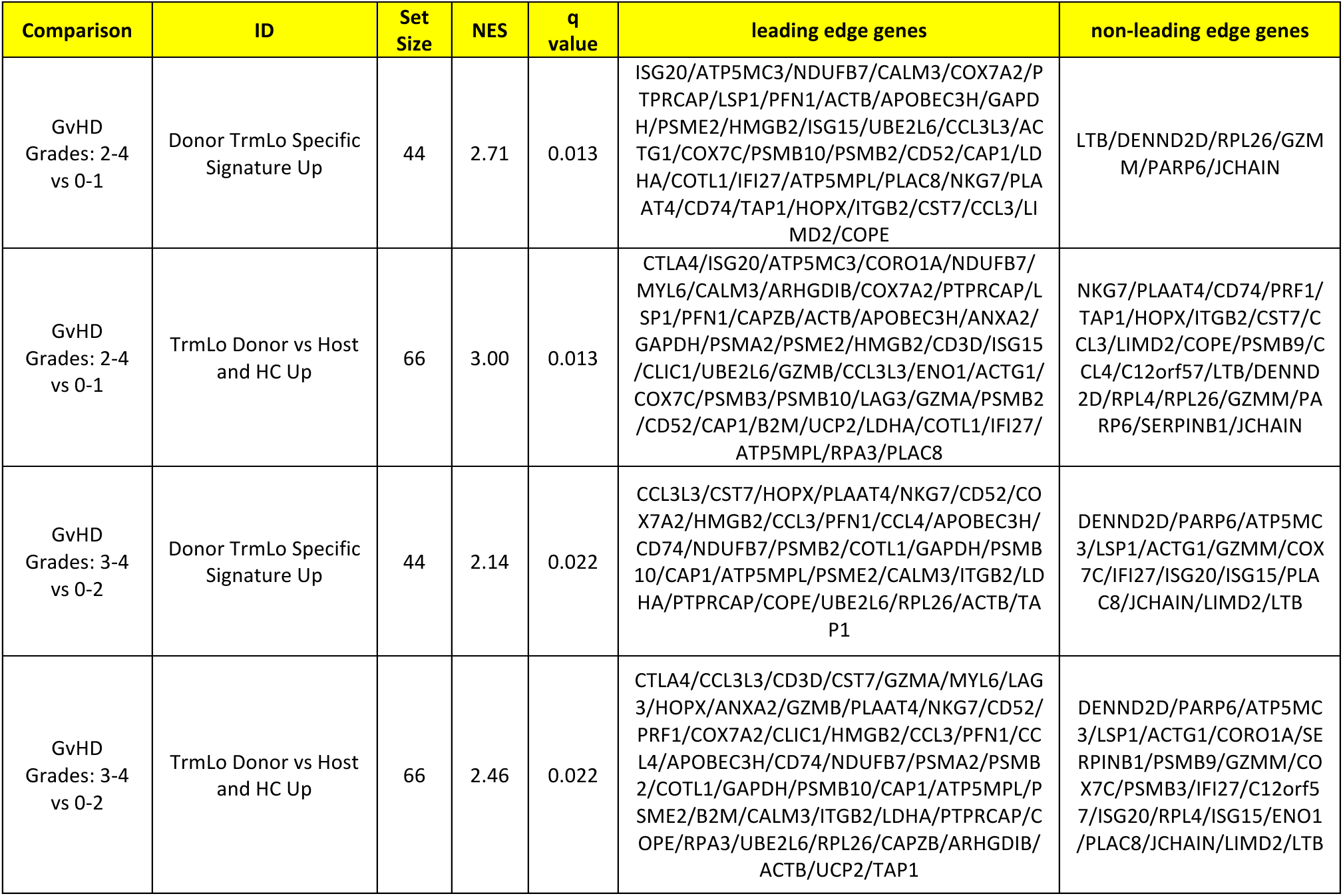
Results of GSEA analysis comparing patients from the ABA2 clinical trial (NCT01743131) with Grades 2-4 and 3-4 aGVHD versus patients with Grades 0-1 and 0-2 aGVHD, respectively.

**Table S15.**
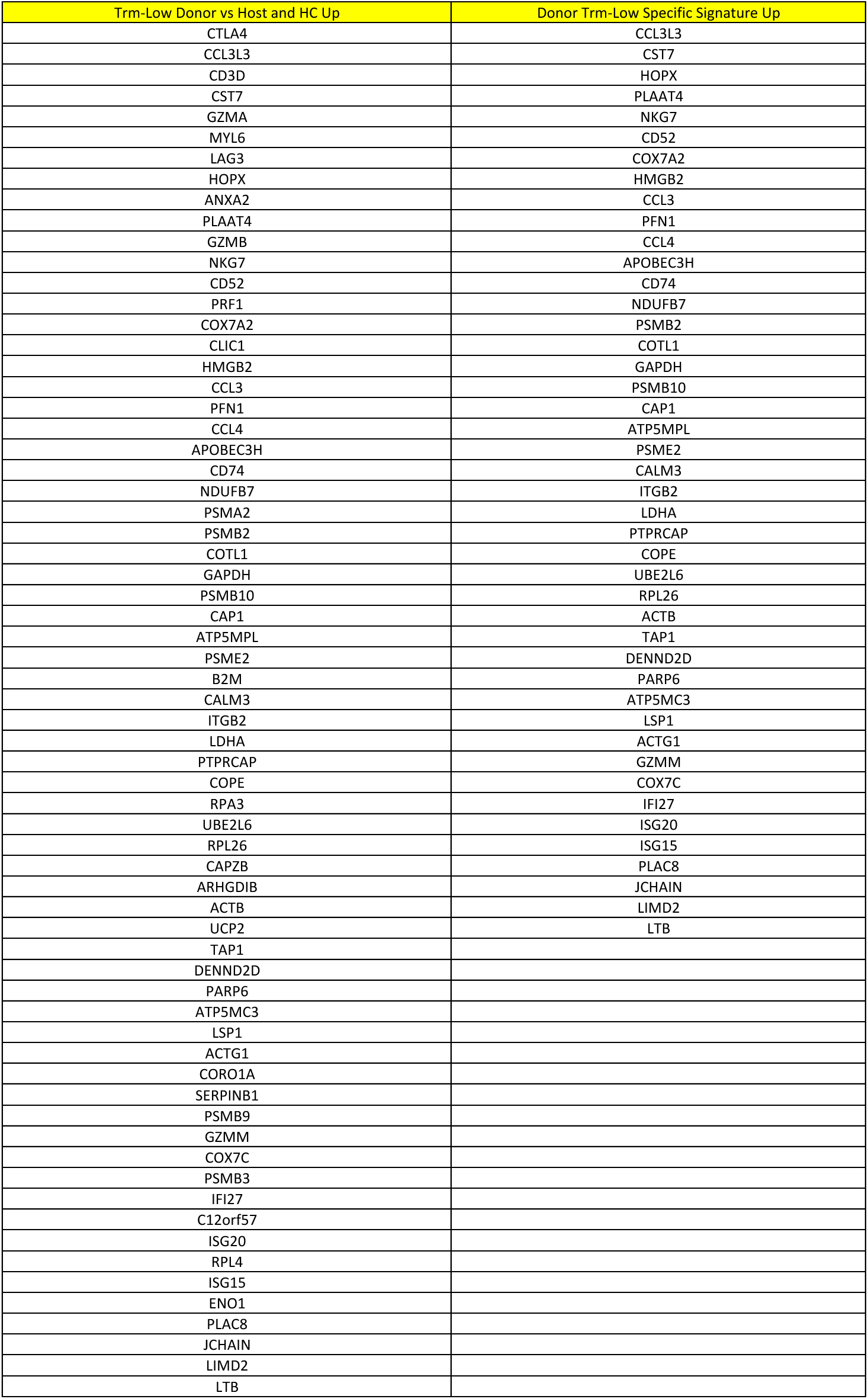
scRNASeq-derived gene sets used for GSEA analysis.

